# SCAMPI: A scalable statistical framework for genome-wide interaction testing harnessing cross-trait correlations

**DOI:** 10.1101/2024.09.10.612314

**Authors:** Shijia Bian, Zeyu Gan, Andrew J. Bass, Junji Jin, Yue Liu, Aliza P. Wingo, Thomas Wingo, David J. Cutler, Xihao Li, Michael P. Epstein

**Author notes:** Contributed equally to this work (Shijia Bian, Zeyu Gan). Contributed equally to this work (Xihao Li, Michael P. Epstein).

## Abstract

Family-based heritability estimates of complex traits are often considerably larger than their single-nucleotide polymorphism (SNP) heritability estimates. This discrepancy may be due to non-additive effects of genetic variation, including variation that interacts with other genes or environmental factors to influence the trait. Variance-based procedures provide a computationally efficient strategy to screen for SNPs with potential interaction effects without requiring the specification of the interacting variable. While valuable, such variance-based tests consider only a single trait and ignore likely pleiotropy among related traits that, if present, could improve power to detect such interaction effects. To fill this gap, we propose SCAMPI (Scalable Cauchy Aggregate test using Multiple Phenotypes to test Interactions), which screens for variants with interaction effects across multiple traits. SCAMPI is motivated by the observation that SNPs with pleiotropic interaction effects induce genotypic differences in the patterns of correlation among traits. By studying such patterns across genotype categories among multiple traits, we show that SCAMPI has improved performance over traditional univariate variance-based methods. Like those traditional variance-based tests, SCAMPI permits the screening of interaction effects without requiring the specification of the interaction variable and is further computationally scalable to biobank data. We employed SCAMPI to screen for interacting SNPs associated with four lipid-related traits in the UK Biobank and identified multiple gene regions missed by existing univariate variance-based tests. SCAMPI is implemented in software for public use.

## Introduction

Genome-wide association studies (GWAS) have successfully improved our understanding of the role of common single-nucleotide polymorphisms (SNPs) on many complex human traits and diseases. Researchers can further use SNP data from a GWAS study to estimate a trait’s narrow-sense heritability (proportion of trait variance due to additive genetic effects) using statistical techniques like GCTA and LD Score Regression (LDSC).^1,2^ Interestingly, for many complex traits, SNP-based heritability estimates are smaller than the corresponding family-based estimates of narrow-sense heritability based on kinship, and the magnitude of this discrepancy varies across traits. For instance, studies have reported SNP-based estimates of narrow-sense heritability for body mass index (BMI) to be 0.3, which is considerably less than the narrow-sense heritability estimates of 0.47-0.90 for BMI reported in twin studies.^3,4^ For Alzheimer’s Disease (AD), family-based heritability estimates of the disease range from 0.60-0.80, whereas a recent population-based AD GWAS meta-analyses estimated the narrow-sense heritability from SNP data to be between 0.06-0.41.^5–13^ Likewise, a GWAS analysis of Amyotrophic Lateral Sclerosis (ALS) estimated SNP-based heritability of approximately 0.21, which is significantly less than the estimates of 0.38-0.85 observed in twin studies.^14^

The gap between family-based estimates of narrow-sense heritability and corresponding SNP-based estimates may be due to several factors, including rare causal variation poorly tagged by common SNPs as well as shared familial environmental effects ignored in traditional family-based heritability estimation.^15,16^ Here, we focus on another possible explanation for this gap -the presence of non-additive effects (including higher-order genetic interactions) on complex traits and diseases. As noted in the Supplementary Materials (S1), we can show that higher-order interactions of a complex trait inflate narrow-sense heritability estimates more among close relatives (traditionally used for family-based estimates of heritability) than distantly related individuals (traditionally used to estimate GWAS heritability via LDSC/GCTA).^17^ Thus, higher-order interactions can explain the discrepancy between family-based and SNP-based heritability estimates observed for many complex human traits. This motivates the search for genetic variants in large-scale genetic studies that demonstrate non-additive effects, including gene-gene and gene-environment interactions.

While recent statistical methods like MonsterLM and GENIE^18–20^ enable estimation of trait variance due to interaction effects in biobank-scale data, the investigation of specific genetic variants with nonadditive effects on such traits in such datasets remains challenging despite some successes^21,22^. Exhaustive searches for pairwise epistasis still face a large computational and multiple testing burden. Gene-environment interaction analyses require fewer tests and are more computationally feasible, but measuring the most relevant environmental determinants can be difficult and is often unknown.^23–26^ To circumvent uncertainty about the right environmental factor yet still test for evidence of interaction, Paré et al. proposed an efficient variance-based method for a quantitative trait that screens for SNPs with possible interactive effects without requiring specification of the interacting factor.^27^ Recognizing that a SNP with an interaction effect on a trait induces trait variance that differs by genotype (see Supplementary Figure S1), Paré screened for SNPs with potential interaction effects by testing for equality of variances across genotype categories using Levene’s test.^28^ Researchers have successfully applied this type of variance-based approach within the UK Biobank to identify genetic variants with interaction effects on obesity phenotypes and cardiometabolic serum biomarkers.^29,30^

The variance-based test of Paré is a univariate test that considers whether a SNP has an interactive effect with a single phenotype. However, biobanks routinely collect detailed information on a large collection of related phenotypes with shared genetic effects. Many recent methods of gene mapping illustrate the appeal of leveraging the ubiquitous phenomenon of pleiotropy across related traits when present.^31–34^ Consequently, if pleiotropic genetic variants with interactive effects exist, we expect a multi-trait statistical method that leverages this information will have improved performance over existing univariate variance-based interaction procedures. Bass et al. recently showed that a SNP with an interaction effect induces not only variance but also covariance patterns between traits that differ by genotype (which we illustrate in Supplementary Figure S2).^35^ Based on this observation, the authors developed a kernel framework for interaction testing that assessed where similarity in variance/covariance patterns among a group of modeled traits correlated with genotypic similarity at a test SNP. While more powerful than standard variance-based testing, the kernel framework of Bass lacks practical features for genetic analysis such as the inability to identify the specific phenotypes (among those modeled) that demonstrate interaction effects with the test SNP. Identifying these specific phenotypes is of substantial value for further downstream analyses.

To this end, we propose here an efficient screening method SCAMPI (Scalable Cauchy Aggregate test using Multiple Phenotypes to test Interactions) for identifying potential SNPs with interaction effects using multiple phenotypes. SCAMPI fits simple regression models relating SNP genotype to (standardized) cross products of all pairwise combinations of traits under consideration and then aggregates the correlated *P*-values from these separate regression tests together into an omnibus test using the Cauchy Combination Test.^36,37^ Similar to variance-based interaction tests, SCAMPI does not require specification of the factor that interacts with the SNP of interest, thereby reducing the computational and testing burden and enabling the scaling of the method to biobank-size datasets. Moreover, SCAMPI scales to handle many related phenotypes and can identify the specific phenotype(s) that have interaction effects among those modeled. Using simulations, we show that SCAMPI can detect interactions under various scenarios and has improved performance over univariate variance-based interaction procedures. We also applied SCAMPI to lipid panel data (an indicator of risk of heart disease and stroke) in the UK Biobank (UKBB) and identified several genes with putative interaction effects that were missed by standard univariate variance-based procedures. Supplementary Table S1 compares the features of SCAMPI with those of related interaction testing approaches. For public use, SCAMPI is implemented as an R package.

## Materials and Methods

### Motivation

We first show that a SNP with a pleiotropic interaction effect yields trait correlation patterns that differ by genotype category. We could analogously show that a SNP with a pleiotropic interaction trait effect influences the covariance patterns between traits but chose to focus on correlation due to the scale-free nature of the latter measure. For subject *i*, define *G_i_* as the subject’s genotype at a test SNP and define *W_i_* as some factor (either genetic or environmental) that interacts with the SNP to influence multiple traits. Suppose subject *i* possesses two correlated traits *Y_i_*_,1_ and *Y_i_*_,2_ that are generated under the relationships:

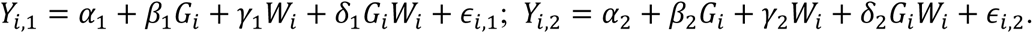

Here, *β_j_, γ_j_, δ_j_* denote the main effect of genotype, the main effect of the factor, and two-way interaction effect between genotype and factor, respectively, on trait *j* (*j* = 1, 2). We further assume each of the error terms *ϵ_i_*_,1_ and *ϵ_i_*_,2_ has a standard normal distribution *ε_i_*_,1_, *ε_i_*_,2_∼*N*(0,1). For this illustration, assume *W_i_* ∼ *N*(0,1), that *W_i_* is independent of *G_i_*, and that the two error terms are mutually independent and independent of *W_i_* and *G_i_*.

Based on the trait models listed above, Paré previously showed that that variance of *Y*_1_(*Y*_2_) differs by *G* when the genotype has an interaction effect on trait 1 (trait 2), respectively.^27^ Additionally, when pleiotropic interaction effects exist, we can show the correlation of traits 1 and 2 also differs by genotype. In Supplementary Materials S2, we derive the correlation between *Y_i_*_,1_ and *Y_i_*_,2_ conditional on genotype *G_i_* as:

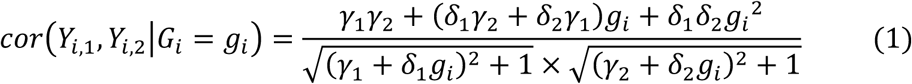

Equation (1) shows that the correlation between two traits differs by genotype when either a) the genotype interacts with the factor on both phenotypes or b) the genotype interacts with the factor on at least one of the phenotypes, provided the factor has a main effect on the other phenotype. We can also see that if the SNP has no interaction effect on either phenotype (*δ*_1_ = *δ*_2_ = 0), Equation 1 simplifies to 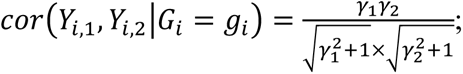 thus, the phenotypic correlation will not differ by genotype even when main effects for the factor exist (*γ*_1_ ≠ 0, *γ*_2_ ≠ 0). We observe a similar formulation when *W_i_* is binary (e.g., smoking, alcohol use) rather than continuous (see Supplementary Materials S2).

The above result suggests an efficient strategy for screening SNPs with potential interaction effects. Instead of performing traditional interaction analyses, which mandates defining potential interacting factors *W_i_*, we can instead screen for SNPs with interaction effects without having to specify *W_i_* by examining whether the correlation between traits differs across genotype categories. Such modeling provides a workaround in situations where interacting covariates are uncollected or inaccurately recorded. The screening procedure further provides an efficient alternative strategy for genome-wide epistatic analysis in that it does not require direct modeling of the interacting genetic factor, which substantially reduces the number of tests to be considered. If we are analyzing *M* SNPs, SCAMPI requires only *M* tests whereas comprehensive epistatic analysis requires 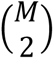 tests. Thus, when *M* = 200K (*M* = 2M), SCAMPI reduces the number of tests required by approximately 5 (6) orders of magnitude.

We implement this comparison using a general genotype model that allows each genotype category to have a separate effect relative to a baseline category. That is, for some outcome *Y\**, the coefficient estimates of *α̂*_1_, *α̂*_2_ and *α̂*_3_ in the regression model *Y*^∗^ = *α*_1_ + *α*_2_*G* + *α*_3_*G*^2^ + *ε*^∗^ can be directly mapped to coefficient estimates *β̂*_1_, *β̂*_2_ and *β̂*_3_ in ANOVA model *Y*^∗^ = *β*_1_ + *β*_2_*G*_1_ + *β*_3_*G*_2_ + *ε*, where *G*_1_ and *G*_2_ are genotype indicators for those with 1 and 2 copies of the minor allele, respectively (those with 0 copies are treated as baseline). Hence, modeling trait correlation as a function of genotype indicators provides a straightforward approach to investigate whether the trait correlation varies across genotype categories. Furthermore, given the familiarity of this general genotype model in GWAS^38–41^, we chose to use this alternative parameterization in our method moving forward.

### Notation and Trait Standardization

Assume a sample of *N* unrelated subjects that possess *J* continuous phenotypes. Let ***Y***_*j*_ = (*Y*_1,*j*_, *Y*_2, *j*_, … , *Y_N_*_,*j*_)^*T*^ A denote the *N* x 1 vector of observations for trait *j* (*j* = 1, … , *J*). Define ***G*** = (*G*_1_, *G*_2_, … , *G_N_*)*^T^* as an *N* × 1 vector of genotypes for one test SNP, where *G_i_* represents [0, 1, 2] copies of the minor allele that subject *i* possesses at the site. We use a general genotype model to compare the three genotype categories without imposing an additive relationship. Consequently, further define 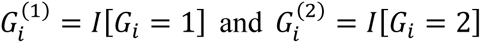 as indicator variables for genotype categories 1 and 2, respectively (we treat genotype category 0 as baseline). Finally, let ***Z*** be an *N* × *K* matrix of covariates. These covariates can be a mixture of continuous or categorical features. Common covariates to adjust for include age, biological sex, batch ID, and principal components of ancestry to deal with population stratification.

Our goal is to detect a SNP with an interaction effect that yields correlation patterns that differ by genotype. Such trait pattern differences can erroneously arise if the main effect of the genotype, as well as main and variance effects of the common covariates (such as population structure), are unaccounted for prior to analysis^42,43^. To avoid this issue, we first standardize and adjust each ***Y***_*j*_(*j* = 1, … , *J*) prior to analysis using a double generalized linear model (DGLM) that corrects for the mean effects of the test SNP and covariates, as well as the potential variance effects of covariates^44,45^.

DGLM is composed of two sub-models, where the first sub-model controls population mean, and the second sub-model controls population variance. For our work, the first sub-model adjusts ***Y***_*j*_ for the mean effects of ***G***^(1)^ & ***G***^(2)^ and covariates ***Z*** using the following framework:

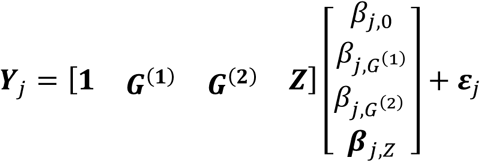

where *β*_j,0_ is the intercept associated with the *j^t^*^ℎ^ trait. *β*_*j*,*G*_(1) and *β*_*j*,*G*_(2) are the regression coefficients for ***G***^(1)^ and ***G***^(2)^ respectively, and ***β***_*j,Z*_ is a *K* × 1 vector of regression coefficients for covariates ***Z***. Finally, ***ε***_*j*_ is a *N* × 1 vector of residual errors that follow

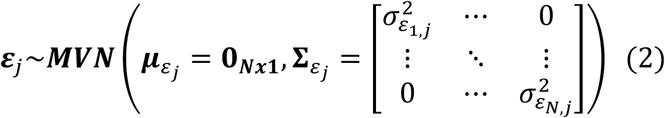

The second sub-model of the DGLM then models ***ε***_*j*_ in (2) as a function of covariates ***Z*** using the following framework using the log link function:

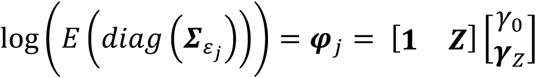

where ***φ***_*j*_ is the *N x 1* column vector representing the expected residual variance of the *j^t^*^ℎ^ observed trait. Here, *γ*_0_ is the intercept while ***γ***_Z_ represents the *K* × 1 column vector of covariate effects on the variance. The error distribution to be used in the two sub-models is Gaussian.

We fit the above DGLM using the R package “dglm”. Let ***Ỹ_*j*_*** (*j* = 1, … , *J*) denote the adjusted and standardized form for trait *j* produced from the DGLM model fit. We subsequently use ***Ỹ_*j*_*** (*j* = 1, … , *J*) to construct appropriate measures for our downstream screening analyses for interaction effects.

### Analysis Strategy

For *J* = 2 traits, we show in Supplementary Materials (S3) that we can approximate the sample Pearson correlation coefficient of traits ***Y***_1_ and ***Y***_2_ as the average of the *N* × 1 vector of cross products of the standardized traits ***Ỹ***_1_ and ***Ỹ***_2_. That is, we estimate the Pearson correlation between ***Y***_1_ and ***Y***_2_ as the sample average of

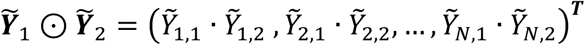

where ⊙ denotes the row-wise product operator of two vectors. Similarly, we can estimate the variance of ***Y***_1_ and ***Y***_2_ by the average of ***Ỹ***_1_ ⊙ ***Ỹ***_1_ and ***Ỹ***_2_ ⊙ ***Ỹ***_2_, respectively.

Using these estimates, we construct a screening procedure to identify a SNP with an interaction effect on trait ***Y***_1_ and/or ***Y***_2_ by assessing whether SNP genotype ***G*** is associated with either ***Ỹ***_1_ ⊙ ***Ỹ***_1_ **, *Ỹ***_2_ ⊙ ***Ỹ***_2_, or ***Ỹ***_1_ ⊙ ***Ỹ***_2_. Examination of the relationship of ***G*** with ***Ỹ***_1_ ⊙ ***Ỹ***_1_ (or ***Ỹ***_2_ ⊙ ***Ỹ***_2_) is similar to assessing whether trait variance differs by genotype (which Paré^27^ investigated using Levene’s test) while the study of ***G*** with ***Ỹ***_1_ ⊙ ***Ỹ***_2_ leverages additional information on interactions based on differences in trait correlations. To implement our procedure, we fit 3 separate ANOVA models; each model treating one of ***Ỹ***_1_ ⊙ ***Ỹ***_1_**, *Ỹ***_2_ ⊙ ***Ỹ***_2_, or ***Ỹ***_1_ ⊙ ***Ỹ***_2_ as outcome with SNP genotype ***G***^(**1**)^ and ***G***^(**2**)^ as predictors. Each ANOVA model produces a *P*-value based on a two-degree-of-freedom test. Since the resulting 3 *P*-values from these ANOVA tests are correlated, we can then combine them into an omnibus *P*-value (described in the next section) to assess whether the SNP has an interaction effect on at least one of the two traits under study.

The above example considered two traits under study. However, the strategy easily extends to the study of *J* > 2 correlated traits as well. Assuming *J* traits, we fit *J* ANOVA models that regress ***Ỹ***_*j*_ ⊙ ***Ỹ***_*j*_ on ***G***^(**1**)^ and ***G***^(**2**)^ (*j* = 1, …, *J*) and further fit 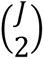 additional ANOVA models that regress ***Ỹ***_*j*_ ⊙ ***Ỹ****_l_* (*j*, *l* = 1, … , *J*; *j* ≠ *l*) on ***G***^(**1**)^ and ***G***^(**2**)^. We then can combine the 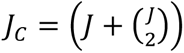 *P*-value from these tests together to assess whether the SNP has an interaction effect on at least one of the *J* traits under study.

### Cauchy Combination Test (CCT)

After obtaining the *J_C_ P*-values above, we create a final omnibus test for whether the test SNP has an interactive effect on any of the traits under consideration using the Cauchy Combination Test (CCT)^36,37^ which is a popular technique for aggregating many potentially dependent tests of high dimension together into an omnibus framework. The CCT has theoretically justified tail behavior at stringent significance levels under broad dependence structures among valid *P*-values. CCT is especially useful when an SNP signal is sparse and only affects a subset of the traits under consideration. The test statistics of CCT is a weighted sum of the Cauchy transformation of individual *P*-values in SCAMPI. Let *p_r_* denote the dependent individual *P*-value from the *r^th^* ANOVA test (*r* = 1,2, … , *J_c_*). The CCT statistic is defined as

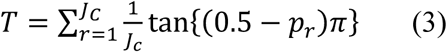

We calculate the SCAMPI omnibus *P*-value at the given genotype ***G*** using the standard Cauchy upper tail, 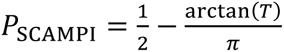 The equal weights in Equation (3) sum to one, corresponding to a Cauchy reference distribution with location zero and scale one. The CCT uses this tail approximation to combine dependent *P*-values.

### Overview of the SCAMPI Framework

Our SCAMPI framework aggregates the ANOVA tests outlined earlier with the CCT to produce an omnibus *P*-value for testing whether the SNP has an interactive effect with at least one of the traits under study. SCAMPI, which is implemented in a public R package of the same name, requires the following inputs:

a. Multiple target traits are denoted as ***Y***. Should these traits not follow a normal distribution, users can apply a rank-based Inverse Normal Transformation (INT) to normalize the traits, if desired.
b. The covariates, represented by ***Z***;
c. One test SNP, represented by ***G*** and coded as ***G***^(1)^ and ***G***^(2)^.

SCAMPI then follows the workflow depicted in Figure 1.

**Figure 1.**
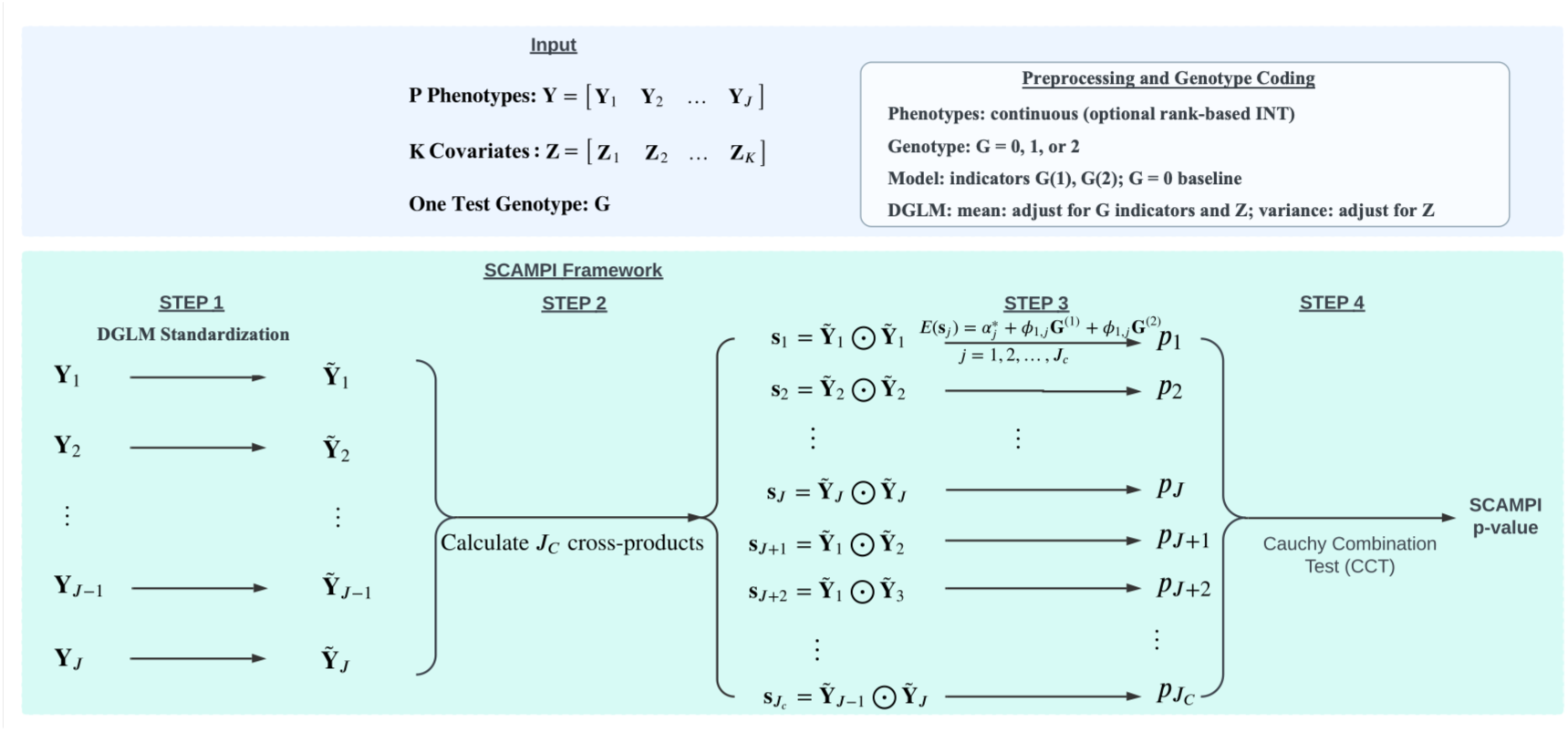
Illustration of the SCAMPI framework. Inputs consist of continuous phenotypes, common covariates, and one test genotype coded as 0, 1, or 2. Continuous phenotypes may optionally undergo rank-based inverse normal transformation, and Step 1 uses a double generalized linear model to adjust the phenotype mean for genotype indicators and covariates and the phenotype variance for covariates. The SCAMPI framework involves a four-step process that consists of 1) adjustment of phenotypes for genotype and common covariates, 2) calculation of cross products from adjusted phenotypes, 3) derivation of *P*-values from ANOVA tests of cross products on test genotype, and 4) aggregating all *P*-values using the Cauchy Combination Test (CCT) to derive the final SCAMPI *P*-values to determine overall significance.

We note the current SCAMPI implementation requires the tested variant to be represented by three discrete genotype categories. Imputed variants may therefore be analyzed using software-generated best-guess genotypes coded as 0, 1, or 2 after appropriate imputation quality and call confidence filtering. Continuous dosages and posterior genotype probability vectors are not accepted directly by the current implementation.

### Application to UK Biobank Data

We applied SCAMPI to identify SNPs with potential interaction effects on lipid measures within the UK Biobank (application ID 42223). We focused attention on four lipid-related measures: high-density lipoprotein cholesterol (HDL-C), low-density lipoprotein cholesterol (LDL-C), triglycerides (TGs), and body mass index (BMI). Both the sample and SNP QC procedures are in accordance with Marderstein et al.^46^ Similar QC procedures were also carried out in multiple studies^47^. From the cohort, we excluded individuals who either (1) had missing heterozygosity information, (2) were outliers in terms of heterozygosity or had missing genotype rates greater than 0.02, (3) had over 10 putative third-degree relatives in the kinship table, (4) were omitted from the kinship inference procedure, or (5) were either self-reported as anything other than ‘White British’ or did not show similar genetic ancestry to this group based on a principal components analysis of the genotypes. After performing this quality control, 337,422 independent subjects remained (NFemale= 181,203; NMale= 156,219). Moreover, the UKB employed two genotyping arrays. In this post-QC sample, we have the UK Biobank Axiom array (NUKBB= 300,345) and the UK BiLEVE array (NUKBL= 37,077). For the SNP QC, genotypes were discarded if they had an INFO score < 0.8, MAF < 0.05 and HWE *P*-value < 10^-10^. After SNP QC procedures, 288,910 SNPs were retained. Finally, 277,653 SNPs were included for analysis using SCAMPI after applying a 10% missing rate threshold.

We first adjusted the four lipid-related traits for common covariates (including the first six genetic principal components, biological sex, age, age squared [age²], and the type of genotyping array) before applying SCAMPI to these traits. Notably, the first six principal components effectively captured population structure at subcontinental geographic scales.^48,49^ Of the initial set of 337,422 independent subjects, 288,709 possessed complete information on all traits and common covariates and were considered moving forward. We first transformed the four traits using the inverse normal transformation (INT) to align the traits, which is a common practice to ensure the residual of traits is normally distributed in a regression model such as DGLM.^50–53^ The distribution of the four traits, both pre- and post-INT, can be found in Supplementary Figure S3 (a) - (d). Correlation between post-INT traits was 0.1246 for HDL-C and LDL-C, -0.4938 for HDL-C and TG, -0.3809 for HDL-C and BMI, 0.2797 for LDL-C and TG, 0.0394 for LDL-C and BMI, and 0.3708 for TG and BMI.

To evaluate sensitivity to phenotype scale, we repeated our analyses in the same 288,709 participants and 351,201 matched quality control variants using three phenotype definitions. The pre-INT analysis used natural-log transformed triglycerides and the original LDL cholesterol, HDL cholesterol, and BMI values. Direct INT transformed each phenotype before covariate adjustment. Indirect INT first residualized each phenotype on the common covariates and then applied the rank-based inverse normal transformation to the residuals^50^. We applied SCAMPI separately to each scale using the same genotype and covariate data. We compared genomic inflation factors, genome-wide significant SNP and LD defined locus counts, and concordance of *P*-values across scales. Genomic inflation factors were calculated as the median of the one degree of freedom chi square statistics corresponding to the SCAMPI *P*-values, divided by their theoretical median of 0.4549. In addition to studying the effect of phenotype scale, we performed additional sensitivity analyses that examined potential direct effects of BMI on the remaining lipid traits (see Supplementary Materials S4). Further, as type-I error rates of variance-based tests of a SNP can be impacted by LD with a neighboring SNP with large additive effects, we further performed conditional analyses of our SCAMPI-identified loci where we adjusted for the main effects of nearby known GWAS lipid loci (see Supplementary Materials S5).

### Simulations: Type-I Error Rate

We conducted comprehensive simulations to evaluate the type I error rate of SCAMPI under a variety of scenarios. In our first scenario, we simulated a sample size of 300,000 to reflect biobank-scale datasets. Each scenario is analyzed based on 100,000 simulations. We assumed *J* = 2,4,8 traits and simulated the trait values for the *i^t^*^ℎ^ individual based on the multivariate normal distribution illustrated below:

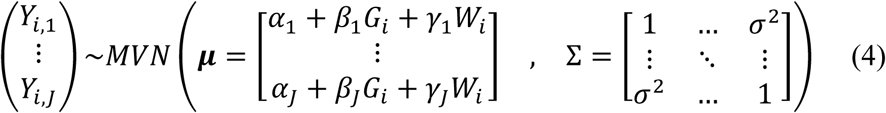

For predictors, we generated the test SNP genotype *G_i_* under Hardy-Weinberg equilibrium, assuming the SNP had a minor-allele frequency of either 0.05 or 0.25. We further generated a factor *W_i_* that followed a standard normal distribution. For the choice of parameters in the equation, we simulated the intercept *α*_*j*_ from *N*(0, 5), the genotype main effect *β*_*j*_ from *Unif*(0, 0.2), and the factor main effect *γ*_*j*_ from *Unif*(0, 0.3) (*j* = 1, … , *J*). In the covariance matrix Σ in (4), the off-diagonal covariance elements are assigned as *σ*^2^. We performed different simulations assuming *σ*^2^ = 0.01 (negligibly correlated traits), 0.25 (moderately correlated traits), and 0.5 (strongly correlated traits). For *J* = 4 traits, we conducted additional simulations where we considered a specific covariance matrix that mirrored the observed covariance structure of the lipid-related traits that we studied in the UKBB dataset (see previous section).

In addition to this initial type-I error rate simulation scenario, we conducted additional null simulations based directly on permutation of the UK Biobank sample. In each permutation, the raw phenotype data and covariates were jointly permuted across participants relative to the fixed genotype data. SCAMPI was then applied in the permuted datasets to variants with MAF of at least 0.05 and genotype missingness below 10%. We performed 200 phenotype permutations, yielding 46,429,908 valid SCAMPI *P*-values under the null hypothesis.

### Simulations: Power

For power simulations, we implemented a similar simulation design as our initial type I error simulation scenario in equation (4) but introduced additional parameters to model the effect of the interaction between SNP and the factor on the simulated traits. Specifically, we generated *J* traits based on the multivariate normal distribution as presented in Eq.

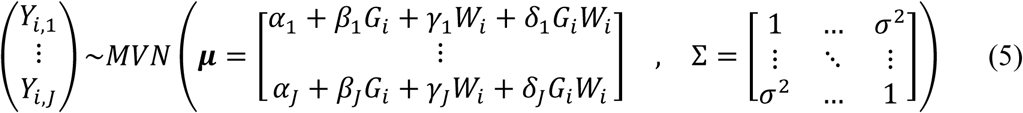

*δ*_*j*_ (*j* = 1, … , *J*) in equation (5) represents the interaction effect of the SNP and factor on trait *j*. Because *W* was generated as a standard normal variable and the residual variance of each trait was set to one, *β*_*j*_ represents the per-allele mean difference in trait *j*, *γ*_*j*_ represents the effect of a one standard deviation increase in *W* in the reference genotype group, and *δ*_*j*_ represents the per-allele change in the effect of *W*, all expressed relative to the residual standard deviation. The range of *δ*_*j*_ was selected to span weak to strong interaction effects and thereby characterize the complete power curve.

For a given simulation scenario, we vary the percentage of traits that possess such an interaction (i.e. the sparsity of the interaction signal) among the values 25%, 50%, 75%, and 100%. For those traits with an interaction effect, we vary the value of *δ*_*j*_ across a range of values from 0.01 to 0.50 to study how the power trends change as *δ*_*j*_ increases for each scenario. The settings for the number of traits, MAF, *α*_*j*_, *β*_*j*_ align with those in the Type I error simulations. However, *γ*_*j*_ is held at fixed values for all traits instead of being simulated from a distribution. Without loss of generality, this approach eliminates the potential for power fluctuations arising from the randomness in *γ*_*j*_. We simulated the results for various combinations under different parameter sets. To illustrate the overall pattern of the power simulation, we selected the simulation with *γ* = 0.05 and 0.25, and *σ*^2^ = 0.1, 0.3 and 0.5. For each simulation scenario, we assumed a sample size of 20K and generated 10K replicates for inference. To align the assessment of null calibration with the power simulation design, we also conducted an additional set of matched null simulations with a sample size of 20,000 and the same parameter settings as our power simulation with the exception that all interaction effects were set to zero. We evaluated these null simulations using 100,000 valid replicates per scenario.

We chose to benchmark SCAMPI against an enhanced multi-phenotype version of Levene’s test that was originally restricted to a single phenotype^27^. This enhanced version is termed as the multivariate Levene’s test in our context. The multivariate Levene’s test applies Levene’s test (described in Supplementary Materials S6) to each trait separately, resulting in *J P*-values. These *J P*-values are then aggregated together into an omnibus test using the CCT methodology detailed in the prior section (see Supplementary Figure S4 for an outline of the framework). While this benchmark examines how variances vary by genotype across different traits, it does not consider differences in correlation patterns among traits that SCAMPI integrates within its framework.

### Sensitivity Analyses

In addition to our initial type-I error and power simulations, we conducted additional simulations to assess the sensitivity of SCAMPI to various modeling assumptions. We performed simulations to investigate the finite-sample behavior of SCAMPI and specifically the impact of genotype-group size and sparsity on type-I error and power (see Supplementary Materials S7). We also performed simulations investigating robustness of SCAMPI to non-normality of outcomes (see Supplementary Materials S8), the impact on SCAMPI inference when one outcome has a direct effect on the remaining outcomes (see Supplementary Materials S9), and impact on SCAMPI when outcomes demonstrate high residual correlation and have heterogenous environmental effects (see Supplementary Materials S10).

## Results

### Simulation Studies

Table 1 provides empirical type I error rates for SCAMPI summarized at a nominal rate *α* of 10^−2^ and 10^−3^ across varying numbers of phenotypes, MAF and Σ when *γ* is simulated from *Unif*(0, 0.3). As described in Supplementary Materials S11, we focused primarily on studying the empirical type I error rate at 10^−3^ based on the number of simulations performed and observed that SCAMPI was well calibrated under the evaluated biobank scale common variant settings at this threshold. To examine whether SCAMPI was well calibrated at more stringent thresholds, we first analyzed more than 1 million null tests generated from the original UK Biobank permutation analysis. The empirical type I error rates were 1.08 × 10^-2^, 1.06 × 10^-3^, and 9.81 × 10^-5^ at nominal significance levels of 10^-2^, 10^-3^, and 10^-4^ respectively. We subsequently expanded this analysis to 200 phenotype permutations, yielding 46,429,908 valid SCAMPI tests under the null hypothesis. The QQ plot from the expanded analysis showed no marked systematic inflation across the empirically observable range, including the tail approaching the genome wide significance region (Supplementary Figure S5). The precision of this null assessment at the genome-wide thresholds is described in Supplementary Section S11^36,54–56^.

**Table 1.** Nominal rate of empirical type I error rates for SCAMPI. The empirical type I error rates for SCAMPI at nominal rates *α* of 10^-2^ and 10^-3^ in 100,000 simulations with 300,000 observations. The result is presented across a range of conditions including varying Minor Allele Frequencies (MAF), numbers of phenotypes (*J*), and covariance of the phenotypes Σ when *γ* is simulated from a uniform distribution between 0 and 0.3. The values presented in the “≤ 0.01” and “≤ 0.001” columns are the empirical rejection rates at the corresponding nominal significance levels.

| **MAF** | ***Σ*** | ***J*** | **≤ 0.01** | **≤ 0.001** |
|---|---|---|---|---|
| **0.05** | **0.01** | **2** | 9.56 × 10<sup>−3</sup> | 1.01 × 10<sup>−3</sup> |
|  |  | **4** | 1.00 × 10<sup>−2</sup> | 1.18 × 10<sup>−3</sup> |
|  |  | **8** | 1.05 × 10<sup>−2</sup> | 1.13 × 10<sup>−3</sup> |
| **0.05** | **0.25** | **2** | 9.96 × 10<sup>−3</sup> | 1.09 × 10<sup>−3</sup> |
|  |  | **4** | 1.09 × 10<sup>−2</sup> | 9.30 × 10<sup>−4</sup> |
|  |  | **8** | 1.12 × 10<sup>−2</sup> | 1.01 × 10<sup>−3</sup> |
| **0.05** | **0.5** | **2** | 1.05 × 10<sup>−2</sup> | 1.16 × 10<sup>−3</sup> |
|  |  | **4** | 1.16 × 10<sup>−2</sup> | 1.02 × 10<sup>−3</sup> |
|  |  | **8** | 1.26 × 10<sup>−2</sup> | 1.23 × 10<sup>−3</sup> |
| **0.25** | **0.01** | **2** | 9.51 × 10<sup>−3</sup> | 1.17 × 10<sup>−3</sup> |
|  |  | **4** | 1.03 × 10<sup>−2</sup> | 1.13 × 10<sup>−3</sup> |
|  |  | **8** | 1.00 × 10<sup>−2</sup> | 1.14 × 10<sup>−3</sup> |
| **0.25** | **0.25** | **2** | 1.01 × 10<sup>−2</sup> | 1.12 × 10<sup>−3</sup> |
|  |  | **4** | 1.09 × 10<sup>−2</sup> | 1.06 × 10<sup>−3</sup> |
|  |  | **8** | 1.09 × 10<sup>−2</sup> | 9.40 × 10<sup>−4</sup> |
| **0.25** | **0.5** | **2** | 1.03 × 10<sup>−2</sup> | 9.80 × 10<sup>−4</sup> |
|  |  | **4** | 1.09 × 10<sup>−2</sup> | 1.03 × 10<sup>−3</sup> |
|  |  | **8** | 1.28 × 10<sup>−2</sup> | 1.09 × 10<sup>−3</sup> |

We next evaluated the finite-sample behavior of SCAMPI under the null as a function of the minimum genotype group count (see Supplementary Materials S7). At nominal *α* = 0.01, the empirical rejection rates at target minimum counts of 100, 150, and 200 were 0.01130, 0.01051, and 0.01057 for *J* = 2 traits; 0.01296, 0.01175, and 0.01184 for *J* = 4; and 0.01396, 0.01306, and 0.01270 for *J* = 8, respectively. Thus, increasing the minimum count generally reduced type I error inflation, although it did not guarantee exact calibration across all trait dimensions. When *J* = 4, *ρ* = 0.25, and the target minimum count was fixed at 150, the rejection rates were similar for *N* = 20,000, 100,000, *and* 300,000 (0.01151, 0.01175, and 0.01155, respectively), indicating that the minimum genotype group count was more directly informative than total sample size alone (Supplementary Table S2 and Supplementary Figure S6A).

We next compared the power of SCAMPI and multivariate Levene testing. Figure 2 shows representative results at *P* < 6.25 × 10^-8^ for four traits, 20,000 individuals, MAF = 0.05, environmental main effects of 0.05, and the UK Biobank matched correlation matrix. Power increased with interaction strength and the number of traits carrying an interaction, and SCAMPI outperformed multivariate Levene testing across the displayed settings (Supplementary Figure S7). These comparisons use the nominal threshold; the expected minimum genotype group count is approximately 50, and the matched null results in Supplementary Table S3 quantify the associated finite sample inflation.

**Figure 2.**
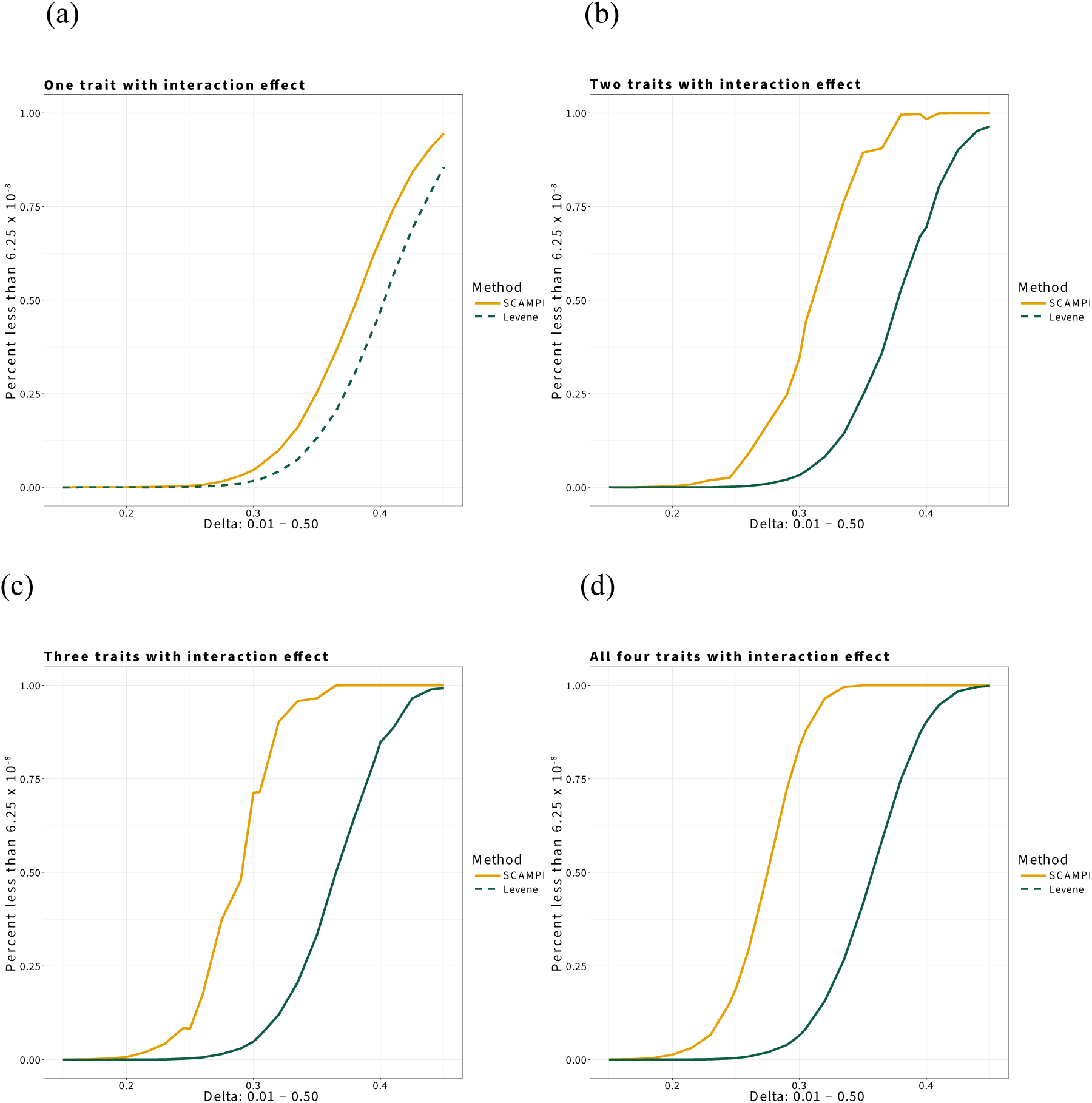
UKBB-matched power comparison of SCAMPI and the multivariate Levene’s test for four traits. The x axis represents the genotype by factor interaction coefficient δ, varied from 0.15 to 0.45. The y axis represents the proportion of 10,000 alternative replicates with an omnibus *P*-value below the genome wide significance threshold of 6.25 × 10^-8^. Each simulation included 20,000 individuals, a test SNP with a minor allele frequency of 0.05, and a factor main effect of 0.05. The trait correlation matrix was matched to that observed among HDL cholesterol, LDL cholesterol, triglycerides, and BMI in the UK Biobank analysis. Panels (a) -(d) assume interaction effects on one, two, three, and all four traits respectively. The yellow solid line represents SCAMPI, and the dashed green line represents the multivariate Levene’s test. Under Hardy-Weinberg equilibrium, this design has an expected minimum genotype group count of approximately 50. The displayed values are therefore nominal alternative rejection probabilities and should be interpreted together with the matched null calibration results in Supplementary Table S3.

In addition to the power simulations inspired by the UKBB, Supplementary Figure S8 provides power results for SCAMPI and multivariate Levene’s test under a broader range of models that vary the number of traits considered, the sparsity of the interaction effect, the correlation among traits, and the main effect of the variable interacting with genotype. Overall, we find the power of SCAMPI increases with a decrease in the sparsity of the interaction effect, a decrease in the trait correlation, and an increase in the effect size of the interaction variable. Assuming these three inputs are fixed, we find that the power of SCAMPI increases as the number of traits modeled increases. Regarding the power comparisons between SCAMPI and the Multivariate Levene’s test under this broader range of models, Supplementary Figure S8 also reaffirms the trends observed in our UKBB-inspired power simulations. Across the spectrum of scenarios tested, SCAMPI consistently exhibited superior performance when compared to the Multivariate Levene’s test, largely because the former method accounts for correlation among traits that the latter method ignores.

Given the sample size of our power calculations (20K) was smaller than that of our earlier type-I error simulations, we conducted additional type-I error rate simulations assuming this smaller sample size under the same parameter settings to ensure our power results were well calibrated. When the minor allele frequency was 0.25, empirical type I error rates ranged from 0.00966 to 0.01319 at the nominal level of 0.01 and from 0.00082 to 0.00142 at the nominal level of 0.001. When the minor allele frequency was 0.05, the corresponding ranges were 0.01134 t 0.01797 and 0.00159 to 0.00350. The average minimum genotype group count was approximately 50 for a minor allele frequency of 0.05 and approximately 1,250 for a minor allele frequency of 0.25. Thus, the deviations observed in the lower frequency settings were consistent with the small genotype group behavior identified in our minimum genotype group count simulations. Complete results and exact 95% binomial confidence intervals are provided in Supplementary Table S3. We also investigated the impact of genotype group sparsity on the empirical power of SCAMPI (Supplementary Materials S7, Supplementary Table S4, and Supplementary Figure S6B).

### Sensitivity Analyses

We additionally studied the performance of SCAMPI when trait outcomes were non-normal (following either a standardized t distribution with three degrees of freedom [t3] or a standardized log-normal distribution) and investigated whether an inverse normality transformation (INT) of such outcomes impacted inference (see Supplementary Materials S8). We applied SCAMPI to the pre-transformed outcome data (pre-INT), transformed data before covariate adjustment (Direct INT), and transformed residuals after covariate adjustment on the original scale (Indirect INT). For a strict null model with no genotypic main effects, we observed that use of pre-transformed outcomes yields type-I error inflation with, at nominal α = 0.01, pre-INT rejection rates of 0.05828 under t3 distribution and 0.05945 under log-normal distribution. However, Direct INT reduced these rejection rates to 0.01299 and 0.01193, and indirect INT reduced such rates to 0.01174 and 0.01156. For null interaction simulations involving genotype main effects, we observed rejection rates for the t3 distribution of 0.05848, 0.01281, 0.01125 for the pre-INT, direct INT, and indirect INT analyses; suggesting the normality transformation is effective in this context. However, under the log-normal model containing genotype main effects but no explicit interaction, the rejection rates were 0.05783, 0.07128, and 0.48001 for the pre-INT, direct INT, and indirect INT analyses. These findings show that transformation can correct severe non-normal behavior in some settings but can also induce genotype-dependent distributional differences in others. Complete results at α = 0.05, 0.01, and 0.001 are reported in Supplementary Table S5.

We next investigated the impact of direct and mediated effects of one outcome on the remaining outcomes on inference using SCAMPI. We performed simulations (Supplementary Materials S9) mimicking our UKBB analyses (Supplementary Materials S4) that studied direct and BMI-mediated interaction mechanisms on the remaining lipid traits we considered in UKBB. For two null simulations (with and without a causal path of BMI to the lipid traits), the type-I error was close to the nominal levels for SCAMPI models that 1) jointly modeled the lipid traits and BMI as outcomes, 2) considered the lipid traits as only outcomes and ignored BMI, and 3) considered the lipid traits as only outcomes but adjusted for linear and quadratic effects of BMI. For a power simulation when interaction effect acted only on BMI and was transmitted to lipid traits, SCAMPI had excellent power when BMI was included as an outcome, negligible power when BMI was ignored, and no power when BMI was treated as a covariate. The contrasting patterns demonstrate that exclusion or adjustment of BMI can distinguish direct lipid effects from signals transmitted primarily through BMI (Supplementary Table S6 and Supplementary Figure S9).

We also studied the performance of SCAMPI when trait outcomes were tightly correlated (Supplementary Materials S10) and observed that the procedure was properly calibrated under the null. Under alternative models, the power of SCAMPI decreased as the correlation among traits increased but that the procedure still retained an advantage over Multivariate Levene’s test (Supplementary Table S7 and Supplementary Figure S10). We also studied the impact of heterogeneous environmental main effects on the traits modeled within SCAMPI and found this phenomenon had little influence on power (Supplementary Table S8 and Supplementary Figure S11).

### Application to UKBB

Figure 3 provides the Manhattan plot of SCAMPI results for detecting interaction effects on four lipid-related traits. SCAMPI identified 210 SNPs across 68 genes and intergenic regions at a study-wide significance level (*α* = 1.67 × 10^−7^, i.e., multiple comparison correction for 300,000 SNPs). Table 2 highlights the SNPs with the smallest SCAMPI *P*-value on each chromosome from the 210 SNPs. A comprehensive list of the 210 SNPs is available in Supplementary Table S9. The observed-data QQ plot shows an upward deviation beginning near *P* = 10^-3^ and a pronounced extreme tail (Supplementary Figure S12). Null calibration was evaluated separately using 200 phenotype permutations, which showed no marked systematic inflation across the empirically observable range (Supplementary Figure S5). SCAMPI is an omnibus test that, by aggregating *P*-value (outputs of Step 3 in Figure 1) from association tests of trait correlation, pinpoints the specific traits that influence the overall signal. Thus, for every lead SNP in Table 2, we examined the *P*-values linked to each trait variance and cross-trait correlation at a genome-wide significance threshold of 1.67 × 10^−7^. Significant variance and correlation terms among traits are noted in the “Significant Variance/Correlation Components” column of the Table. For example, rs7528419 at the CELSR2–-PSRC1–-SORT1 locus is significantly associated with the correlation of triglycerides and LDL, as well as the variance of LDL alone, suggesting the SNP may have an interaction effect with other genetic or environmental factors on these two specific traits that merit further investigation.

**Figure 3.**
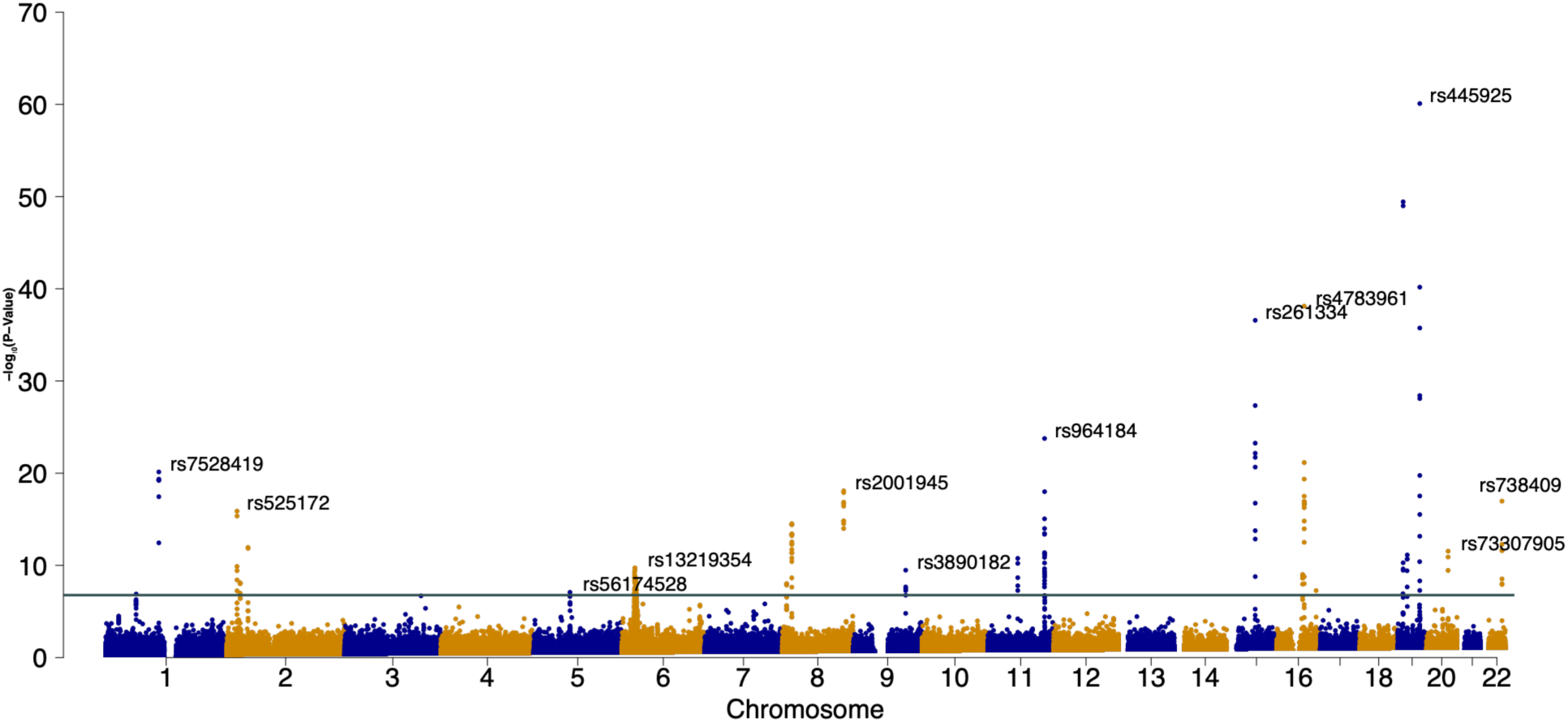
Genome-wide results on lipid traits in UKBB using SCAMPI. The x axis shows genomic position ordered by chromosome, and the y axis shows −log10(*P*) for the omnibus SCAMPI test of HDL cholesterol, LDL cholesterol, triglycerides, and BMI. After variant quality control, 277,653 SNPs were analyzed. The green horizontal line marks the genome wide significance threshold of *P*-value = 1.67 × 10⁻⁷. SCAMPI identified 210 significant SNPs across 68 annotated genes and intergenic regions. The most significant SNP on each chromosome is labeled; rs445925 at the APOC1 locus produced the smallest SCAMPI *P*-value.

**Table 2.** The lead SNPs, identified by SCAMPI within each chromosome, implies interaction effects for the four lipid traits in UKBB. SCAMPI identified 12 lead SNPs from 12 chromosomes. The position of the SNPs is based on the Genome Reference Consortium Human Build 37 (GRCh37). The “Gene” column provides a positional gene or locus annotation. Column “SCAMPI P-value” shows the SCAMPI *P*-value. Column “Significant Variance/Correlation Components” indicates the variance or the correlation components of the four lipids that are significantly associated with the corresponding SNP at the pre-specified study-wide significance (*α* = 1.67 × 10^−7^). The column “Previously Reported GWAS-associated Traits” lists traits implicated by the significant SCAMPI variance or correlation components for which the corresponding variant has a previously reported marginal association in the GWAS Catalog or UK Biobank datasets available through the Open Targets Platform. These associations provide biological context but should not be interpreted as independent replication of the interaction signals detected by SCAMPI. Gene labels are GRCh37 positional annotations and do not imply causal gene assignments. For intergenic variants, a nearby gene is shown in parentheses when listed; otherwise, the entry is labeled “Intergenic”. The chromosome 1 variant rs7528419 is annotated to CELSR2 but lies within the broader CELSR2-PSRC1-SORT1 lipid locus.

| Chr | Pos | Alt | Ref | RS # | Gene/Locus Annotation | SCAMPI *P*-value | Significant Variance/Correlation Components | Previously Reported GWAS-associated Traits |
|---|---|---|---|---|---|---|---|---|
| **1** | **109817192** | **G** | **A** | rs7528419 | *CELSR2* | 7.33E-21 | Corr(TRIG, LDL), Var(LDL) | TRIG, LDL |
| **2** | **21382976** | **G** | **T** | rs525172 | Intergenic | 1.33E-16 | Corr(TRIG, HDL), Var(LDL) | TRIG, LDL |
| **5** | **74400516** | **C** | **G** | rs56174528 | *ANKRD31* | 8.26E-08 | Var(LDL) | LDL |
| **6** | **27185664** | **C** | **T** | rs13219354 | *PRSS16* | 1.83E-10 | Var(BMI) | BMI |
| **8** | **126477978** | **C** | **G** | rs2001945 | (*TRIB1*) | 8.34E-19 | Var(LDL) | LDL |
| **9** | **107647655** | **A** | **G** | rs3890182 | *ABCA1* | 3.34E-10 | Var(HDL) | HDL |
| **11** | **116648917** | **C** | **G** | rs964184 | *ZPR1* | 1.73E-24 | Corr(TRIG, LDL), Corr(TRIG, HDL), Corr(LDL, HDL), Corr(LDL, BMI), Var(Trig), Var(LDL) | TRIG, LDL, HDL |
| **15** | **58726744** | **C** | **G** | rs261334 | *LIPC;*<br>*LIPC-AS1* | 2.63E-37 | Corr(HDL, BMI), Var(TRIG) | TRIG, HDL |
| **16** | **56994894** | **A** | **G** | rs4783961 | *CETP* | 7.61E-39 | Var(HDL) | HDL |
| **19** | **45415640** | **A** | **G** | rs445925 | *APOC1* | 8.10E-61 | Corr(TRIG, LDL), Corr(TRIG, HDL), Corr(LDL, HDL), Corr(LDL, BMI), Var(Trig), Var(LDL), Var(HDL) | TRIG, LDL, HDL |
| **20** | **44545773** | **C** | **A** | rs73307905 | (*PLTP*) | 2.90E-12 | Var(HDL) | HDL |
| **22** | **44324727** | **G** | **C** | rs738409 | *PNPLA3* | 1.07E-17 | Corr(TRIG, BMI), Var(TRIG) | BMI |

We next explored the impact of phenotypic scale on our SCAMPI results. The matched phenotype-scale comparison showed that direct INT (applying inverse normal transformation before covariate adjustment) and indirect INT (applying inverse normal transformation to residuals after covariate adjustment) produced highly concordant results, whereas the pre-INT analysis differed more substantially. Across 351,201 variants, the Pearson correlation of −log10(*P*) was 0.993 and the Spearman correlation of *P*-values was 0.978 between direct INT and indirect INT. At *P* = 1.67 × 10⁻⁷, the pre-INT, direct INT, and indirect INT analyses identified 496, 257, and 242 SNPs, corresponding to 52, 25, and 24 LD defined loci, respectively. At *P* = 6.25 × 10⁻⁸, the corresponding counts were 441, 231, and 214 SNPs and 47, 23, and 21 loci. The genomic inflation factors were 1.009, 0.940, and 0.940. Thus, the two INT procedures yielded similar inference, but analysis on the pre-INT scale produced substantially more significant signals (Supplementary Table S10 and Supplementary Figure S13).

We also performed additional sensitivity analyses that examined potential direct effects of BMI on the remaining lipid traits (see Supplementary Materials S4). In addition to our original model M0 that included BMI, triglycerides, LDL cholesterol, and HDL cholesterol, we also considered the model M1 that included the three lipid traits without BMI and the model M2 that included the same three lipid traits and adjusted for raw standardized BMI and BMI squared through the SCAMPI covariate matrix. The BMI sensitivity analysis showed substantial agreement across the three models. At 1.67 × 10^-7^, M0, M1, and M2 identified 257, 183, and 168 significant variants, corresponding to 25, 23, and 20 LD defined loci respectively; 18 loci were shared across all three models. At 6.25 × 10^-8^, the corresponding counts were 231, 168, and 154 variants and 23, 21, and 18 loci, with 15 loci shared across all three models. The genomic inflation factors were 0.940, 0.958, and 0.965 for M0, M1, and M2 respectively (Supplementary Table S11 and Supplementary Figure S14). These findings indicate that most loci were not dependent on including BMI as an outcome, although attenuation of a subset of signals suggests that BMI related pathways contributed to some associations.

### Investigation of SCAMPI Signals

We also cross-referenced the lead variants in Table 2 with previously reported trait associations from the GWAS Catalog and UK Biobank datasets available through the Open Targets Platform (version 22.10).^57^ These marginal genetic associations provide additional biological context for the loci identified by SCAMPI. For example, rs738409 in PNPLA3, which SCAMPI identified through the triglyceride-BMI correlation and triglyceride variance components, has previously been associated with BMI. The relevant previously reported trait associations are summarized in the “Previously reported GWAS-associated traits” column of Table 2. Beyond the lead SNPs, Supplementary Table S12 provides the individual variance and correlation component *P*-values for all 210 significant SNPs.

Genotype-stratified diagnostics for the five leading variants showed that all genotype groups were sufficiently large for descriptive comparison. Across the five variants, the smallest genotype group contained 3,468 participants, and genotype missingness ranged from 0.075% to 0.755%. All five variants remained significant on the pre-INT, direct INT, and indirect INT scales, although the magnitude of the *P*-values varied substantially by scale. Phenotype distributions and pairwise correlations also differed across genotype categories and analysis scales (Supplementary Table S13 and Supplementary Figures S15-S16). These plots show the genotype specific variance and covariance patterns underlying the leading SCAMPI signals.

Overall, SCAMPI identified several established lipid- and BMI-related genes that also demonstrate potential interaction effects. For example, APOC1, which contained the smallest SCAMPI *P*-value (*p* = 8.1 × 10^−61^), has pleiotropic effects on lipid metabolism, influencing various processes through its actions on lipoprotein receptors and enzyme activity modulation. By controlling the lipids plasma level, the influence of APOC1 spans several disease areas, including cardiovascular physiology, inflammation, immunity, sepsis, diabetes, cancer, viral infectivity, and cognition.^58^ CETP contained a SNP with a potential interaction effect on HDL cholesterol (*p* = 7.61 × 10^−39^). CETP influences HDL and LDL metabolism, and clinical trials of CETP inhibition have yielded heterogeneous cardiovascular outcomes,^59^ ranging from harm with torcetrapib to a modest reduction in major coronary events with anacetrapib.^60–62^

Another top gene identified by SCAMPI was LIPC. Evidence suggests the LIPC promoter polymorphism (T-514C) affects the activity of hepatic lipase (HL) and, in concert with other factors, modifies the therapeutic response in coronary artery disease (CAD) patients, with those having the CC genotype benefiting the most from intensive lipid-lowering treatments due to their predisposition to high HL activity and smaller, denser LDL particles.^63^ The chromosome 1 lead variant rs7528419 is positionally annotated to CELSR2 but lies within the established CELSR2-PSRC1-SORT1 lipid locus. We describe this association as a signal at the CELSR2-PSRC1-SORT1 locus.^64^

Interestingly, SCAMPI identified several SNPs (shown in Supplementary Table S14) exclusively through the correlation among traits (such that they were not detected by the multivariate Levene’s test that only considered variance terms). Noteworthy among these are rs2228603 (*NCAN*), rs58542926 (*TM6SF2*), and rs10415849 (*GATAD2A*). For each of these three SNPs, SCAMPI detected a significant effect exclusively via the correlation of BMI and triglycerides (each *p* <10^−8^); the SNP was not significantly associated with the variance of either trait and, as such, was not picked up by Levene’s test. These three variants are located within the NCAN-TM6SF2-CILP2 region on chromosome 19, which has previously demonstrated genome-wide significant marginal associations with triglycerides and LDL cholesterol in large lipid GWAS.^65–67^ A similar pattern was observed for three variants in NECTIN2: each was associated with the LDL-HDL correlation (each *p* < 10^−8^) but not with the variance of either trait. These variants are located within the established APOE-NECTIN2 lipid locus, which has previously been associated with LDL cholesterol, total cholesterol, and HDL cholesterol. These previous GWAS findings provide locus level context for the SCAMPI associations.

In addition to the information from the Open Targets platform, we also want to highlight that the SNPs identified by SCAMPI have been implicated in other studies of lipid traits and BMI. For example, numerous studies suggest that rs2228603(*NCAN*) and rs58542926(*TM6SF2*) are genetic variants associated with an increased likelihood of liver inflammation and fibrosis that is closely associated with weight change, indeed impacting BMI.^68–70^ rs10415849 (*GATAD2A*) is significantly associated with *α*-Tocopherol (one type of vitamin E), which interacts with biological sex to modify BMI.^71^ Recent G×E studies provide exact-variant support for several SCAMPI signals (Supplementary Table S15). The strongest examples include rs58542926 at *TM6SF2* and rs769449 at *APOE* in a large cross-population biobank analysis,^72^ rs4420638 in the APOC1 region in a CHARGE and UK Biobank analysis,^73^ and rs964184 at ZPR1/APOA5 in a randomized long-term dietary intervention.^74^ Additional exact-variant evidence was observed for loci including CELSR2, CMIP, PLTP, GCKR, TRIB1, and PNPLA3, although some of these results lacked stringent external replication or involved phenotypes adjacent to the serum lipid traits analyzed here. These overlaps support the biological plausibility of the SCAMPI findings, but they do not constitute direct replication because SCAMPI detects genotype-dependent multivariate covariance without observing the interacting exposure, whereas conventional G×E studies explicitly model a measured environmental or contextual factor.

For our APOC1 SCAMPI result, we performed a secondary conditional analysis adjusting for an approximation of the canonical APOE genotype structure represented by five indicator variables derived from the joint unphased genotypes at rs429358 and rs7412 (see Supplementary Materials S5); we adjusted for the canonical genotype structure in the DGLM mean model (see Supplementary Materials S5 and Supplementary Table S16) to assess whether additive effects of APOE on our outcomes could explain the nearby APOC1 interaction signal. Reanalyzing 47 variants in APOC1 using this APOE conditional model, we show in Supplementary Figure S17 that the lead variant rs445925 remained significant after adjustment (*P* = 9.41 × 10^-39^). Moreover, of the additional 14 APOC1 variants that were significant in our original SCAMPI analyses, 10 and 9 of these variants remained significant at 1.67 × 10^-7^ and 6.25 × 10^-8^, respectively, in these APOE-adjusted SCAMPI analyses. The regional SCAMPI signal for APOC therefore persisted after adjustment for nearby APOE genotype effects on phenotype means.

We next assessed whether our other SCAMPI signals outlined in Table 2 could be explained by mean effects of nearby lipid-associated variants in LD (see Supplementary Materials S5). We identified such nearby lipid-associated variants based on being within 500 kb of our SCAMPI signal, being significantly associated with a lipid trait (*P* < 5 × 10^-8^) in the Global Lipids Genetic Consortium GWAS^67^, and being directly observed in the UKBB sample. The SCAMPI test variant itself was excluded, and redundant conditioning variants were removed at r² ≥ 0.80 using LD estimated in the analysis sample. For each SCAMPI signal, we included the selected variants as additive covariates in the DGLM mean model. After conditioning on the established lipid-associated variants, we observed that all SCAMPI variants noted in Table 2 remained significant with adjusted *P*-values ranging from 6.58 × 10^-38^ to 1.49 × 10^-7^ (Supplementary Table S17). Moreover, of the original set of 210 SNPs identified by SCAMPI, 189 (90.0%) remained significant after adjustment for the mean effects of nearby lipid associated variants at *P* < 1.67 × 10^-7^, and 168 (80.0%) met the more stringent threshold of *P* < 6.25 × 10^-8^ (Supplementary Table S9). These conditional analyses help separate SNPs with potential interaction effects on complex traits from those that are spurious and arising instead from a nearby SNP with a strong main additive effect on the outcomes under consideration; such information can guide the selection of variants for subsequent studies of specific interacting factors.

### Computational Performance

We benchmarked the computational performance of SCAMPI across varying sample sizes and numbers of traits for analyzing a single genotype using the High-Performance Computing (HPC) cluster hosted by Emory University Rollins School of Public Health (RSPH). SCAMPI was computationally tractable across the benchmark settings evaluated (Figure 4). Runtime per variant increased approximately linearly with sample size and quadratically with the number of analyzed traits. For 300,000 individuals and four traits, the mean runtime was 20.17 seconds per variant on the RSPH high-performance computing system. Because genome-wide wall-clock time depends on the parallel resources available, complete hardware specifications, benchmark settings, total computational effort, and parallel-runtime projections are provided in Supplementary Materials S12 and Supplementary Figures S18-S19.

**Figure 4.**
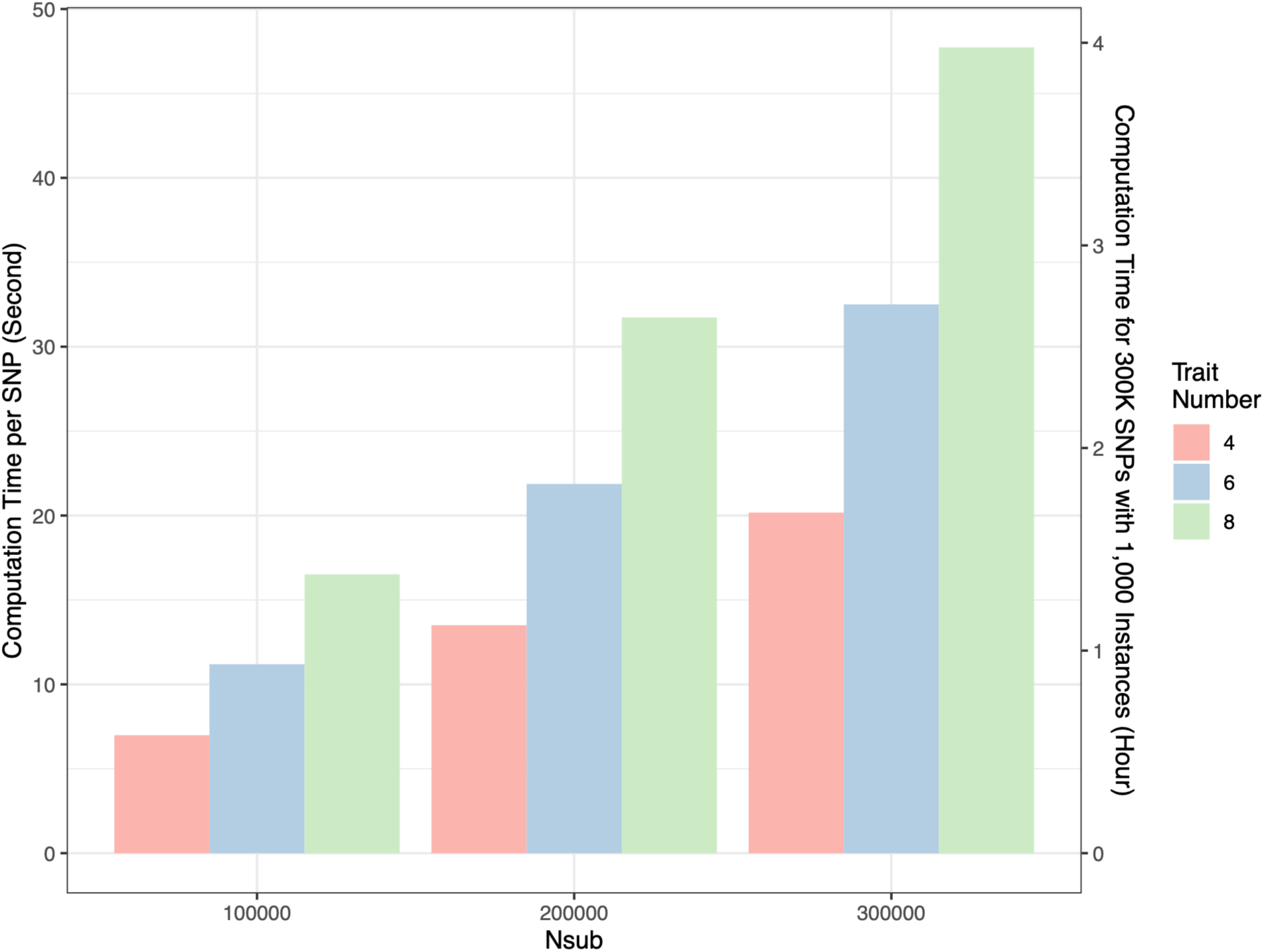
Computational performance of SCAMPI. The x-axis represents sample size (Nsub =100,000, 200,000, or 300,000), and bar colors represent analyses of 4, 6, or 8 traits. The left y axis shows the mean computation time in seconds for analyzing one SNP, estimated from 1,000 benchmark runs. The right y axis shows the corresponding projected wall-clock time for analyzing 300,000 SNPs when the workload is distributed across 1,000 independent worker instances. Genome-wide times were obtained by linear extrapolation from the measured per-SNP runtimes and therefore represent projected rather than directly observed runtimes.

## Discussion

For some complex traits, narrow-sense heritability estimates are larger when estimated from close relatives than from distantly related individuals, although the magnitude of this discrepancy and the potential contribution of gene-gene or gene-environment interactions is likely to be trait dependent. In this work, we develop our method SCAMPI to help screen for such variants that can then be prioritized for subsequent interaction analyses using standard tools. By studying correlation patterns among multiple traits, we showed using simulated data that SCAMPI has improved power relative to univariate variance-based screening procedures. Like variance-based procedures, SCAMPI does not require the specification of the factor that interacts with the variant to influence the traits under study. This means that users do not need prior knowledge of potential interacting factors, which can often be overlooked, unavailable, or difficult to collect. Furthermore, while SCAMPI produces an omnibus test to assess whether a SNP has an interactive effect on at least one of the traits under study, the method allows a user to identify the specific traits that are driving the signal by inspection of the individual cross-product *P*-values that are aggregated to form the omnibus test. The method, implemented in R code, is scalable to biobank-scale data and can handle many phenotypes.

SCAMPI prioritizes variants with genotype dependent multivariate variability for interaction studies. While such variability can arise from SNP interaction effects, we note that it can also arise erroneously if the test SNP is in linkage disequilibrium with a nearby SNP with a large additive effect on outcomes under consideration. Conditional SCAMPI analysis that adjusts for main effects of nearby SNPs therefore provides a practical way to increase rigor and confidence that the original SNP signals identified by SCAMPI are truly due to interaction effects. In our UK Biobank application, we performed such conditional analyses and observed that all top SCAMPI signals identified in Table 2 remained significant after mean adjustment for nearby SNPs with previously reported main effects on the outcomes under study (see Supplementary Figure S17 and Supplementary Table S17). Moreover, over 80% of the complete set of SCAMPI signals shown in Supplementary Table S9 remained significant in conditional analyses.

Phenotype scale is an important consideration when applying SCAMPI. The current implementation uses a Gaussian DGLM working model. Direct and indirect INT produced highly concordant results in the UK Biobank application and substantially reduced inflation under the strict heavy tailed and log-normal null simulations. In the log-normal model with genotype and environmental main effects, however, transformation induced additional rejection even without an explicit interaction. Comparing scientifically appropriate phenotype scales therefore provides useful information about the stability of SCAMPI findings.

The minimum genotype group count is a useful quality control measure because SCAMPI compares three genotype categories. Our simulations support reporting all three counts and using a minimum count of 150 as a practical criterion for common variant analyses. Calibration also depends on trait dimension and the analysis setting, so an appropriate empirical null assessment is useful when genotype groups are sparse. The additional simulations showed that SCAMPI retained a power advantage at residual correlations as high as 0.9, although high correlation reduced power. Heterogeneity in environmental main effects had little influence when the environmental variable was included in the DGLM covariate adjustment.

Our SCAMPI framework complements a recent kernel-based method Latent Interaction Testing (LIT) for interaction testing that used kernel distance covariance techniques to test whether similarity of sample trait correlation patterns correlate with genotype similarity at a test SNP.^35^ SCAMPI has practical features that LIT lacks, including the ability to directly assess which phenotypes among those modeled demonstrate interaction effects (as illustrated in Table 2 and Supplementary Table S14). Additionally, because SCAMPI is based on aggregating results across multiple cross-trait ANOVA tests, it can handle missing data more efficiently than LIT (which requires complete information on all traits for inference). To illustrate, suppose we have a sample where N subjects possess information on two phenotypes while only half of these subjects further possess additional information on a third phenotype. For joint analysis of all 3 phenotypes, LIT only considers the N/2 subjects with complete trait data for inference. SCAMPI, on the other hand, can incorporate the remaining N/2 subjects that have only information on phenotypes 1 and 2 within its cross-trait statistic. The flexible regression framework that forms the backbone of SCAMPI also enables extensions to perform interaction screening for a variety of other study designs used in genetic projects, including longitudinal and family-based designs. Moreover, SCAMPI can be extended to meta-analysis settings where individual-level data cannot be shared across studies. We will explore these SCAMPI extensions in future work. The SCAMPI R Package is available for installation on GitHub: https://github.com/epstein-software/SCAMPI.

## Funding Support

This work was supported by NIH grants R01 AG071170 (AJB, SB, DJC, MPE), R01 AG098817 (MPE), R01HL173044 (ZG, XL), R01AG085581 (ZG, XL), and R01 AG075827 (TSW, APW).

## Notes

### Competing Interest Statement

The authors have declared no competing interest.

### Summary of Updates

We performed a variety of additional simulations requested in peer review to support the power and validity of our approach and further conducted an assortment of sensitivity analyses to emphasize the robustness of our applied findings in the UK Biobank

