## Supplementary Text and Figures for "SCAMPI: A scalable statistical framework for genome-wide interaction testing harnessing cross-trait correlations"

**S1. Narrow-sense heritability estimates are inflated by higher-order genetic interactions to a greater degree in close than distant relatives**

For a relative pair with trait values $Y_1$ and $Y_2$, the covariance between $Y_1$ and $Y_2$ follows:[1]

$$Cov(Y_1, Y_2) = rV_A + k_2V_D + r^2V_{AA} + rk_2V_{AD} + (k_2)^2V_{DD} + r^3V_{AAA} + \cdots \quad \text{(Eq. S1)}$$

where $r$ is the coefficient of relatedness (genome fraction of the pair shared identical by descent (IBD)) and $k_2$ is the Cotterman coefficient (genome fraction where the pair share both alleles by IBD). $V_A$ and $V_D$ are the additive and dominance genetic variance of the trait, respectively. $V_{AA}$ and $V_{AAA}$ represent the two-way and three-way additive interaction variance of the traits, while $V_{AD}$ and $V_{DD}$ represent the two-way additive-dominance and dominance-dominance variance of the traits. Note that additional higher order multi-way additive, additive-dominance, and dominance-dominance variance effects also exist but are not explicitly stated in (Eq. S1) for brevity.

Estimation of narrow-sense heritability requires estimation of the additive genetic variance $V_A$ shown in Eq. S1. Suppose we estimate this quantity using parent-offspring pairs (close relatives), such that $r = 0.5$ and $k_2 = 0$. In this situation, studies often ignore interaction terms and estimate the additive genetic variance as twice the estimated covariance between the trait value of a parent $Y_P$ and its offspring $Y_O$, that is $\hat{V}_A$=$\widehat{Cov}(Y_P, Y_O)$. Comparing this quantity to the formula in (Eq. S1), we observe that $\hat{V}_A$ inflates the true $V_A$ by $\frac{1}{2}V_{AA} + \frac{1}{4}V_{AAA} + \frac{1}{8}V_{AAAA} + \cdots$. Now, suppose we instead estimate $V_A$ using GWAS data from 'distant' relatives (using GCTA or analogous method). In this situation, the samples are putatively unrelated such that $r$ is likely on the order of approximately $\frac{1}{100}$ and $k_2$ is nearly 0. In this case, based on (Eq. S1), $\hat{V}_A$ inflates the true $V_A$ by approximately $\frac{1}{100}V_{AA} + \left(\frac{1}{100}\right)^2 V_{AA} + \left(\frac{1}{100}\right)^3 V_{AAA} + \left(\frac{1}{100}\right)^4 V_{AAAA}$ ..... .

By comparing the $\hat{V}_A$ inflation for parent-offspring pairs (close relatives) to that using GCTA (distant relatives), it becomes apparent that the inflation among the former group will be substantially greater than the latter group when two-way, three-way, and higher-order additive interaction variance exist. This will also lead to greater inflation in narrow-sense heritability estimates when using close relatives for inference compared to distant relatives. Thus, this result suggests that the empirical discrepancy in narrow-sense heritability estimates using close relatives versus distant relatives can possibly be due to interaction effects.

#### S2. Detailed derivation of Eq. (1)

Given

$$Y_{i,1} = \alpha_1 + \beta_1 G_i + \gamma_1 W_i + \delta_1 G_i W_i + \epsilon_{i,1}$$

and

$$Y_{i,2} = \alpha_2 + \beta_2 G_i + \gamma_2 W_i + \delta_2 G_i W_i + \epsilon_{i,2}.$$

$$cov\left(Y_{i,1}, Y_{i,2} \middle| G_i = g_i\right)$$

$$= cov\left(\alpha_1 + \beta_1 g_i + \gamma_1 W_i + \delta_1 g_i W_i + \epsilon_{i,1}, \alpha_2 + \beta_2 g_i + \gamma_2 W_i + \delta_2 g_i W_i + \epsilon_{i,2}\right)$$

$$= cov(\alpha_1, \alpha_2) + cov(\alpha_1, \beta_2 g_i) + cov(\alpha_1, \gamma_2 W_i) + cov(\alpha_1, \delta_2 g_i W_i)$$

$$+ cov\left(\alpha_1, \epsilon_{i,2}\right) + cov(\beta_1 g_i, \alpha_2) + cov(\beta_1 g_i, \beta_2 g_i) + cov(\beta_1 g_i, \gamma_2 W_i)$$

$$+ cov(\beta_1 g_i, \delta_2 g_i W_i) + cov\left(\beta_1 g_i, \epsilon_{i,2}\right) + cov(\gamma_1 W_i, \alpha_2) + cov(\gamma_1 W_i, \beta_2 g_i)$$

$$+ cov(\gamma_1 W_i, \gamma_2 W_i) + cov(\gamma_1 W_i, \delta_2 g_i W_i) + cov\left(\gamma_1 W_i, \epsilon_{i,2}\right) + cov(\delta_1 g_i W_i, \alpha_2)$$

$$+ cov(\delta_1 g_i W_i, \beta_2 g_i) + cov(\delta_1 g_i W_i, \gamma_2 W_i) + cov(\delta_1 g_i W_i, \delta_2 g_i W_i)$$

$$+ cov\left(\delta_1 g_i W_i, \epsilon_{i,2}\right) + cov\left(\epsilon_{i,1}, \alpha_2\right) + cov\left(\epsilon_{i,1}, \beta_2 g_i\right) + cov\left(\epsilon_{i,1}, \gamma_2 W_i\right)$$

$$+ cov\left(\epsilon_{i,1}, \delta_2 g_i W_i\right) + cov\left(\epsilon_{i,1}, \epsilon_{i,2}\right)$$

$$= cov(\gamma_1 W_i, \gamma_2 W_i) + cov(\gamma_1 W_i, \delta_2 g_i W_i) + cov(\delta_1 g_i W_i, \gamma_2 W_i)$$

$$+ cov(\delta_1 g_i W_i, \delta_2 g_i W_i)$$

$$= \gamma_1 \gamma_2 cov(W_i, W_i) + \gamma_1 \delta_2 g_i cov(W_i, W_i) + \gamma_2 \delta_1 g_i cov(W_i, W_i)$$

$$+ \delta_1 \delta_2 g_i^2 cov(W_i, W_i)$$

$$= \gamma_1 \gamma_2 + (\gamma_1 \delta_2 + \gamma_2 \delta_1) g_i + \delta_1 \delta_2 g_i^2$$

and

$$var(Y_{i,1}|G_i = g_i) = var(\alpha_1 + \beta_1 g_i + \gamma_1 W_i + \delta_1 g_i W_i + \epsilon_{i,1})$$

$$= \gamma_1^2 var(W_i) + \delta_1^2 g_{i,j}^2 var(W_i) + 2\gamma_1\delta_1 g_{i,j} cov(W_i, W_i) + 1$$

$$= \gamma_1^2 + \delta_1^2 g_{i,j}^2 + 2\gamma_1\delta_1 g_{i,j} + 1$$

$$= \left(\gamma_1 + \delta_1 g_{i,j}\right)^2 + 1$$

As $cor(Y_{i,1}, Y_{i,2}|G_i = g_i) = \frac{cov(Y_{i,1}, Y_{i,2}|G_i = g_i)}{\sqrt{var(Y_{i,1}|G_i=g_i)}\sqrt{var(Y_{i,2}|G_i=g_i)}}$, we can use our formulas above to show that

$$cor(Y_{i,1}, Y_{i,2}|G_i = g_i) = \frac{\gamma_1\gamma_2 + (\delta_1\gamma_2 + \delta_2\gamma_1)g_i + \delta_1\delta_2 {g_i}^2}{\sqrt{(\gamma_1 + \delta_1 g_i)^2 + 1} \times \sqrt{(\gamma_2 + \delta_2 g_i)^2 + 1}}$$

We next consider a single dichotomous exposure, consistent with the single continuous exposure considered above. Let $W_i \sim \text{Bernoulli}(p)$, where $0 < p < 1$, and assume that $W_i$ is independent of $G_i$. The error terms are mutually independent, independent of $W_i$ and $G_i$, and have mean zero and variance one. For $j = 1,2$, suppose that

$$Y_{i,j} = \alpha_j + \beta_j G_i + \gamma_j W_i + \delta_j G_i W_i + \varepsilon_{i,j}$$

Let $v = p(1-p)$. Conditional on $G_i = g$, the covariance and variances are

$$\text{Cov}(Y_{i,1}, Y_{i,2} \mid G_i = g) = (\gamma_1 + \delta_1 g)(\gamma_2 + \delta_2 g)v$$

$$\text{Var}(Y_{i,j} \mid G_i = g) = \left(\gamma_j + \delta_j g\right)^2 v + 1, \;\; j = 1,2$$

Therefore,

$$\text{Cor}(Y_{i,1}, Y_{i,2} \mid G_i = g) = \frac{(\gamma_1+\delta_1 g)(\gamma_2+\delta_2 g)v}{\sqrt{[(\gamma_1+\delta_1 g)^2 v+1][(\gamma_2+\delta_2 g)^2 v+1]}} \text{ (S2)}$$

When $\delta_1 = \delta_2 = 0$, the conditional variances, covariance, and correlation do not depend on genotype. Interaction effects can induce differences in these quantities across genotype categories, supporting the same screening principle used by SCAMPI.

**S3: Approximating the Pearson correlation coefficient for two variables**

Suppose we have two variables, $\boldsymbol{V}_1$ and $\boldsymbol{V}_2$, that are each a column vector with $N$ elements. The correlation coefficient between $\boldsymbol{V}_1$ and $\boldsymbol{V}_2$ can be estimated using the sample Pearson correlation formula. Define the row-wise pairs as $[\boldsymbol{V}_1, \boldsymbol{V}_2] = \left[(v_{1,1}, v_{2,1}), \ldots, (v_{1,n}, v_{2,n})\right]^T$. The sample Pearson correlation coefficient of $\boldsymbol{V}_1$ and $\boldsymbol{V}_2$ is expressed as:

$$r_{\boldsymbol{V}_1\boldsymbol{V}_2} = \frac{\sum_{i=1}^{n}(v_{1i} - \bar{v}_1)(v_{2i} - \bar{v}_2)}{\sqrt{\sum_{i=1}^{n}(v_{1i} - \bar{v}_1)^2} \times \sqrt{\sum_{i=1}^{n}(v_{2i} - \bar{v}_2)^2}} \quad \text{(Eq. S3)}$$

Define $s_{n,\boldsymbol{V}_1}$ and $s_{n,\boldsymbol{V}_2}$ as the sample standard deviation of the two variables, i.e., $s_{n,\boldsymbol{V}_1} = \sqrt{\frac{1}{n}\sum_{i=1}^{n}(v_{1i} - \bar{v}_1)^2}$ and $s_{n,\boldsymbol{V}_2} = \sqrt{\frac{1}{n}\sum_{i=1}^{n}(v_{2i} - \bar{v}_2)^2}$, respectively. We want to point out that the standard deviation (SD) of each variable is the average, not the Bessel-corrected average using $n - 1$. This decision is because the typical sample size is large in biobank-scale data that we consider in this work and, as such, the average and Bessel-corrected average are asymptotically equivalent. Using the SD formulas, we can rewrite (S3) as

$$\frac{\sum_{i=1}^{n}(v_{1i} - \bar{v}_1)(v_{2i} - \bar{v}_2)}{\sqrt{\sum_{i=1}^{n}(v_{1i} - \bar{v}_1)^2} \times \sqrt{\sum_{i=1}^{n}(v_{2i} - \bar{v}_2)^2}} = \frac{1}{n}\sum_{i=1}^{n}\left(\frac{v_{1i} - \bar{v}_1}{s_{n,\boldsymbol{V}_1}}\right)\left(\frac{v_{2i} - \bar{v}_2}{s_{n,\boldsymbol{V}_2}}\right) = \frac{1}{n}\sum_{i=1}^{n}\tilde{v}_{1i}\tilde{v}_{2i}$$

where we can interpret $\tilde{v}_{1i} = \frac{v_{1i}-\bar{v}_1}{s_{n,V_1}}$ and $\tilde{v}_{2i} = \frac{v_{2i}-\bar{v}_2}{s_{n,V_2}}$ as the standardized samples of $v_{1i}$ and $v_{2i}$ from $\boldsymbol{V}_1$ and $\boldsymbol{V}_2$, respectively. This step transforms $\boldsymbol{V}_1$ and $\boldsymbol{V}_2$ into $\widetilde{\boldsymbol{V}}_1$ and $\widetilde{\boldsymbol{V}}_2$, such that each has a mean of zero and a standard deviation of one. We can then approximate the sample Pearson correlation coefficient, $r_{\boldsymbol{V_1V_2}}$ in (S3) as the average of these pairwise products:

$$r_{\boldsymbol{V_1V_2}} = \frac{1}{n}\sum_{i=1}^{n}\left(\frac{v_{1i}-\bar{v}_1}{s_{n,\boldsymbol{V}_1}}\right)\left(\frac{v_{2i}-\bar{v}_2}{s_{n,\boldsymbol{V}_2}}\right) = \frac{1}{n}\sum_{i=1}^{n}\tilde{v}_{1i}\tilde{v}_{2i}$$

We note that the pairwise products of these standardized variables can also be conveniently denoted as $\widetilde{\boldsymbol{V}}_1 \odot \widetilde{\boldsymbol{V}}_2$, which is a N×1 dimensional column vector. Then, we can express the sample average of $\widetilde{\boldsymbol{V}}_1 \odot \widetilde{\boldsymbol{V}}_2$ as $\frac{1}{N}J_1^n(\widetilde{\boldsymbol{V}}_1 \odot \widetilde{\boldsymbol{V}}_2)$, where $J_1^n$ is a $1 \times n$ vector with all elements equal to 1. Using (S3), it then follows that we can express the sample Pearson correlation coefficient as

$$r_{\boldsymbol{V}_1\boldsymbol{V}_2} = \frac{1}{N}J_1^n(\widetilde{\boldsymbol{V}}_1 \odot \widetilde{\boldsymbol{V}}_2).$$

To determine if the correlation coefficient differs by genotype category, $\boldsymbol{G} \in \{0, 1, 2\}$, we can examine whether $r_{\boldsymbol{V_1V_2}}$ varies across genotype category by comparing $r_{\boldsymbol{V_1V_2,G=0}}$, $r_{\boldsymbol{V_1V_2,G=1}}$ and $r_{\boldsymbol{V_1V_2,G=2}}$, which can help to evaluate the presence of interaction effects in at least one of the two variables under study.

#### S4. UKBB sensitivity analyses investigating potential direct effects of BMI on lipid traits

We evaluated whether the SCAMPI associations from the original four-trait analyses in UKBB were dependent on BMI by applying three distinct models M0, M1, and M2 to the 288,709

UK Biobank participants. M0 was our original model that considered BMI, triglycerides, LDL cholesterol, and HDL cholesterol as outcomes. M1 included only the three lipid traits as outcomes. M2 included the same outcomes, with standardized raw BMI and its square added as covariates in both the DGLM mean and dispersion submodels. For the matched comparison, variants were required to have MAF $\geqslant$ 0.05, genotype missingness $\leqslant$ 10%, Hardy-Weinberg equilibrium *P*-value $\geqslant 1 \times 10^{-10}$, minimum genotype group count $\geqslant$ 150, and finite successful results in all three models. A total of 351,201 variants met these criteria and were included in the comparison. This analysis set included the 12 lead variants reported in the Table 2, and their M0 *P*-values agreed with the originally reported values within the reported rounding precision.

At $1.67 \times 10^{-7}$, M0, M1, and M2 identified 257, 183, and 168 significant variants and 25, 23, and 20 LD defined loci, respectively. Eighteen loci were shared across all three models. At $6.25 \times 10^{-8}$, the corresponding counts were 231, 168, and 154 variants and 23, 21, and 18 loci, with 15 loci shared across all three models. The genomic inflation factors were 0.940, 0.958, and 0.965. Loci were defined using common boundaries across models. From the union of significant variants at each threshold, variants on the same chromosome were connected when they were within 1 Mb and had pairwise genotype $r^2 \geqslant 0.1$ in the fixed sample; connected components defined loci.

**S5. UKBB sensitivity analyses adjusting for marginal effects of nearby GWAS lipid loci**

For the 11 non-APOE lead variants in Table 2, we assessed whether the SCAMPI signals persisted after adjustment for the mean effects of nearby lipid-associated variants in LD. Conditioning variants were selected based on their location within 500 kb of the SCAMPI lead variant, association with HDL-C, LDL-C, or triglycerides at $P < 5 \times 10^{-8}$ in the Global Lipids Genetics Consortium GWAS, [2] and availability of directly genotyped data in our UK Biobank sample. The SCAMPI test variant itself was excluded, and redundant conditioning variants were removed at $r^2 \geq 0.80$ using LD estimated in the analysis sample. The selected variants were included as additive covariates in the DGLM mean model. For each lead variant, the unadjusted and conditioned analyses used the same participants; all of whom had observed genotypes for the tested variant and all conditioning variants. Overall, we observed that all 11 lead variants remained significant after conditioning on nearby lipid-associated variants, with adjusted $P$-values ranging from $6.58 \times 10^{-38}$ to $1.49 \times 10^{-7}$. We report the complete results in Supplementary Table S17. We further report conditioning results for all 210 significant SNPs in our original SCAMPI analysis in Supplementary Table S9.

For conditional analysis of 47 SNPs found in APOC1 (including significant variants shown in Table 2 and Supplementary Table S9), we performed conditional analysis adjusting for canonical APOE genotype effects on phenotype means, using the joint genotypes at rs429358 and rs7412. Each canonical variant was coded 0, 1, or 2, and the joint unphased genotype was treated as a categorical covariate. The six common categories were (0, 0), (1, 0), (2, 0), (0, 1), (1, 1), and (0, 2), with (0, 0) as the reference. Five indicator variables were included in the DGLM mean model. Of the 288,709 participants, 242,399 had non-missing genotypes at both rs429358 and rs7412. Twelve participants belonged to the rare categories (2, 1), (1, 2), or (2, 2), leaving 242,387 participants in the six common categories before variant specific genotype missingness was applied. The common category counts were 140,878, 57,997, 5,914, 29,942, 6,238, and 1,418, respectively. The (1, 1) category is phase ambiguous and therefore was not assigned to a specific diplotype.

For each of the 47 variants, we fitted an unadjusted model and a model with mean adjustment for the canonical APOE genotype effects. All 47 variants passed subset quality control and completed successfully; variant specific sample sizes ranged from 239,203 to 242,342. We show unadjusted and conditional results in Supplementary Figure S17. The lead variant rs445925 had $P = 8.10 \times 10^{-61}$ in the original full sample, $P = 3.51 \times 10^{-52}$ in the matched subset without joint genotype adjustment, and $P = 9.41 \times 10^{-39}$ after mean adjustment. Fourteen of the 47 variants were significant before adjustment at both $1.67 \times 10^{-7}$ and $6.25 \times 10^{-8}$. After mean adjustment, 10 and nine remained significant at these thresholds, respectively. The regional signal therefore persisted after adjustment for APOE genotype effects on phenotype means.

**S6: Derivation of Levene's test for a single trait**

Assume a sample of size $N$ divided into k subgroups, where $N_i$ is the sample size of the $i^{\text{th}}$ subgroup. To test whether variance of a trait $Y$ differs by subgroup, we define the Levene's test statistic as:

$$W = \frac{(N-k)\sum_{i=1}^{k} N_i(\bar{Z}_{i\cdot} - \bar{Z}_{\cdot\cdot})^2}{(k-1)\sum_{i=1}^{k}\sum_{j=1}^{N_i}\left(Z_{ij} - \bar{Z}_{i\cdot}\right)^2}$$

- $Z_{ij} = |Y_{ij} - \bar{Y}_{i\cdot}|$: $Y_{ij}$ the value of $Y$ for the $j$th observation of the $i$th subgroup
- $\bar{Y}_{i\cdot}$: the mean of the $i$th subgroup
- $\bar{Z}_{i\cdot}$: the subgroup means of the $Z_{ij}$
- $\bar{Z}_{\cdot\cdot}$: overall means of the $Z_{ij}$

The Levene test rejects the hypothesis that the variances are equal if $W > F_{(\alpha,k-1,N-k)}$ where $F_{(\alpha,k-1,N-k)}$ is the upper critical value of the F distribution with $k-1$ and $N-k$ degrees of freedom at a significance level of $\alpha$.

**S7: Impact of genotype-group sparsity on SCAMPI type-I error and power**

Because SCAMPI models genotype as a categorical variable with three groups, its finite sample behavior may depend on the number of participants in the least frequent genotype group. We therefore defined the minimum genotype group count as $C_{\min} = \min(n_{G=0}, n_{G=1}, n_{G=2})$, where $n_{G=g}$ denotes the number of participants with genotype dosage $g$. Compared with minor allele frequency or minor allele count alone, $C_{\min}$ directly measures the amount of information available for estimating effects associated with the least frequent genotype category.

We first conducted an exploratory screening experiment to identify the approximate range in which sparse genotype groups began to affect the calibration of SCAMPI. Genotypes were generated under Hardy-Weinberg equilibrium. The screening experiment included 19 combinations of sample size and minor allele frequency: minor allele frequencies of 0.01, 0.02, 0.03, 0.04, 0.05, 0.075, and 0.10 at $N = 20{,}000$; 0.01, 0.015, 0.02, 0.025, 0.03, and 0.05 at $N = 100{,}000$; and 0.005, 0.01, 0.015, 0.02, 0.03, and 0.05 at $N = 300{,}000$. Each scenario contained four traits with residual correlation $\rho = 0.25$ and was evaluated using $10{,}000$ replicates, giving 190,000 exploratory null replicates in total. This stage was used only to locate the relevant range of minimum genotype group counts and to select scenarios for the formal experiment.

The formal null validation focused on target minimum genotype group counts of 100, 150, and 200. Under Hardy–Weinberg equilibrium, for a target minimum genotype group count $c$ and sample size $N$, the minor allele frequency was set to $p = \sqrt{\frac{c}{N}}$ so that the expected minor allele homozygote count was equal to $c$. The realized minimum genotype group count was retained for quality control. The primary experiment used $N = 100{,}000$, $\rho = 0.25$, and 2, 4, or 8 traits, producing nine primary scenarios. Additional sensitivity experiments evaluated $\rho = 0.01$ and $\rho = 0.50$ at $N = 100{,}000$ and target count 150 for 2, 4, and 8 traits. Sample size sensitivity was evaluated at $N = 20{,}000$, $100{,}000$, and $300{,}000$ with four traits, $\rho = 0.25$, and target count 150.

In total, the formal null validation included 17 scenarios with 100,000 replicates per scenario, corresponding to 1.7 million null replicates. The remaining phenotype generating model and coefficient distributions followed the simulation framework described in the main Methods. No genotype by factor interaction was included under the null hypothesis.

For each scenario, empirical type I error was calculated as the proportion of valid replicates with a SCAMPI interaction test $P$-value no greater than the nominal significance level. Results were evaluated at nominal significance levels of 0.05, 0.01, and 0.001, with 0.01 specified as the primary level. Exact two-sided 95% Clopper-Pearson confidence intervals were calculated from the number of rejected null replicates. All 1.7 million formal null replicates completed successfully, with no failed model fits or missing genotype categories. Detailed results are reported in Table S2 and Figure S6A.

In the primary scenarios, empirical type I error at the nominal level of 0.01 ranged from 0.01051 to 0.01396. At a target minimum genotype group count of 150, the corresponding rates ranged from 0.01033 to 0.01393 across the residual correlation sensitivity scenarios. With four traits, $\rho = 0.25$, and a target count of 150, empirical type I error was 0.01151, 0.01175, and 0.01155 at sample sizes of 20,000, 100,000, and 300,000, respectively. These findings indicate that calibration was determined primarily by the amount of information in the least frequent genotype group rather than by total sample size alone. Calibration improved as the minimum genotype group count increased, although mild residual inflation remained in settings with more traits or stronger residual correlation.

We next evaluated power at target minimum genotype group counts of 30, 75, 100, 150, 200, and 250. These simulations used $N = 100{,}000$, four traits, and $\rho = 0.25$. Interaction effects of 0.025, 0.05, 0.075, 0.10, and 0.15 were applied to half of the traits. Each combination of minimum genotype group count and interaction effect was evaluated using 10,000 alternative replicates, giving 300,000 alternative replicates in total. To separate power from residual variation in null calibration, each target count was paired with 100,000 null replicates generated under the same design. The empirical significance threshold for each count was defined as the first percentile of its matched null *P*-value distribution. Empirical power was then calculated as the proportion of alternative replicates with *P*-values below this threshold, and exact two-sided 95% Clopper–Pearson confidence intervals were calculated. Detailed results are reported in Table S4 and Figure S6B.

Under the strongest simulated interaction effect of 0.15, calibrated power increased from 0.0370 at a target minimum genotype group count of 30 to 0.1238 at a count of 250. The corresponding values were 0.0584, 0.0673, 0.0901, and 0.1000 at counts of 75, 100, 150, and 200, respectively. Weaker interaction effects produced power estimates close to the empirical significance level. These results support reporting all three genotype group counts and using a minimum count of 150 as a practical quality control criterion for common variant analyses. Calibration also depends on trait dimension and the analysis setting.

#### S8. Impact of phenotype transformation on non-normal outcomes

We performed simulations to evaluate whether phenotype transformation could induce or attenuate apparent SCAMPI signals under non-normal outcomes. Each replicate included 20,000 participants, four traits, MAF = 0.10, residual correlation 0.30, an observed standard normal environmental covariate, and trait-specific environmental main effects of 0.20, 0.30, 0.40, and 0.50. Residuals followed a multivariate normal, standardized t distribution with three degrees of freedom, or standardized log-normal distribution. For each distribution, we evaluated a strict null without genotype effects, a genotype main effect model without an interaction, and interaction effects of 0.08 or 0.12. The same generated data were analyzed on the pre-INT, direct INT, and indirect INT scales. Each strict null and genotype main effect setting used 100,000 paired replicates, and each interaction setting used 10,000 paired replicates. Rejection rates were evaluated at $\alpha = 0.05$, 0.01, and 0.001, with exact two-sided 95% Clopper-Pearson confidence intervals.

We provide complete simulation results in Supplementary Table S5. For a strict null model with no genotypic main effects, we observed that use of pre-transformed outcomes yields type-I error inflation with, at nominal $\alpha = 0.01$, pre-INT rejection rates of 0.05828 under t3 distribution and 0.05945 under log-normal distribution. However, Direct INT reduced these rejection rates to 0.01299 and 0.01193, and indirect INT reduced such rates to 0.01174 and 0.01156. For null interaction simulations involving genotype main effects, we observed rejection rates for the t3 distribution of 0.05848, 0.01281, 0.01125 for the pre-INT, direct INT, and indirect INT analyses; suggesting the normality transformation is effective in this context. However, under the log-normal model containing genotype main effects but no explicit interaction, the rejection rates were 0.05783, 0.07128, and 0.48001 for the pre-INT, direct INT, and indirect INT analyses. These findings show that transformation can correct severe non-normal behavior in some settings but can also induce genotype-dependent distributional differences in others.

#### S9. Impact of Direct and Mediated Effects of One Outcome on Remaining Outcomes

Following the work in S4, we performed simulations in the spirit of our UKBB analyses to investigate impact of direct and BMI-mediated interaction mechanisms on SCAMPI inference. Parameters governing the BMI to lipid relationships and the residual lipid correlation structure were estimated from the UK Biobank data. We considered three models: M0 included BMI, triglycerides, LDL cholesterol, and HDL cholesterol, M1 included the three lipid traits without BMI, M2 included the same three lipid traits and adjusted for raw standardized BMI and BMI squared through the SCAMPI covariate matrix.

We compared M0, M1, and M2 under two null models, with and without the estimated BMI to lipid path, and two alternative mechanisms: direct interactions on lipid traits and interactions acting on BMI that were transmitted to the lipid traits. Each generated BMI and lipid phenotype was separately transformed using the direct rank based inverse normal transformation applied in the UK Biobank analysis, $x_i^{\mathrm{INT}} = \Phi^{-1}\left\{\frac{\mathrm{rank}(x_i)-0.5}{n}\right\}$, before SCAMPI was fitted. M2 adjusted for raw generated standardized BMI and its squared term. Each null setting used 100,000 replicates and each alternative setting used 10,000 replicates. Empirical rejection rates were evaluated at $\alpha = 0.05$, 0.01, and 0.001 and at the two genome wide thresholds; exact two-sided 95% Clopper–Pearson confidence intervals were calculated for the null rejection rates.

We additionally simulated direct and BMI mediated interaction mechanisms using parameters estimated from the UK Biobank phenotype data. Each replicate used 288,709 individuals and MAF = 0.05. The null without a BMI to lipid path retained the specified genotype main effects, whereas the null with the empirical BMI to lipid path set genotype main effects to zero. The direct and mediated alternatives also set these genotype main effects to zero, allowing the interaction mechanisms to be examined separately. The simulation compared the same M0, M1, and M2 definitions under a null model without a causal path, a null model containing the empirically estimated BMI to lipid path, direct interaction effects on lipid traits, and interactions acting on BMI and transmitted to the lipid traits. Each generated phenotype was separately transformed by the same direct rank based inverse normal transformation used in the UK Biobank analysis. Each null setting used 100,000 replicates, and each alternative setting used 10,000 replicates.

We provide complete results of these simulations in Supplemental Table S6 and Supplemental Figure S9. Under the null without a causal path, empirical rejection rates for M0, M1, and M2 were 0.01062, 0.01058, and 0.01046 at $\alpha = 0.01$ and 0.00098, 0.00095, and 0.00094 at $\alpha = 0.001$. Under the null containing the empirical BMI to lipid path, the corresponding rates were 0.01061, 0.01044, and 0.01091 at $\alpha = 0.01$ and 0.00117, 0.00100, and 0.00109 at $\alpha = 0.001$. At an interaction effect of 0.18 and the $6.25 \times 10^{-8}$ threshold, power under a direct lipid effect was 0.542, 0.588, and 0.904 for M0, M1, and M2. When an interaction effect of 0.40 acted on BMI and was transmitted to the lipid traits, power was 1.000, 0.064, and 0.000. These results show that the three model comparison can distinguish effects that remain in lipid traits after BMI adjustment from signals that are primarily carried through BMI.

**S10. Simulations with high residual correlation and heterogeneous environmental effects**

We examined high residual correlation and heterogeneous environmental main effects using the phenotype model in Equation (5). The environmental factor followed a standard normal distribution and was included as a covariate in the DGLM mean and dispersion models for both SCAMPI and multivariate Levene testing.

The null experiments used 20,000 individuals, MAF = 0.25, and 100,000 replicates per setting. The high correlation design considered 4 or 8 traits, homogeneous environmental main effects of 0.05, and residual correlations of 0.1, 0.3, 0.5, 0.8, or 0.9. At $\alpha = 0.001$, SCAMPI type I error ranged from 0.00090 to 0.00116 for four traits and from 0.00095 to 0.00142 for eight traits. The heterogeneous main effect design used four traits, residual correlation 0.3, and four coefficient vectors: (0.05, 0.05, 0.05, 0.05), (0.25, 0.25, 0.25, 0.25), (0.05, 0.05, 0.25, 0.25), and (0.02, 0.08, 0.20, 0.35). The corresponding type I error rates were 0.00107, 0.00082, 0.00091, and 0.00118 (Supplemental Tables S7A and S8A).

The power experiments followed Figure 2, using 20,000 individuals, four traits, MAF = 0.05, and a threshold of $6.25 \times 10^{-8}$. Interaction effects ranged from 0.15 to 0.45 in increments of 0.01. Each setting contained 100,000 replicates. The high correlation experiment used the five residual correlations above, homogeneous environmental main effects of 0.05, and interactions on two of the four traits. The heterogeneous main effect experiment used the four coefficient vectors above, interactions on all four traits, and the following fixed residual correlation matrix, in the order HDL cholesterol, LDL cholesterol, triglycerides, and BMI:

$$\Sigma = \begin{pmatrix} 1 & 0.1246 & -0.4938 & -0.3809 \\ 0.1246 & 1 & 0.2797 & 0.0394 \\ -0.4938 & 0.2797 & 1 & 0.3708 \\ -0.3809 & 0.0394 & 0.3708 & 1 \end{pmatrix}.$$

These designs comprised 155 high correlation settings and 124 heterogeneous main effect settings, with 27,900,000 simulated data sets in total. Both methods returned valid results for all replicates. At $\delta = 0.45$, SCAMPI power decreased from 0.95614 to 0.53838 as residual correlation increased from 0.1 to 0.9; multivariate Levene power decreased from 0.39658 to 0.30754 across these same settings. Across the four environmental main effect patterns at $\delta = 0.35$, power ranged from 0.56838 to 0.56886 for SCAMPI and from 0.02712 to 0.02857 for multivariate Levene testing. Thus, high residual correlation reduced power, whereas differences in the adjusted environmental main effects had little influence on the power curves (Tables S7B and S8B; Figures S10–S11). Power is reported at the nominal threshold; the genotype group count and null calibration results are provided in Supplementary Tables S2–S3, S7A, and S8A.

#### S11: Significance threshold for evaluating type I error

To explain why $\alpha = 0.001$ is an adequate choice for our nominal type I error $\alpha$ level when there are 100,000 replicates, we are given $X_1, \ldots, X_m \sim_{i.i.d} Binom(n, \alpha)$ by the Central Limit Theorem, then $\sqrt{m}(\bar{X}_m - \alpha) \xrightarrow{d} \mathbb{N}(0, \alpha(1-\alpha))$, equivalently, $\frac{\bar{X}_m - \alpha}{\sqrt{\frac{\alpha(1-\alpha)}{m}}} \xrightarrow{d} \mathbb{N}(0,1)$. Thus, the asymptotic mean ($\mu$) and asymptotic variance ($\sigma^2$) of $\bar{X}_m$ are $\alpha$ and $\frac{\alpha(1-\alpha)}{m}$.

In our simulation setting, we have fixed the number of simulated replicates as $M = 100{,}000$, and we let $\alpha$ be the pre-specified threshold. This table provides the corresponding pre-specified significance level (col "$\alpha$"), asymptotic mean (col "$\mu = \alpha$"), asymptotic standard error (col "$\sigma = \sqrt{\frac{\alpha(1-\alpha)}{M}}$"), and 95% single-side CI bandwidth (col "95% CI bandwidth 1.96* $SE(\bar{X}_n)$") when the pre-specified thresholds vary at the given fixed $M = 100{,}000$.

In general, the bandwidth of the 95% CI for a proportion such as $\alpha$ should not overlap but be at least one order of magnitude smaller than the asymptotic mean.[3] In this table, when $\alpha = 0.01$ or $\alpha = 0.001$, their 95% CI bandwidths are smaller than their corresponding asymptotic mean at least at one order of magnitude. When $\alpha = 0.0001$ or smaller, their asymptotic mean and 95% CI bandwidth are very close in value and do not differ by an order of magnitude.

| $M$ | $\alpha$ | $\mu = \alpha$ | $\sigma = \sqrt{\frac{\alpha(1-\alpha)}{M}}$ | 95% CI bandwidth 1.96* $SE(\bar{X}_n)$ |
|---|---|---|---|---|
| 1E5 | 1E-2 | 1E-2 | 3.15E-4 | 6.17E-4 |
| | 1E-3 | 1E-3 | 9.99E-5 | 1.96E-4 |
| | 1E-4 | 1E-4 | 3.16E-5 | 6.20E-5 |
| | 1E-5 | 1E-5 | 1.00E-5 | 1.96E-5 |

We further evaluated the null distribution using 200 UK Biobank phenotype permutations, which yielded 46,429,908 valid SCAMPI $P$-values. The QQ plot showed no marked systematic inflation across the empirically observable range (Figure S5). Under a uniform null distribution, this number of tests gives approximately 2.90 expected exceedances at $6.25 \times 10^{-8}$ and 7.75 at $1.67 \times 10^{-7}$. The plot therefore assesses the shape of the null tail, with limited precision for estimating the rejection rate at either exact threshold.

**S12: Computational Cost**

We benchmarked SCAMPI on the High-Performance Computing (HPC) system hosted by the Emory University Rollins School of Public Health. At the time of benchmarking, the system comprised 25 compute nodes, including 24 nodes equipped with 32 CPU cores and 192 GB of memory and one high-memory node with 1.5 TB of memory. For each combination of sample size and number of traits, we measured the elapsed time required to analyze one SNP and averaged the runtime over 1,000 independent benchmark repetitions. The measured runtime increased approximately linearly with sample size and quadratically with the number of traits. For an analysis of four traits in 300,000 individuals, the mean runtime was 20.17 seconds per SNP on the RSPH HPC system. The total single-worker computational requirement for analyzing (M) SNPs can be approximated as $\left[T_{\text{compute}} = \frac{M \times t_{\text{SNP}}}{3600}\right]$, where ($t_{\text{SNP}}$) is the mean runtime per SNP in seconds. Based on the observed runtime of 20.17 seconds, analyzing 300,000 SNPs would require approximately 1,681 single-worker hours, whereas analyzing 6,000,000 SNPs would require approximately 33,617 single-worker hours. Because SNPs can be analyzed independently, SCAMPI is readily parallelizable. With (C) concurrently available workers, the idealized wall-clock time can be approximated as $T_{\text{wall}} \approx \frac{T_{\text{compute}}}{C}$. For example, the previously reported projections of 1.68 hours for 300,000 SNPs and 33.62 hours for 6,000,000 SNPs correspond to an idealized setting with 1,000 concurrently available workers. These values are computational projections rather than observed end-to-end runtimes. Actual wall-clock time depends on the available computing resources, scheduler policy, workload balance, data input/output, and other system overhead. We additionally benchmarked SCAMPI on a MacBook Pro equipped with an Apple M1 processor. For four traits and 300,000 individuals, the mean runtime was 8.65 seconds per SNP. This corresponds to approximately 721 single-worker hours for 300,000 SNPs and 14,417 single-worker hours for 6,000,000 SNPs. Under the same idealized assumption of 1,000 concurrently available workers, these computational requirements would correspond to approximately 0.72 and 14.42 hours, respectively. These results illustrate that absolute runtime depends on the hardware and software environment, whereas the observed scaling with sample size and number of traits was consistent across the evaluated platforms. Detailed results are presented in Figures 4 and S18–S19.

**Figure S1**

**(a)**

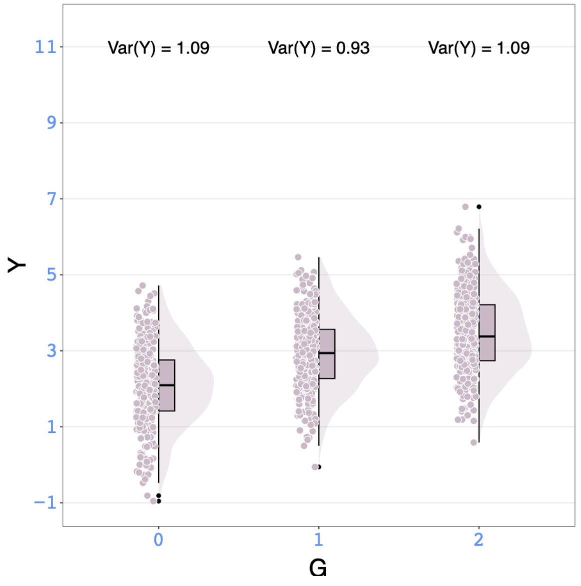


**(b)**

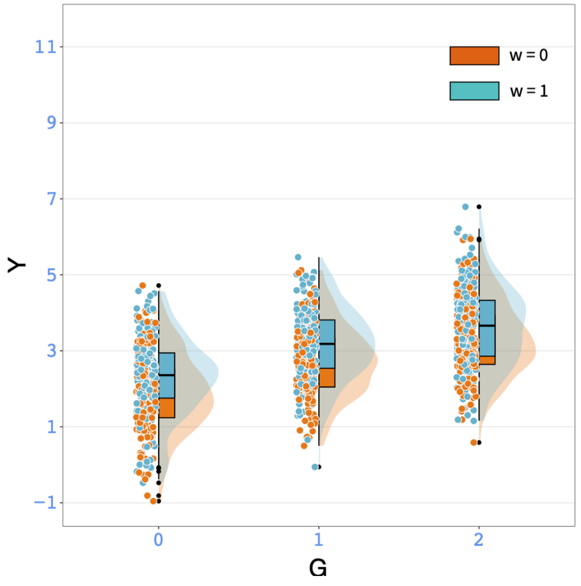


**(c)**

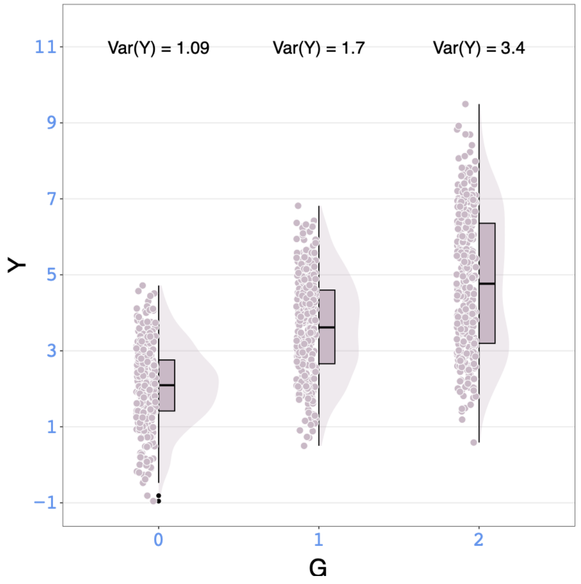


**(d)**

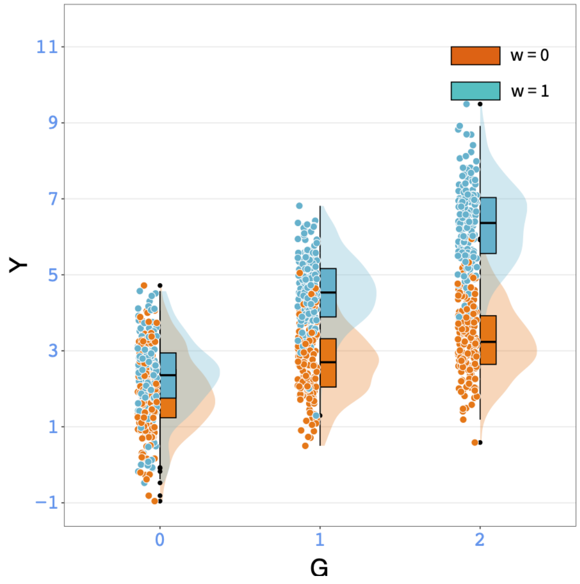


**Figure S1. Interaction effect induce variance heteroscedasticity in a phenotypic trait.**

Sub-figures (a) and (c) depict simulations of one phenotypic trait samples from $Y_i = \alpha_1 + \beta_1 G_i + \gamma_1 W_i + \epsilon_i$ (without interaction effect) and $Y_i = \alpha_1 + \beta_1 G_i + \gamma_1 W_i + \delta_1 G_i W_i + \epsilon_i$ (with interaction effect), respectively. In these figures, the interacting covariate $W_i$ is simplified as a binary factor for clarity. The genotype categories are plotted on the x-axis, and the simulated outcomes $\boldsymbol{Y}$ are on the y-axis, with variances for each genotype category annotated. The distributions of $\boldsymbol{Y}$ differ across the three genotype categories as shown in the sub-figure (c), but not observed in the sub-figure (a). The sub-figures (b) and (d) mirror (a) and (c), respectively, with the addition of labels by $\boldsymbol{W}$ using the color orange ($W_i = 0$) and blue ($W_i = 1$). By comparing the sub-figures (b) and (d), the variation in $\boldsymbol{Y}$ distributions across genotype categories, as observed in sub-figure (c), highlights the variance heterogeneity driven by the interacting effect, $\boldsymbol{GW}$.

**Figure S2.**

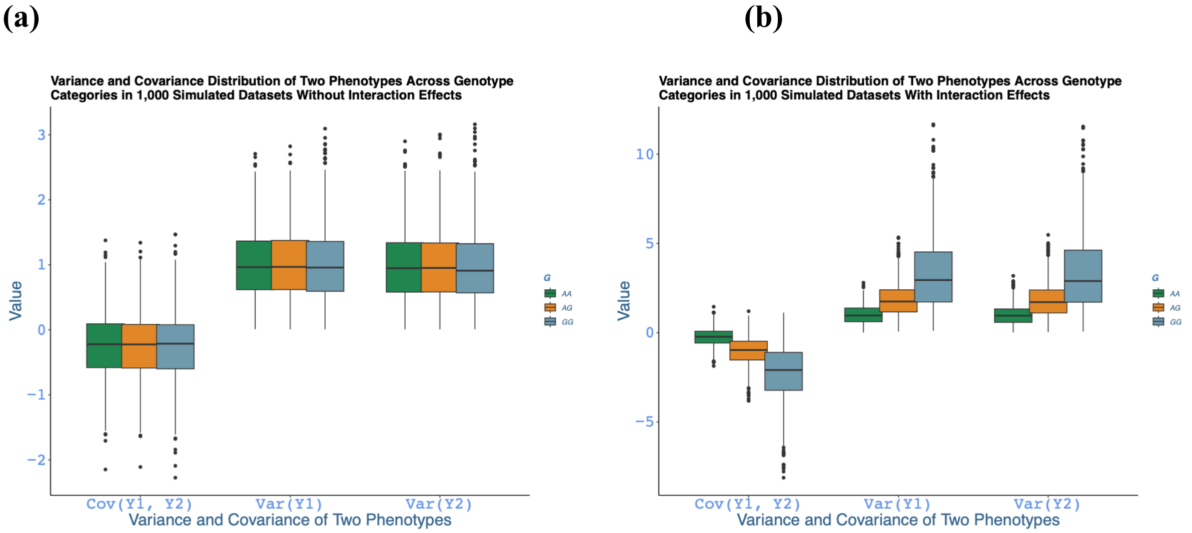


**Figure S2. Interaction effect induces both variance and covariance heteroscedasticity in two phenotypic traits.**

Sub-figures (a) and (b) depict the distribution of three covariance elements on the x-axis, including $cov(\boldsymbol{Y}_1, \boldsymbol{Y}_2)$, $var(\boldsymbol{Y}_1)$ and $var(\boldsymbol{Y}_2)$, across different genotypes, marked by varying colors within each covariance element. These measures are plotted using 1,000 datasets, and each dataset contains 1,000 samples of $\boldsymbol{Y}_1$ and 1,000 samples of $\boldsymbol{Y}_2$, which were simulated from multivariate normal distributions with and without gene-environment interaction terms:

$\begin{pmatrix} Y_{1,i} \\ Y_{2,i} \end{pmatrix} \sim MVN\left(\begin{matrix} \alpha_1+\beta_1 G_i+\gamma_1 W_i \\ \alpha_2+\beta_2 G_i+\gamma_2 W_i \end{matrix}, \Sigma = \begin{bmatrix} Unif(0,1) & Unif(0,1) \\ Unif(0,1) & Unif(0,1) \end{bmatrix}\right)$ and

$\begin{pmatrix} Y_{1,i} \\ Y_{2,i} \end{pmatrix} \sim MVN\left(\begin{matrix} \alpha_1+\beta_1 G_i+\gamma_1 W_i+\delta_1 G_i W_i \\ \alpha_2+\beta_2 G_i+\gamma_2 W_i+\delta_2 G_i W_i \end{matrix}, \Sigma = \begin{bmatrix} Unif(0,1) & Unif(0,1) \\ Unif(0,1) & Unif(0,1) \end{bmatrix}\right)$, as shown in sub-figures (a) and (b), respectively. We then can calculate the three covariance/variance components, $cov(\boldsymbol{Y}_1, \boldsymbol{Y}_2)$, $var(\boldsymbol{Y}_1)$ and $var(\boldsymbol{Y}_2)$, for each dataset. The two sub-figures draw the distribution of the covariance/variance components of the 1,000 datasets. Sub-figure (a) shows no visual distinction in the distribution of these statistical measures across genotypes when interaction is absent. In contrast, sub-figure (b) demonstrates noticeable differences in $cov(\boldsymbol{Y}_1, \boldsymbol{Y}_2)$ , $var(\boldsymbol{Y}_1)$ and $var(\boldsymbol{Y}_2)$ across genotypes when interaction effects are present.

**Figure S3.**

(a)

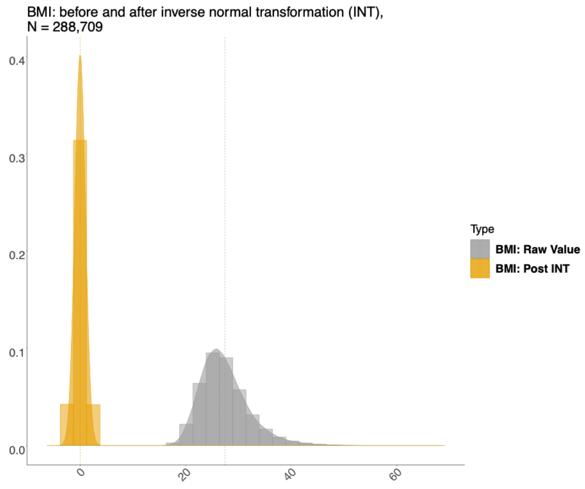


(b)

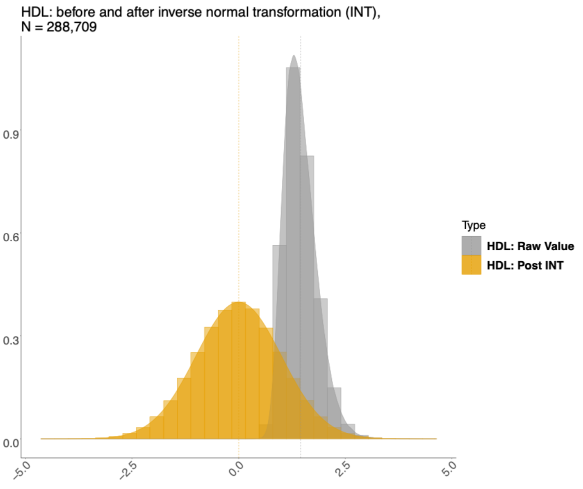


(c)

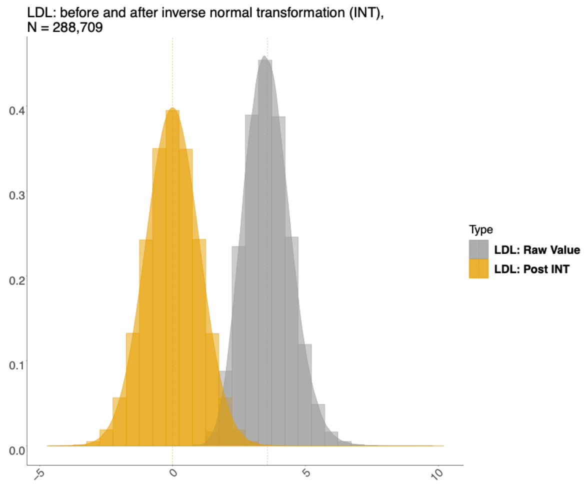


(d)

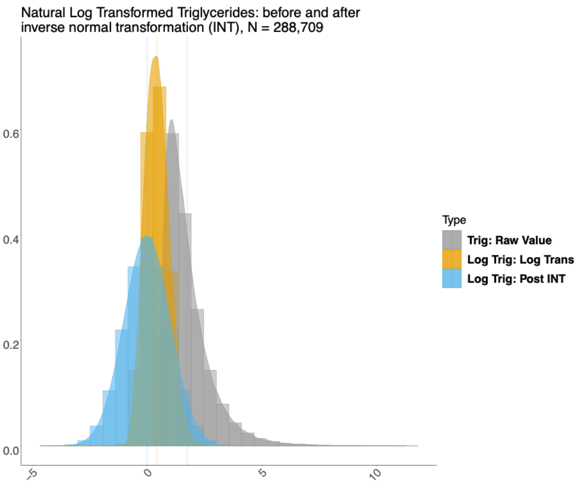


**Figure S3. Distribution of the four lipids panel-related traits before and after the inverse normal transformation (INT) in UKBB analysis**

Panels (a)-(d) show BMI, HDL-C, LDL-C, and triglycerides, respectively, among 288,709 participants. The x-axis represents the trait value on the indicated raw or transformed scale, and the y-axis represents the estimated probability density. Gray distributions show the raw trait values, and yellow distributions show values after rank-based inverse normal transformation (INT). For triglycerides in panel (d), the raw, natural log-transformed, and post-INT distributions are displayed separately. Dashed vertical lines indicate the mean of each displayed distribution.

**Figure S4.**

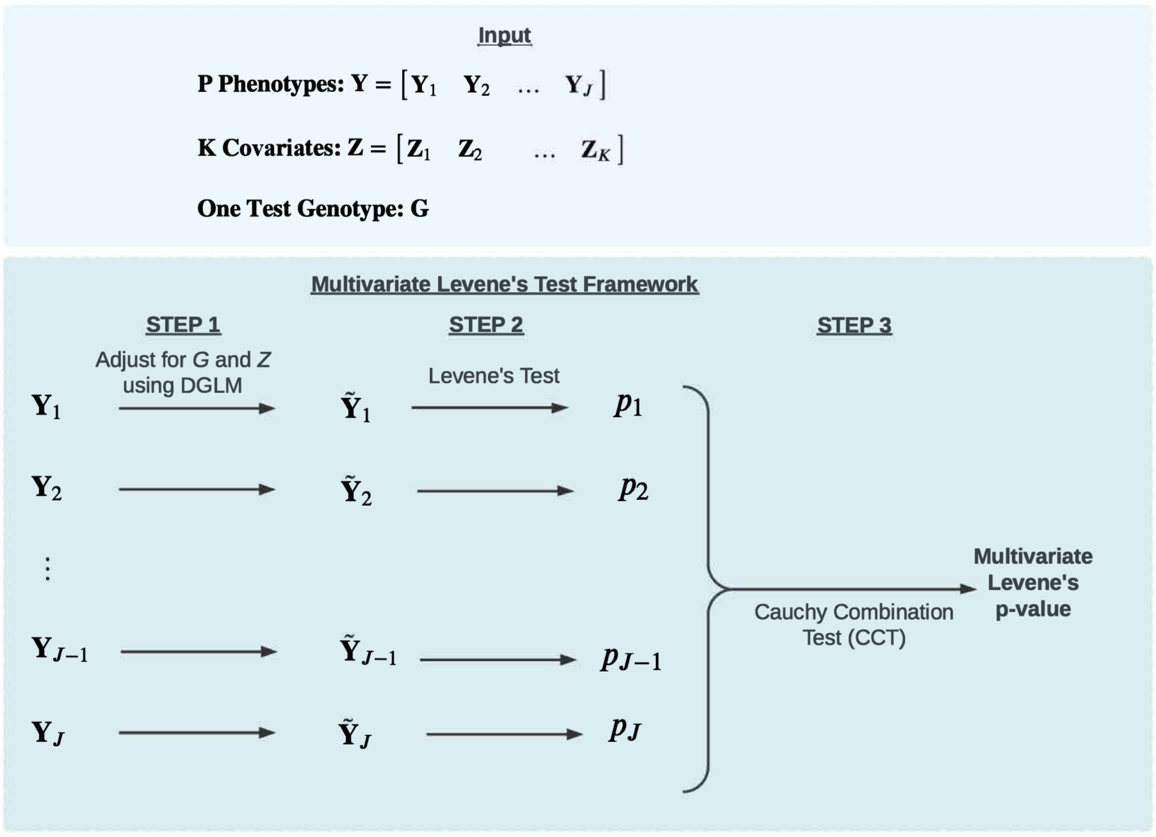


**Figure S4. Illustration of the multivariate Levene's test framework.**

Multivariate Levene's test framework is a three-step process that includes adjustment for genotype and covariates, applying univariate Levene's test for each one of the $J$ traits, and then combines the $J$ $P$-values using Cauchy Combination Test (CCT) to compute the final Multivariate Levene's $P$-value to determine overall significance.

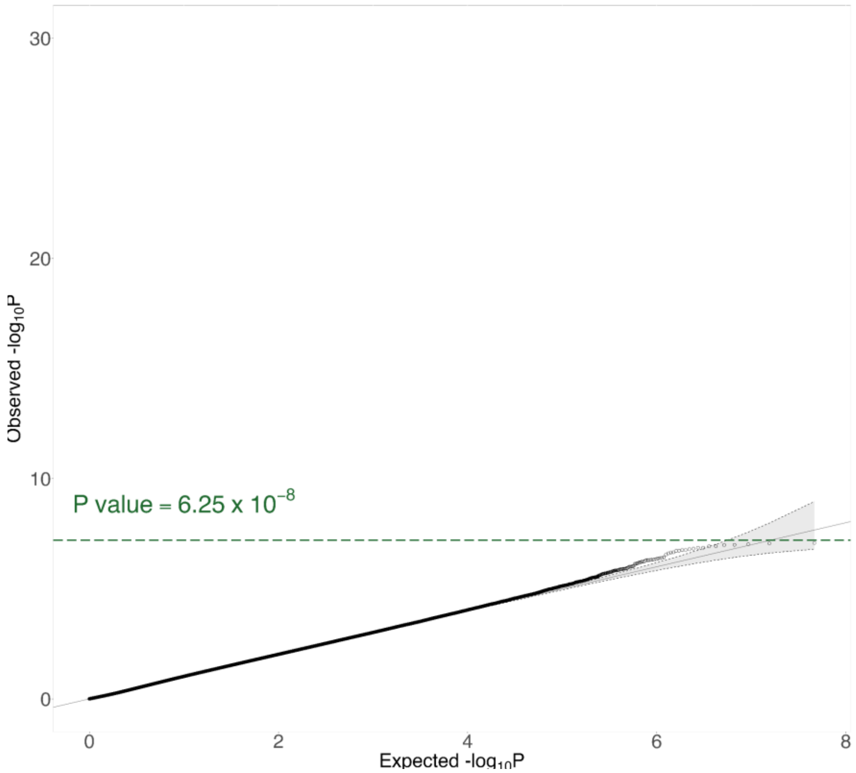


**Figure S5. Q-Q plot of SCAMPI null results from 200 UK Biobank phenotype permutations**

The x axis shows the expected −log10(*P*) under the global null hypothesis, and the y axis shows the observed −log10(*P*) across 46,429,908 valid SCAMPI tests. In each permutation, the phenotype data and covariates were permuted together across participants relative to the fixed genotype data. Variants with MAF of at least 0.05 and genotype missingness below 10% were included. The gray diagonal line represents equality between expected and observed values, and the green dashed line marks P = $6.25 \times 10^{-8}$. The permutation design and tail precision are described in Supplementary Section S11.

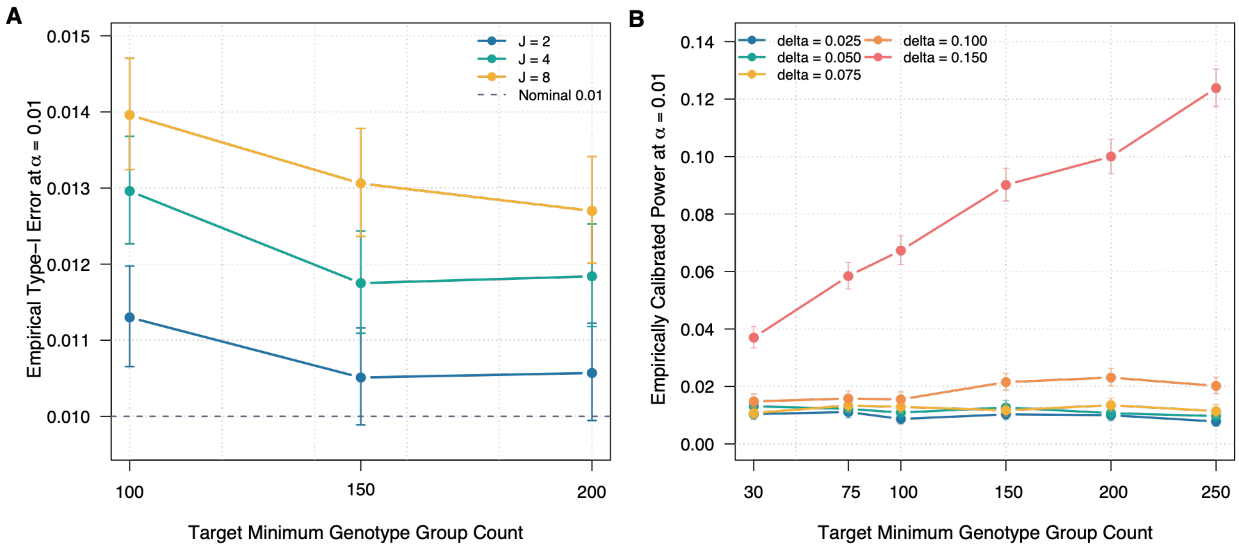


**Figure S6. Effects of minimum genotype group count on the finite-sample performance of SCAMPI**

(A) Empirical type I error at a nominal significance level of 0.01 for target minimum genotype group counts of 100, 150, and 200 and analyses of 2, 4, or 8 traits. Each setting contained 100,000 null replicates. The dashed line indicates the nominal 0.01 level.

(B) Empirically calibrated power for target minimum genotype group counts of 30, 75, 100, 150, 200, and 250 and interaction effects ranging from 0.025 to 0.15. Each alternative setting contained 10,000 replicates, and the count specific critical values were estimated from 100,000 matched null replicates. Points show empirical rejection proportions, and vertical bars show exact two-sided 95% Clopper-Pearson confidence intervals.

**Figure S7.**

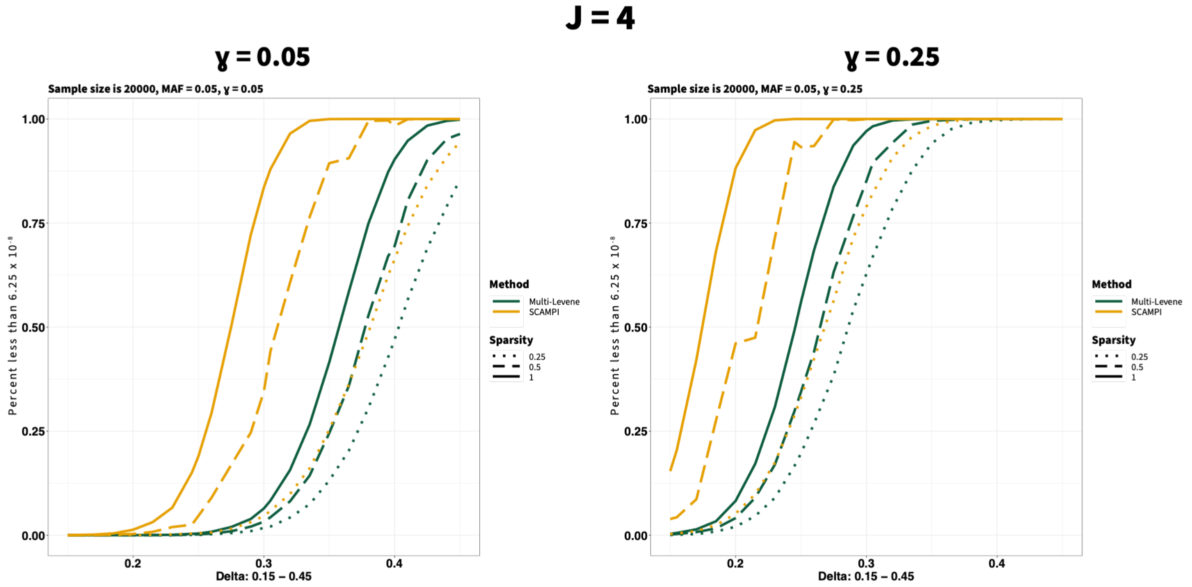


**Figure S7. Power results for UKBB inspired simulations of SCAMPI and Multivariate Levene's test varying proportions of traits with interactions.**

The x axis represents the genotype by factor interaction coefficient δ, and the y axis represents the proportion of 10,000 alternative replicates with a *P*-value below $6.25 \times 10^{-8}$. Four traits were simulated for 20,000 individuals with a minor allele frequency of 0.05 and a UK Biobank matched trait correlation matrix. The left and right panels assume factor main effects of 0.05 and 0.25, respectively. Line type represents interaction effects on 25%, 50%, or 100% of the traits. Yellow lines represent SCAMPI, and green lines represent the multivariate Levene's test. Under Hardy–Weinberg equilibrium, this design has an expected minimum genotype group count of approximately 50. The displayed values are therefore nominal alternative rejection probabilities and should be interpreted together with the matched null calibration results in Table S3.

### Figure S8.

## (a)

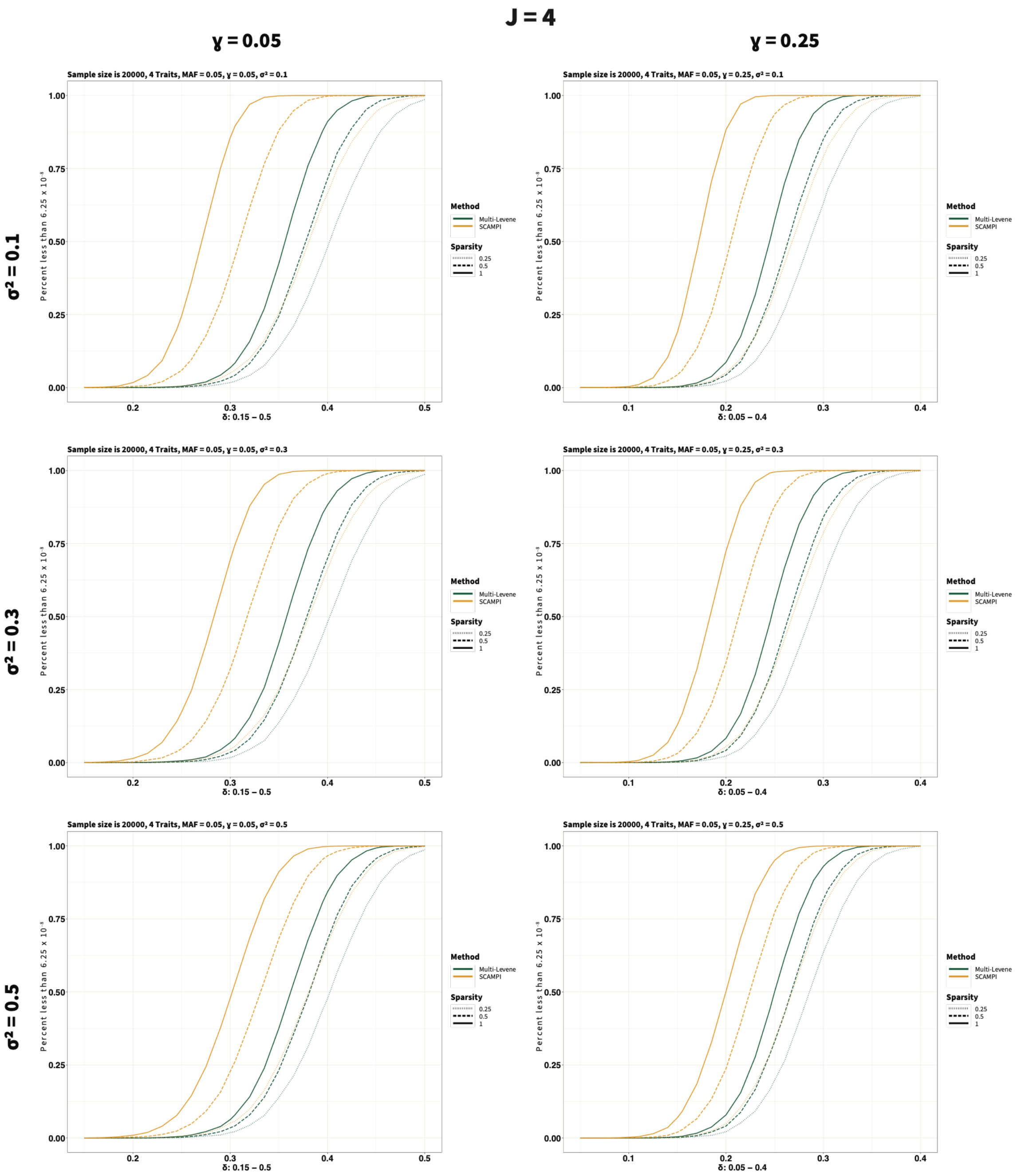
J = 4
ɣ = 0.05
ɣ = 0.25
σ² = 0.1
σ² = 0.3
σ² = 0.5
Sample size is 20000, 4 Traits, MAF = 0.05, ɣ = 0.05, σ² = 0.1
Sample size is 20000, 4 Traits, MAF = 0.05, ɣ = 0.25, σ² = 0.1
Sample size is 20000, 4 Traits, MAF = 0.05, ɣ = 0.05, σ² = 0.3
Sample size is 20000, 4 Traits, MAF = 0.05, ɣ = 0.25, σ² = 0.3
Sample size is 20000, 4 Traits, MAF = 0.05, ɣ = 0.05, σ² = 0.5
Sample size is 20000, 4 Traits, MAF = 0.05, ɣ = 0.25, σ² = 0.5
Percent less than 6.25 x 10⁻⁸
δ: 0.15 – 0.5
δ: 0.05 – 0.4
Method
Multi-Levene
SCAMPI
Sparsity
0.25
0.5
1

**(b)**

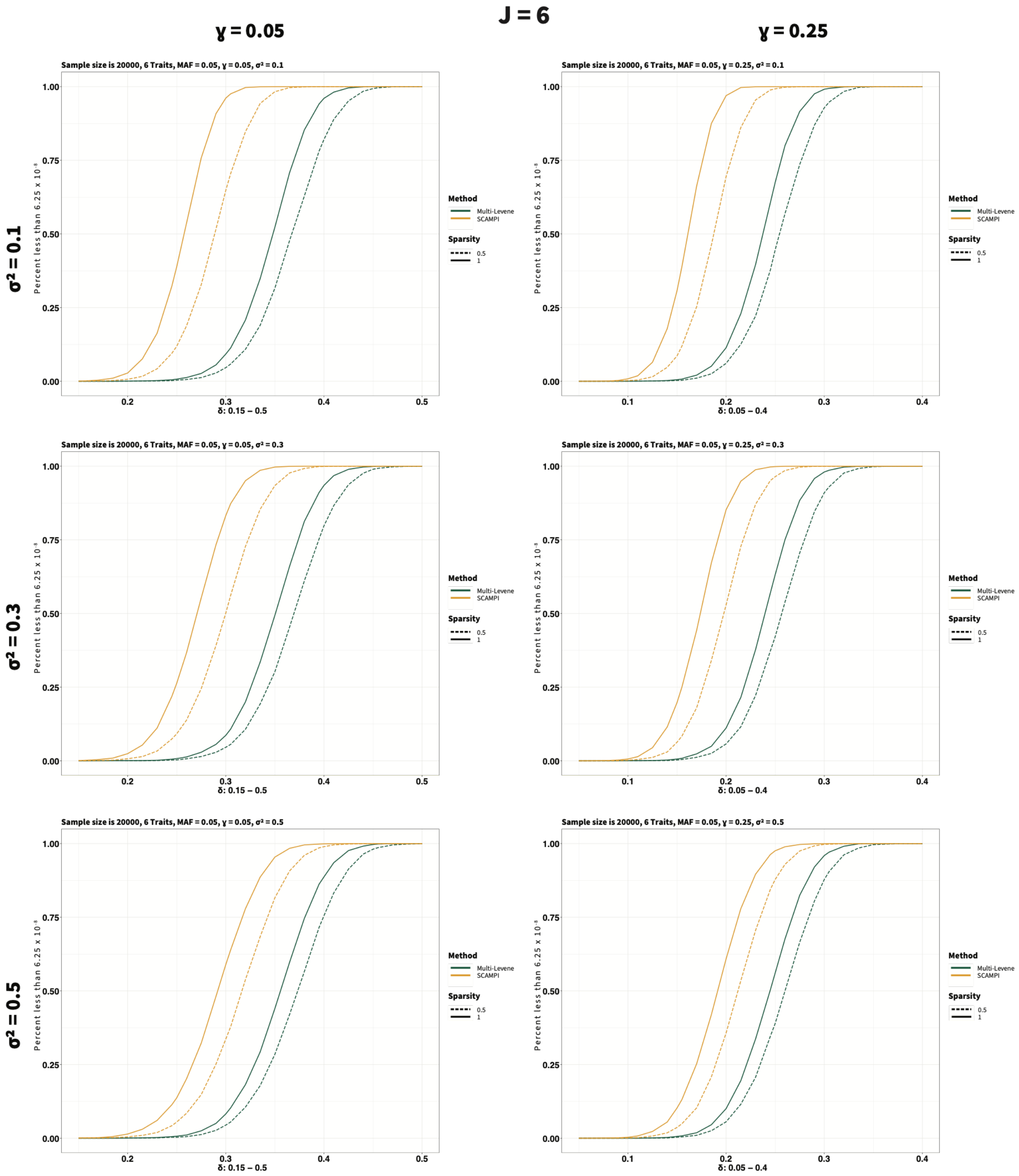
J = 6
γ = 0.05
γ = 0.25
σ² = 0.1
σ² = 0.3
σ² = 0.5
Sample size is 20000, 6 Traits, MAF = 0.05, γ = 0.05, σ² = 0.1
Sample size is 20000, 6 Traits, MAF = 0.05, γ = 0.25, σ² = 0.1
Sample size is 20000, 6 Traits, MAF = 0.05, γ = 0.05, σ² = 0.3
Sample size is 20000, 6 Traits, MAF = 0.05, γ = 0.25, σ² = 0.3
Sample size is 20000, 6 Traits, MAF = 0.05, γ = 0.05, σ² = 0.5
Sample size is 20000, 6 Traits, MAF = 0.05, γ = 0.25, σ² = 0.5
Percent less than 6.25 x 10⁻⁸
δ: 0.15 – 0.5
δ: 0.05 – 0.4
Method
Multi-Levene
SCAMPI
Sparsity
0.5
1

(c)

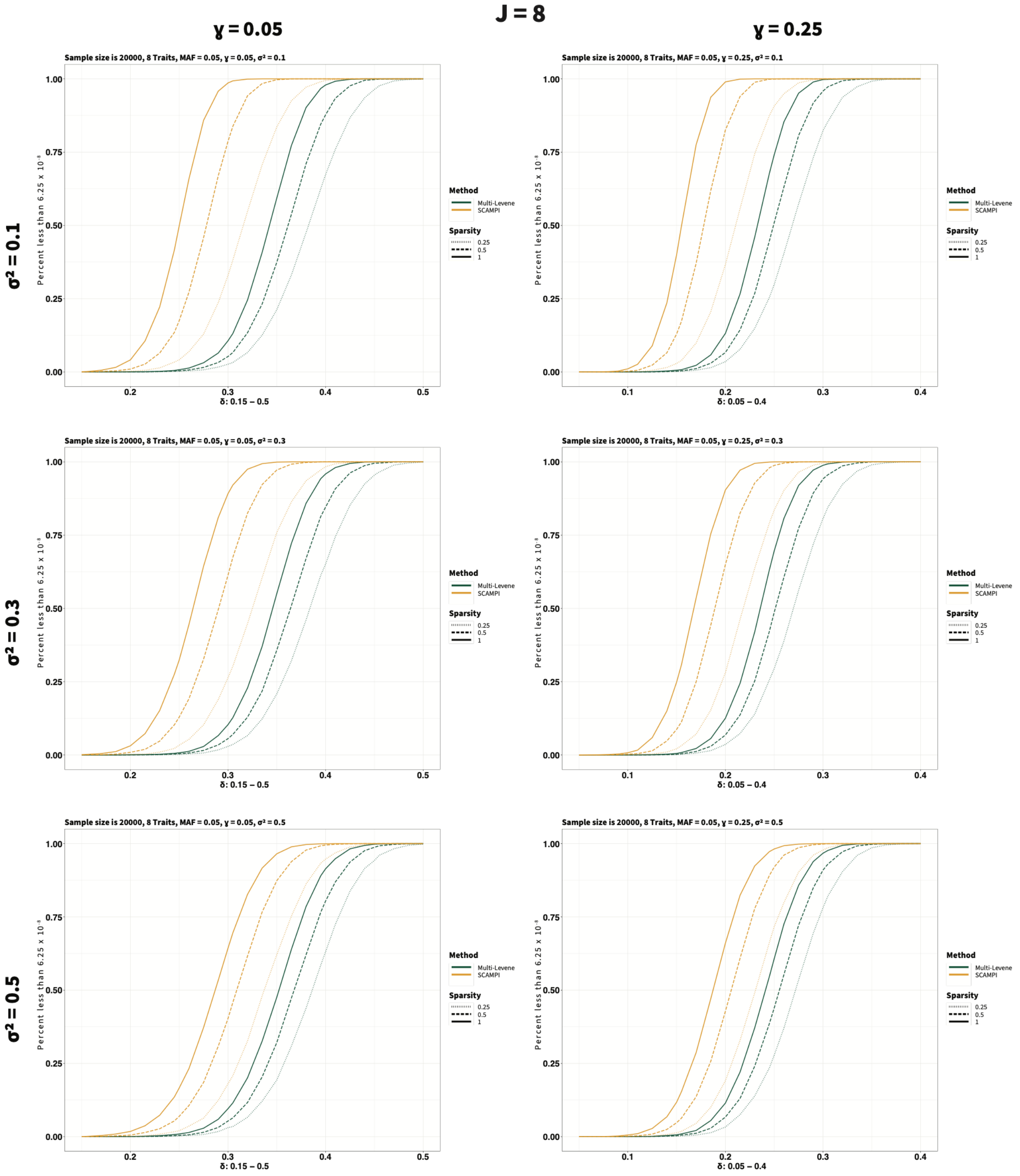

J = 8
γ = 0.05
γ = 0.25
σ² = 0.1
σ² = 0.3
σ² = 0.5
Sample size is 20000, 8 Traits, MAF = 0.05, γ = 0.05, σ² = 0.1
Sample size is 20000, 8 Traits, MAF = 0.05, γ = 0.25, σ² = 0.1
Sample size is 20000, 8 Traits, MAF = 0.05, γ = 0.05, σ² = 0.3
Sample size is 20000, 8 Traits, MAF = 0.05, γ = 0.25, σ² = 0.3
Sample size is 20000, 8 Traits, MAF = 0.05, γ = 0.05, σ² = 0.5
Sample size is 20000, 8 Traits, MAF = 0.05, γ = 0.25, σ² = 0.5
Percent less than $6.25 \times 10^{-8}$
0.00
0.25
0.50
0.75
1.00
0.1
0.2
0.3
0.4
0.5
δ: 0.15 – 0.5
δ: 0.05 – 0.4
Method
Multi-Levene
SCAMPI
Sparsity
0.25
0.5
1

**Figure S8. Additional power results for SCAMPI and Multivariate Levene's test varying proportions of traits with interactions.**

The x axis shows the genotype by factor interaction coefficient $\delta$, and the y axis shows the proportion of 10,000 alternative replicates with an omnibus *P*-value below $6.25 \times 10^{-8}$. Panels are grouped by the number of traits, with J = 4, 6, and 8 in panels (a), (b), and (c), respectively. Within each panel, the subplots represent combinations of factor main effect $\gamma = 0.05$ or 0.25 and residual correlation $\rho = 0.1$, 0.3, or 0.5. Yellow lines represent SCAMPI, green lines represent the multivariate Levene's test, and line type indicates that interactions affect 25%, 50%, or 100% of the traits. Each setting used N = 20,000 and MAF = 0.05, corresponding to an expected minimum genotype group count of approximately 50 under Hardy-Weinberg equilibrium. The displayed values are nominal alternative rejection probabilities and should be interpreted together with the matched null calibration results in Supplementary Table S3.

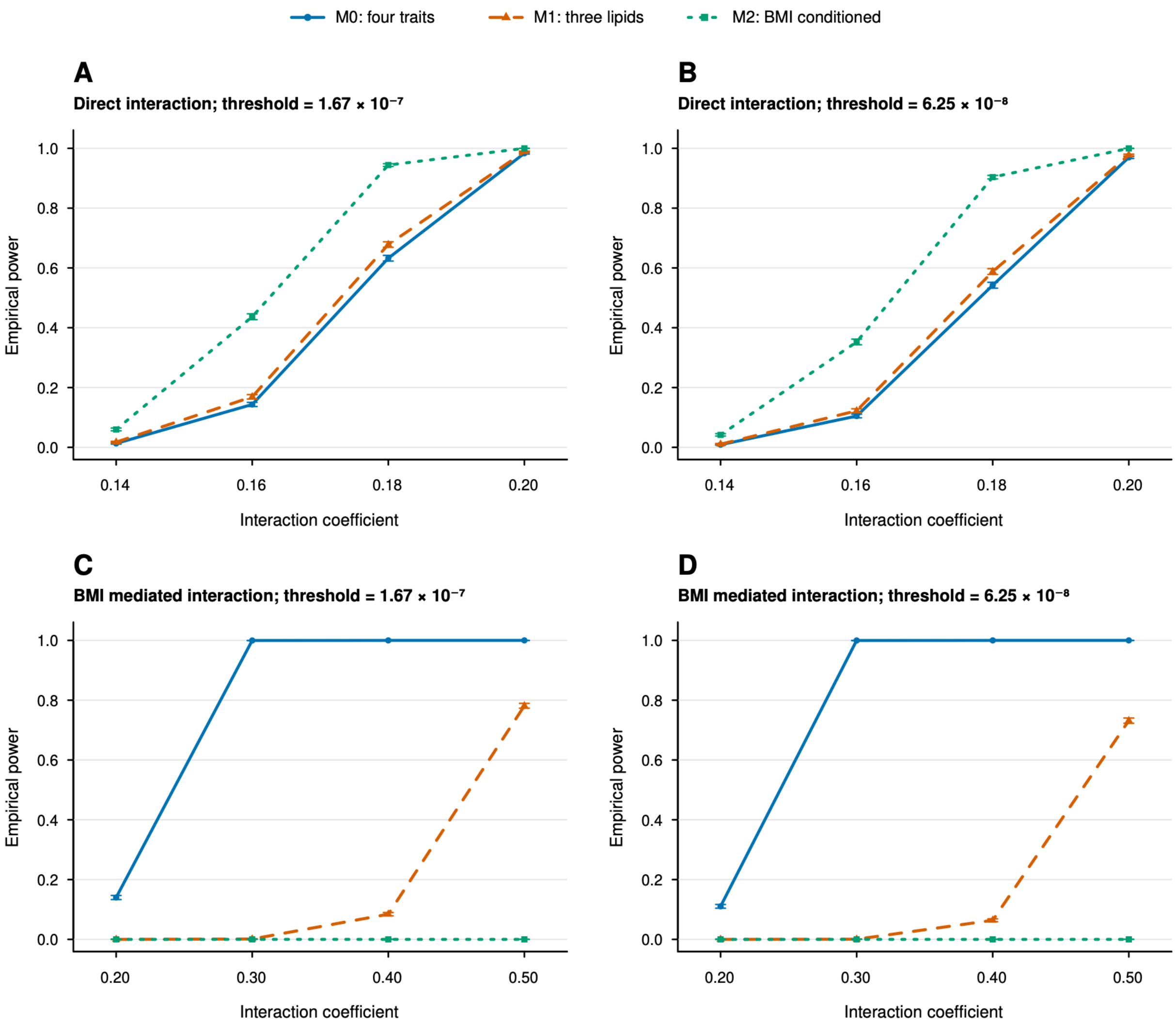


**Figure S9. Power of the original and BMI sensitivity models under direct and BMI mediated interaction mechanisms**

Panels A and B show direct lipid interactions, and panels C and D show BMI mediated interactions. Panels A and C use $P < 1.67 \times 10^{-7}$, panels B and D use $P < 6.25 \times 10^{-8}$. The x axis shows the interaction coefficient, and the y axis shows empirical power. Each generated phenotype was separately transformed using the direct rank based inverse normal transformation before SCAMPI analysis. M0 included BMI and the three lipid traits, M1 included the three lipid traits only, and M2 included the three lipid traits with adjustment for raw standardized BMI and

BMI squared. The direct mechanism applied the interaction to the lipid traits, whereas the mediated mechanism applied the interaction to BMI and transmitted its effect to the lipid traits using UK Biobank estimated coefficients. Each point is based on 10,000 replicates. Error bars show exact two-sided 95% Clopper-Pearson confidence intervals.

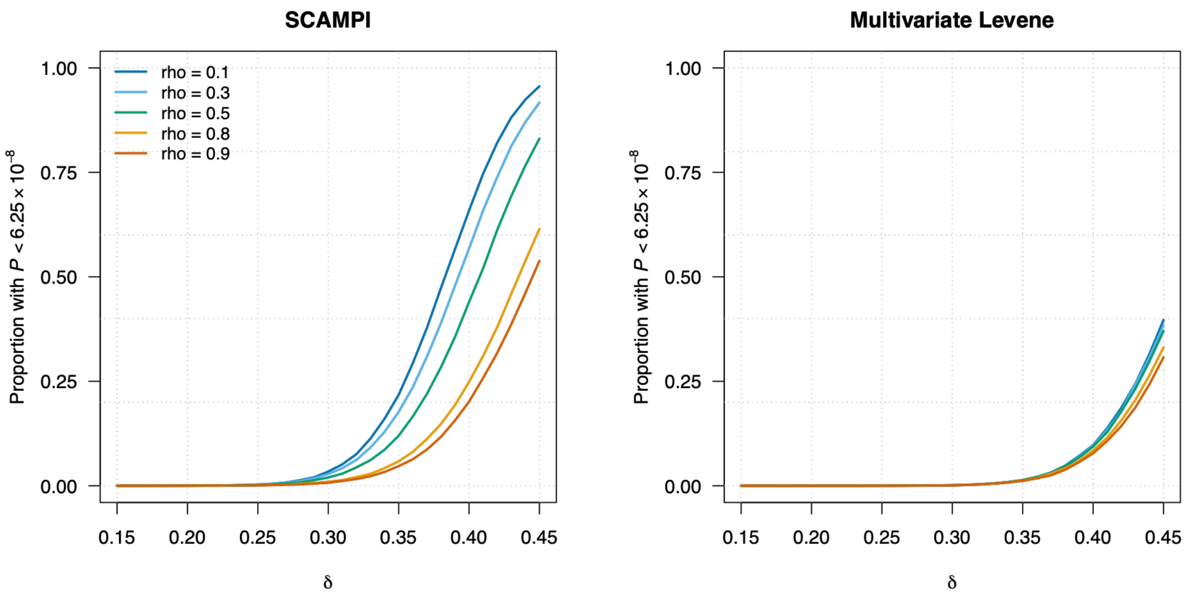


**Figure S10. Power under high residual correlation at the genome wide threshold**

Power is shown for SCAMPI and the multivariate Levene test using 20,000 individuals, four traits, MAF = 0.05, homogeneous environmental main effects of 0.05, and residual correlation $\rho$ = 0.1, 0.3, 0.5, 0.8, or 0.9. Two traits carried an interaction. The x axis shows the interaction coefficient, from 0.15 to 0.45 in increments of 0.01; the y axis shows the proportion of 100,000 alternative replicates with an omnibus *P*-value below $6.25 \times 10^{-8}$. The environmental variable was adjusted for in both methods. Power is evaluated at the nominal threshold, with null calibration results reported separately in Tables S3 and S7A.

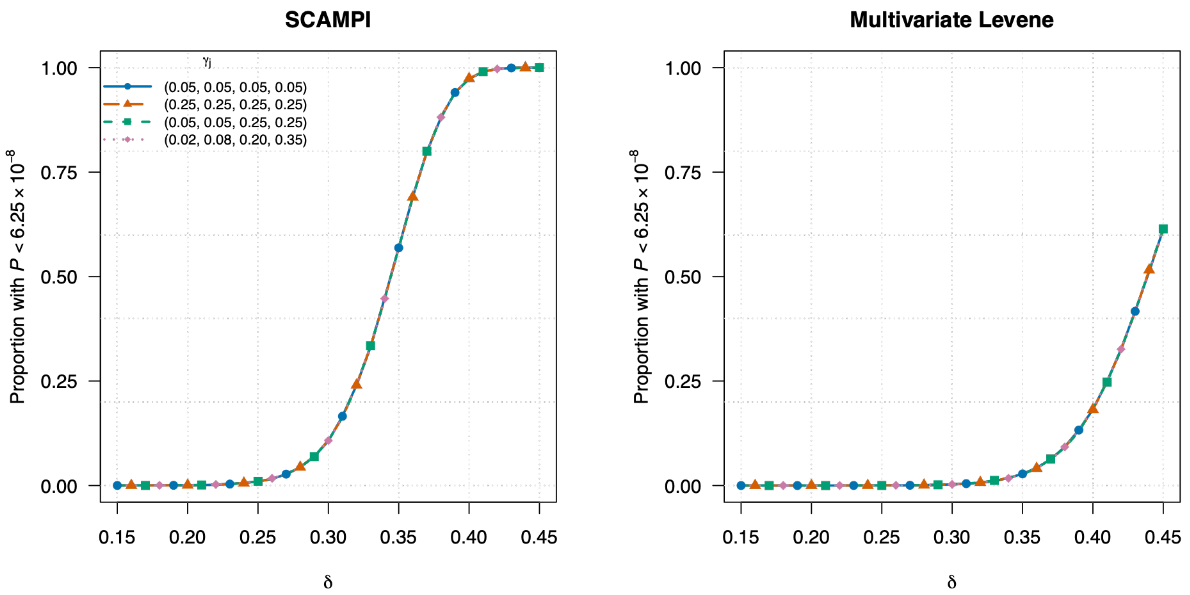


**Figure S11. Power under heterogeneous environmental main effects at the genome wide threshold**

Power is shown for SCAMPI and the multivariate Levene test using 20,000 individuals, four traits, MAF = 0.05, and the UK Biobank matched residual correlation matrix. The four environmental main effect vectors were (0.05, 0.05, 0.05, 0.05), (0.25, 0.25, 0.25, 0.25), (0.05, 0.05, 0.25, 0.25), and (0.02, 0.08, 0.20, 0.35). All four traits carried an interaction. The x axis shows the interaction coefficient, from 0.15 to 0.45 in increments of 0.01; the y axis shows the proportion of 100,000 alternative replicates with an omnibus P-value below $6.25 \times 10^{-8}$. The environmental variable was adjusted for in both methods. Colors and line types distinguish the four patterns; markers are shown at alternating simulated effect sizes to make overlapping curves visible, without shifting the plotted estimates. Power is evaluated at the nominal threshold, with null calibration results reported separately in Tables S3 and S8A.

**Figure S12.**

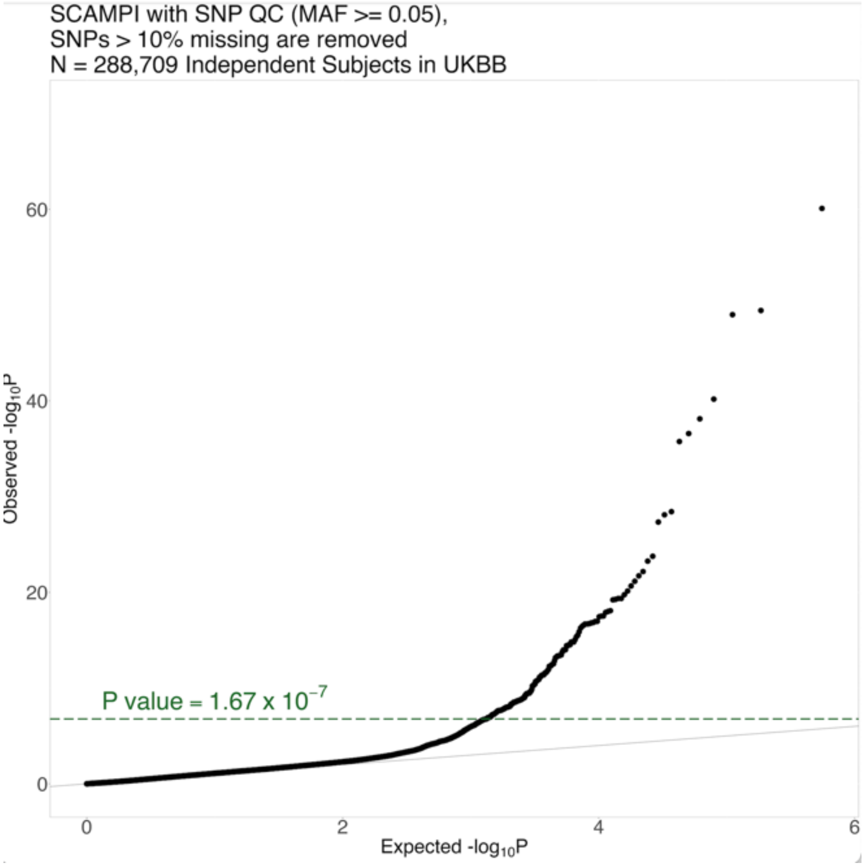


**Figure S12. Q-Q plot of the genome-wide results on lipid traits in UKBB using SCAMPI**

The x-axis shows the expected $-\log_{10}P$ under the global null hypothesis, and the y-axis shows the observed $-\log_{10}P$ for all SNPs analyzed by SCAMPI. The gray diagonal line represents equality between expected and observed values. The green dashed horizontal line marks the study-wide significance threshold $P$-value$=1.67 \times 10^{-7}$. The upward deviation in the extreme tail reflects the 210 genome-wide significant signals shown in Figure 3.

(a)

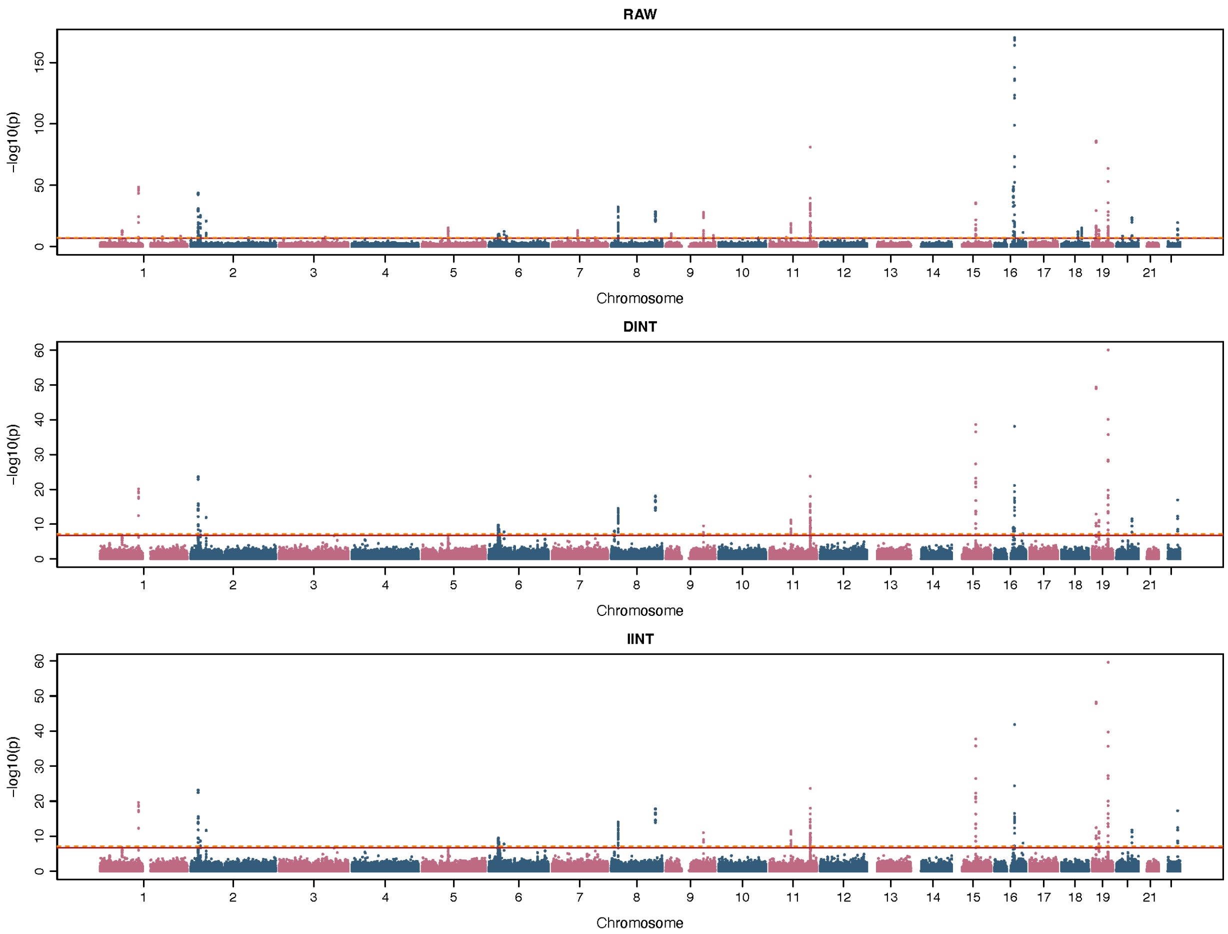


(b)

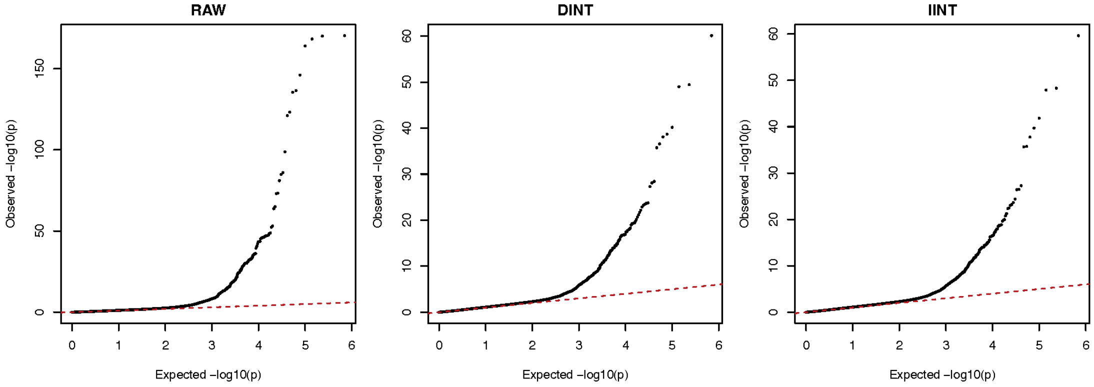


**Figure S13. Matched UK Biobank SCAMPI results across three phenotype scales**

(A) Manhattan plots for the pre-INT, direct INT, and indirect INT analyses. The x axis shows genomic position ordered by chromosome, and the y axis shows −log10(*P*). Horizontal lines mark $P = 1.67 \times 10^{-7}$ and $P = 6.25 \times 10^{-8}$.

(B) Quantile-quantile plots for the same analyses. The x axis shows expected −log10(P) under the global null, and the y axis shows observed −log10(*P*). All three analyses used the same 288,709 participants and 351,201 variants. The pre-INT scale used natural-log transformed triglycerides and original LDL cholesterol, HDL cholesterol, and BMI values.

(a)

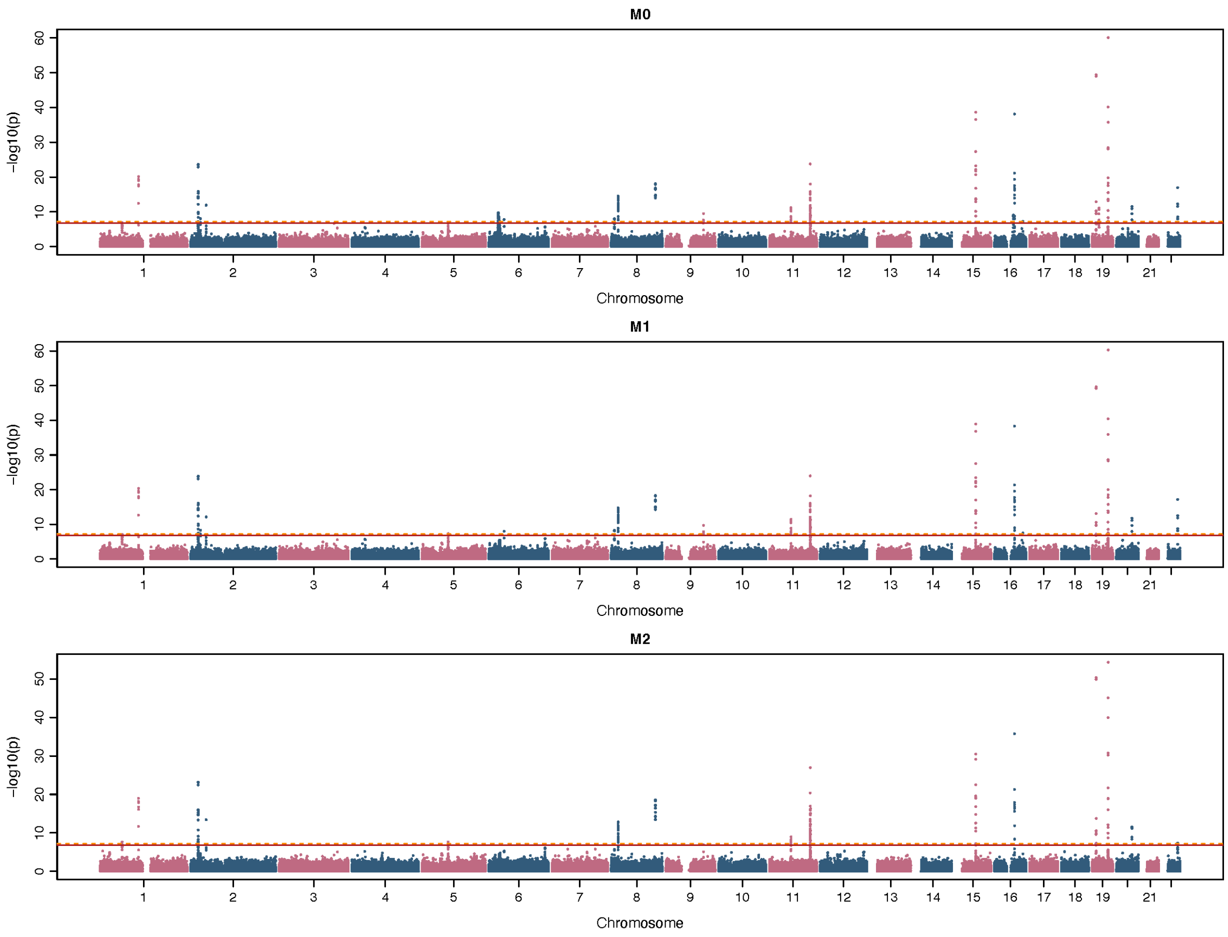


(b)

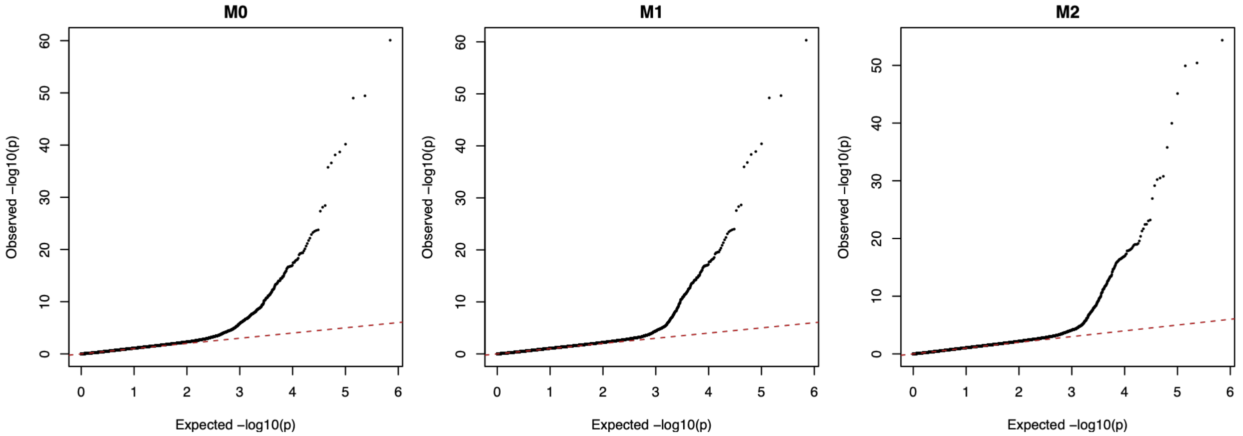


**Figure S14. UK Biobank comparison of the original four trait model and BMI sensitivity models.**

(A) Manhattan plots for M0, the original analysis of BMI and three lipid traits; M1, the three lipid traits without BMI; and M2, the three lipid traits adjusted for raw standardized BMI and BMI squared. The horizontal lines indicate $P$-value = $1.67 \times 10^{-7}$ and $P$-value = $6.25 \times 10^{-8}$.
(B) Corresponding quantile-quantile plots. All models used the same 288,709 participants and 351,201 matched variants. Genomic inflation factors were 0.940, 0.958, and 0.965 for M0, M1, and M2 respectively.

(a)

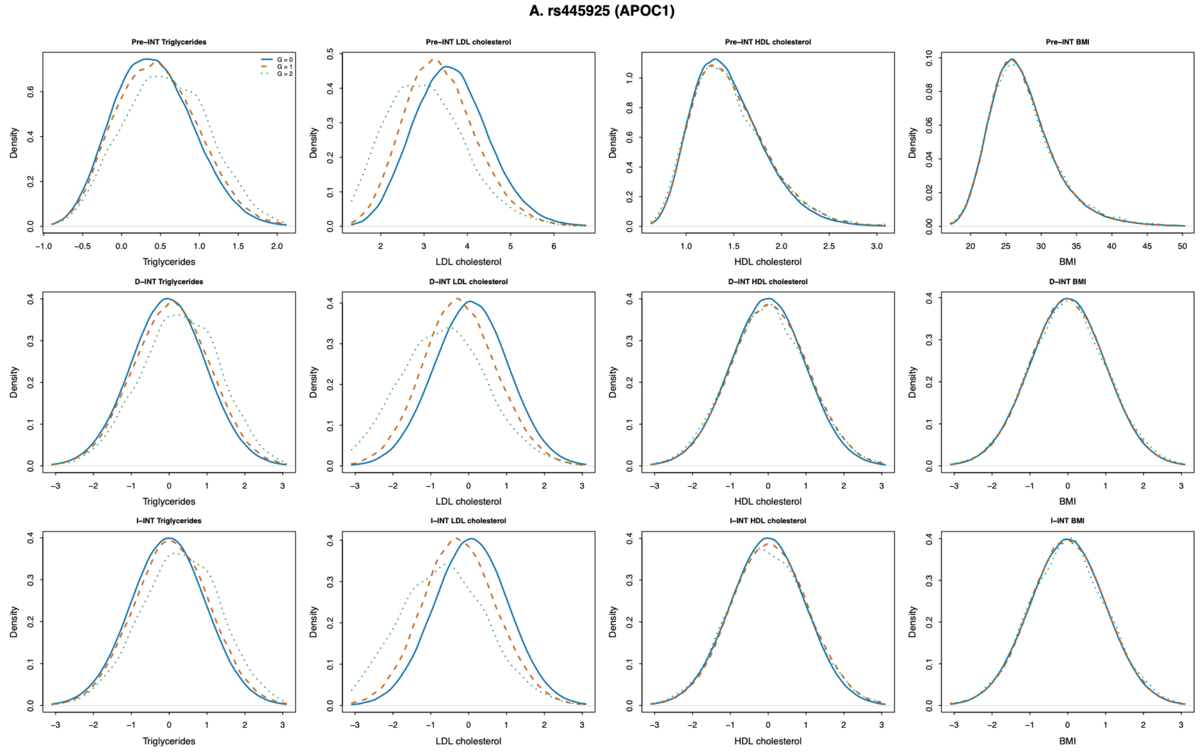
A. rs445925 (APOC1)
Pre–INT Triglycerides
Pre–INT LDL cholesterol
Pre–INT HDL cholesterol
Pre–INT BMI
D–INT Triglycerides
D–INT LDL cholesterol
D–INT HDL cholesterol
D–INT BMI
I–INT Triglycerides
I–INT LDL cholesterol
I–INT HDL cholesterol
I–INT BMI
G = 0
G = 1
G = 2
Density
Triglycerides
LDL cholesterol
HDL cholesterol
BMI

(b)

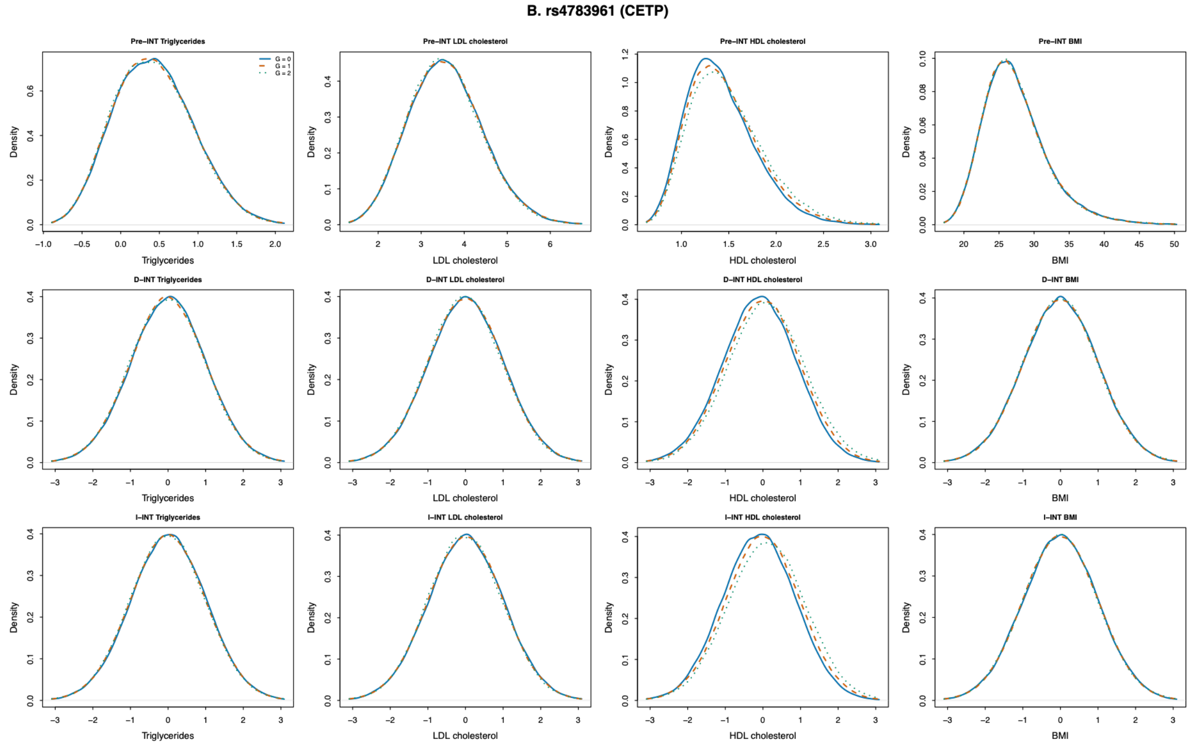
B. rs4783961 (CETP)
Pre–INT Triglycerides
Pre–INT LDL cholesterol
Pre–INT HDL cholesterol
Pre–INT BMI
D–INT Triglycerides
D–INT LDL cholesterol
D–INT HDL cholesterol
D–INT BMI
I–INT Triglycerides
I–INT LDL cholesterol
I–INT HDL cholesterol
I–INT BMI
G = 0
G = 1
G = 2
Density
Triglycerides
LDL cholesterol
HDL cholesterol
BMI

(c)

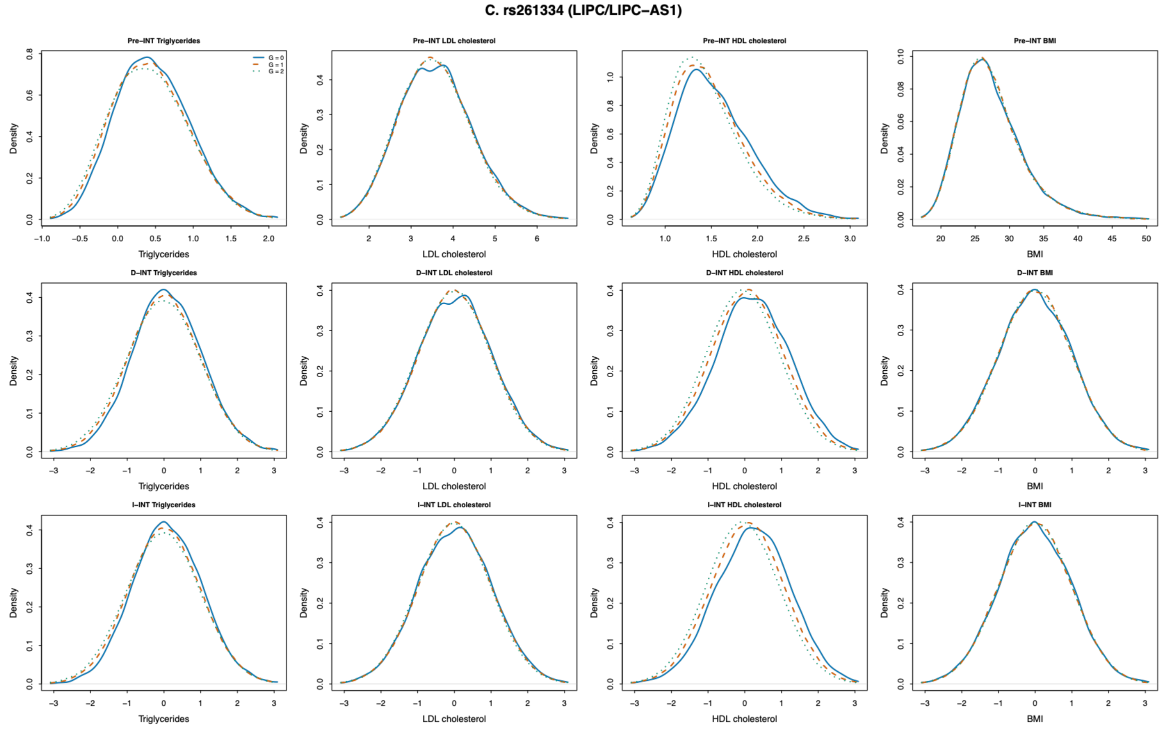
C. rs261334 (LIPC/LIPC−AS1)
Pre−INT Triglycerides
Pre−INT LDL cholesterol
Pre−INT HDL cholesterol
Pre−INT BMI
D−INT Triglycerides
D−INT LDL cholesterol
D−INT HDL cholesterol
D−INT BMI
I−INT Triglycerides
I−INT LDL cholesterol
I−INT HDL cholesterol
I−INT BMI
G = 0
G = 1
G = 2
Density
Triglycerides
LDL cholesterol
HDL cholesterol
BMI

(d)

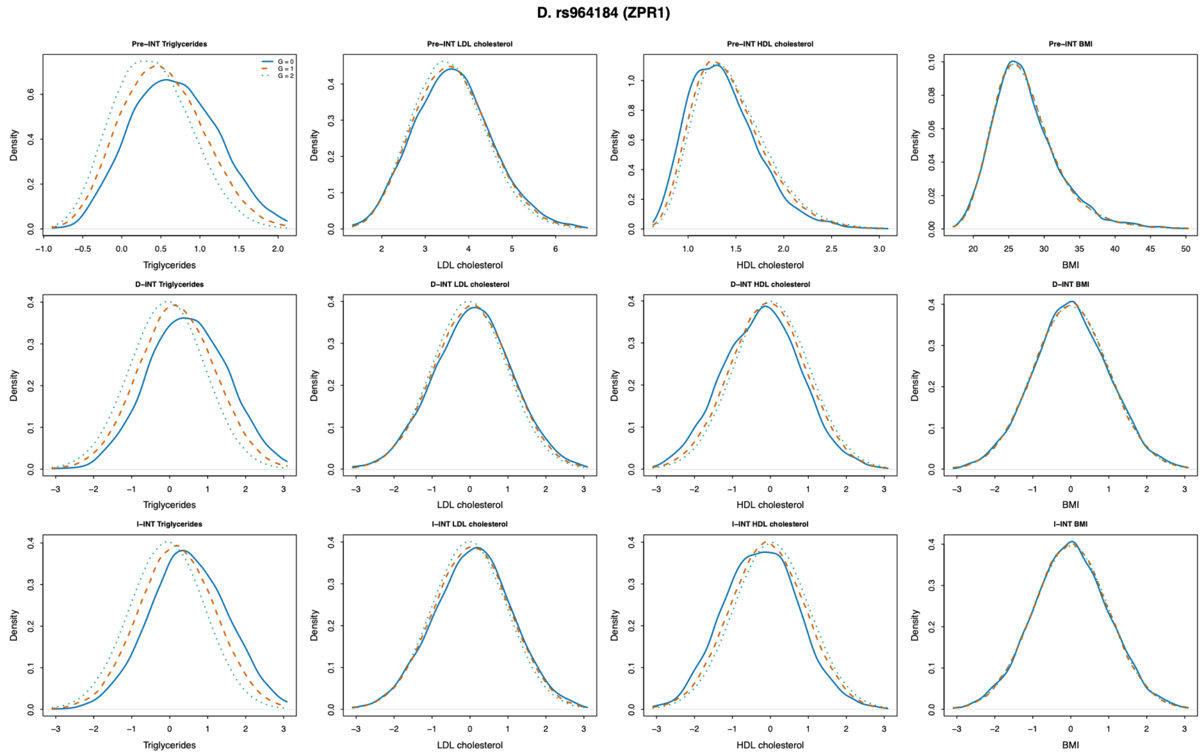
D. rs964184 (ZPR1)
Pre−INT Triglycerides
Pre−INT LDL cholesterol
Pre−INT HDL cholesterol
Pre−INT BMI
D−INT Triglycerides
D−INT LDL cholesterol
D−INT HDL cholesterol
D−INT BMI
I−INT Triglycerides
I−INT LDL cholesterol
I−INT HDL cholesterol
I−INT BMI
G = 0
G = 1
G = 2
Density
Triglycerides
LDL cholesterol
HDL cholesterol
BMI

(e).

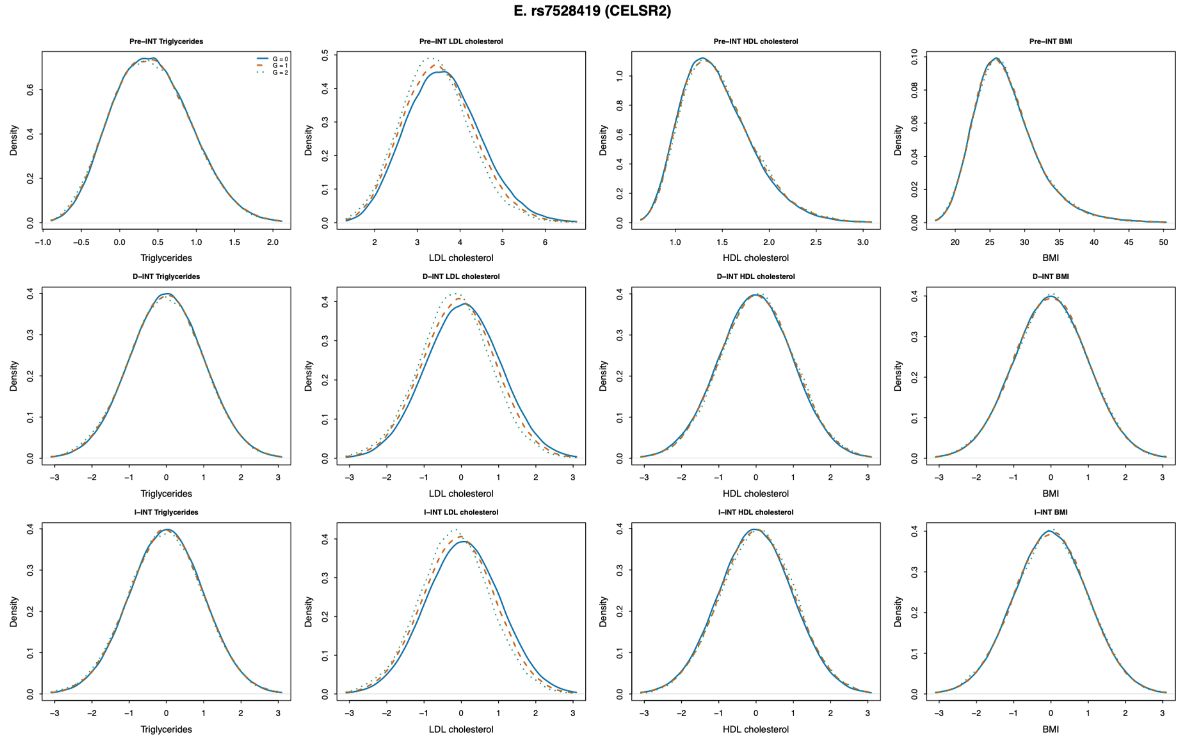


**Figure S15. Genotype-stratified phenotype distributions for the five leading SCAMPI variants across three analysis scales**

Density plots are shown for triglycerides, LDL cholesterol, HDL cholesterol, and BMI within genotype categories 0, 1, and 2 for rs445925, rs4783961, rs261334, rs964184, and rs7528419. Results are displayed on the pre-INT, direct INT, and indirect INT scales. The pre-INT triglyceride phenotype was natural-log transformed. Panels (a)-(e) correspond to rs445925, rs4783961, rs261334, rs964184, and rs7528419, respectively. Within each panel, rows show the pre-INT, direct INT, and indirect INT scales, and columns show triglycerides, LDL cholesterol, HDL cholesterol, and BMI. Colors identify genotype categories. These plots are descriptive and illustrate the scale dependence of genotype-specific phenotype distributions.

(a)

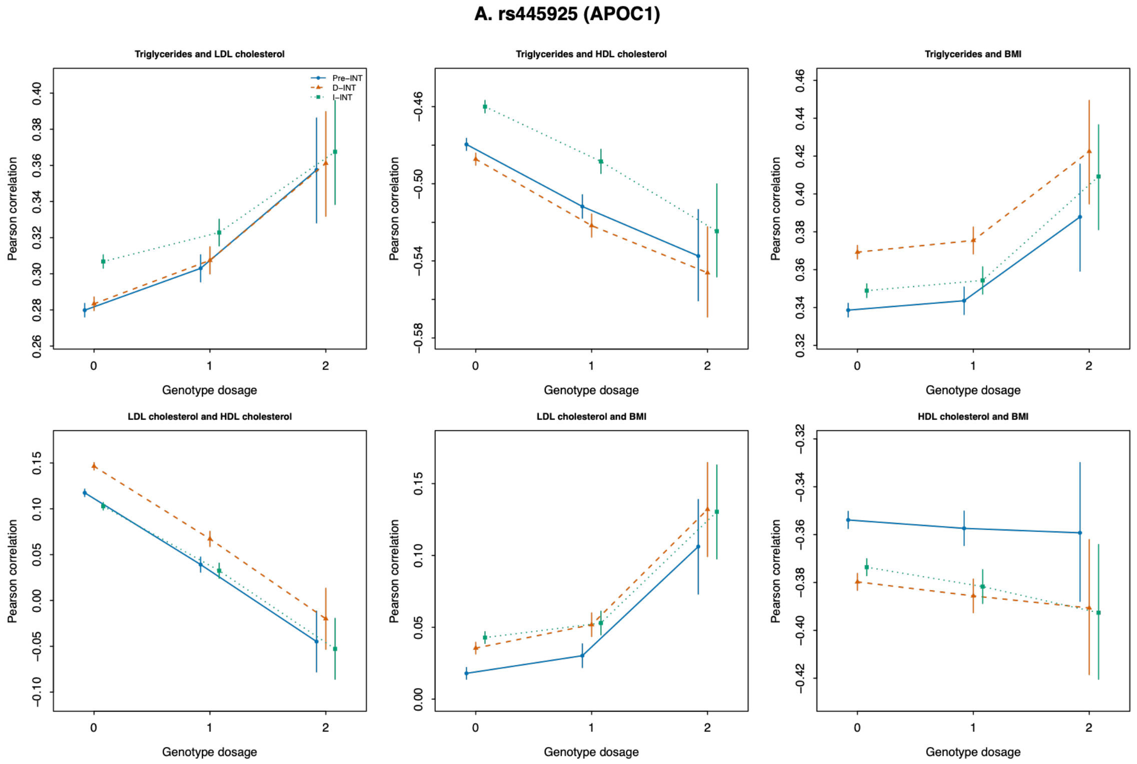
A. rs445925 (APOC1)
Triglycerides and LDL cholesterol
Triglycerides and HDL cholesterol
Triglycerides and BMI
LDL cholesterol and HDL cholesterol
LDL cholesterol and BMI
HDL cholesterol and BMI
Pre–INT
D–INT
I–INT
Pearson correlation
Genotype dosage

(b)

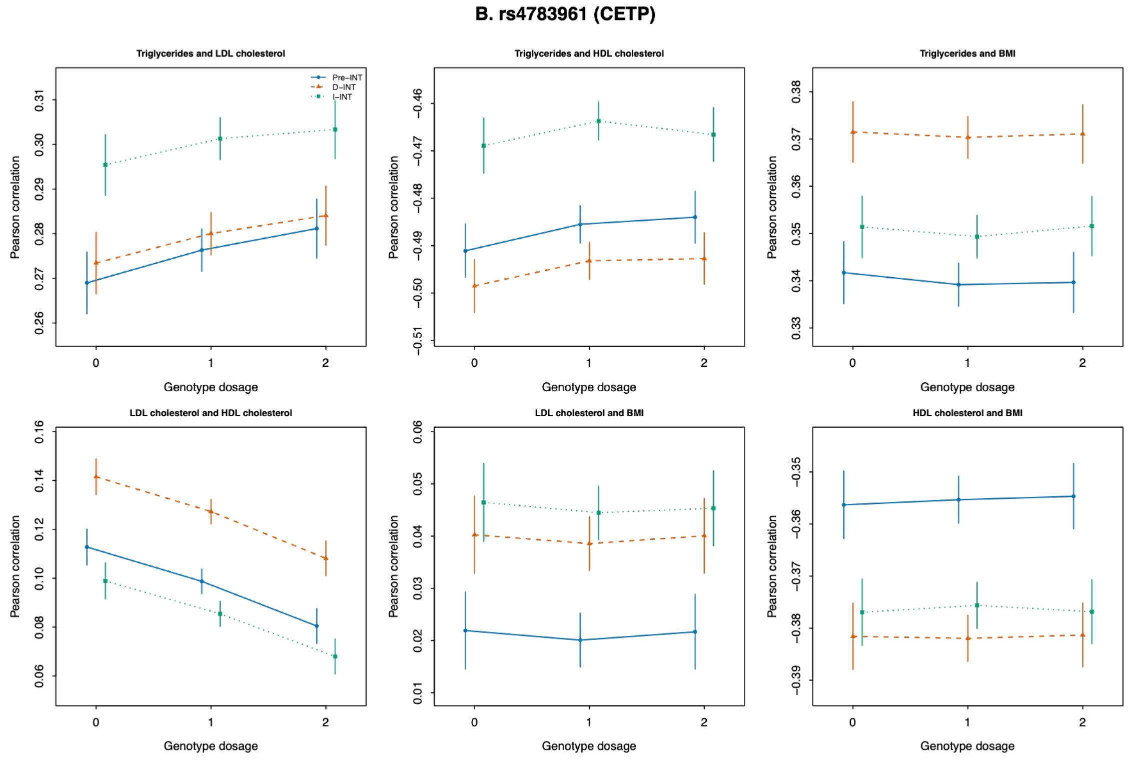
B. rs4783961 (CETP)
Triglycerides and LDL cholesterol
Triglycerides and HDL cholesterol
Triglycerides and BMI
LDL cholesterol and HDL cholesterol
LDL cholesterol and BMI
HDL cholesterol and BMI
Pre–INT
D–INT
I–INT
Pearson correlation
Genotype dosage

(c)

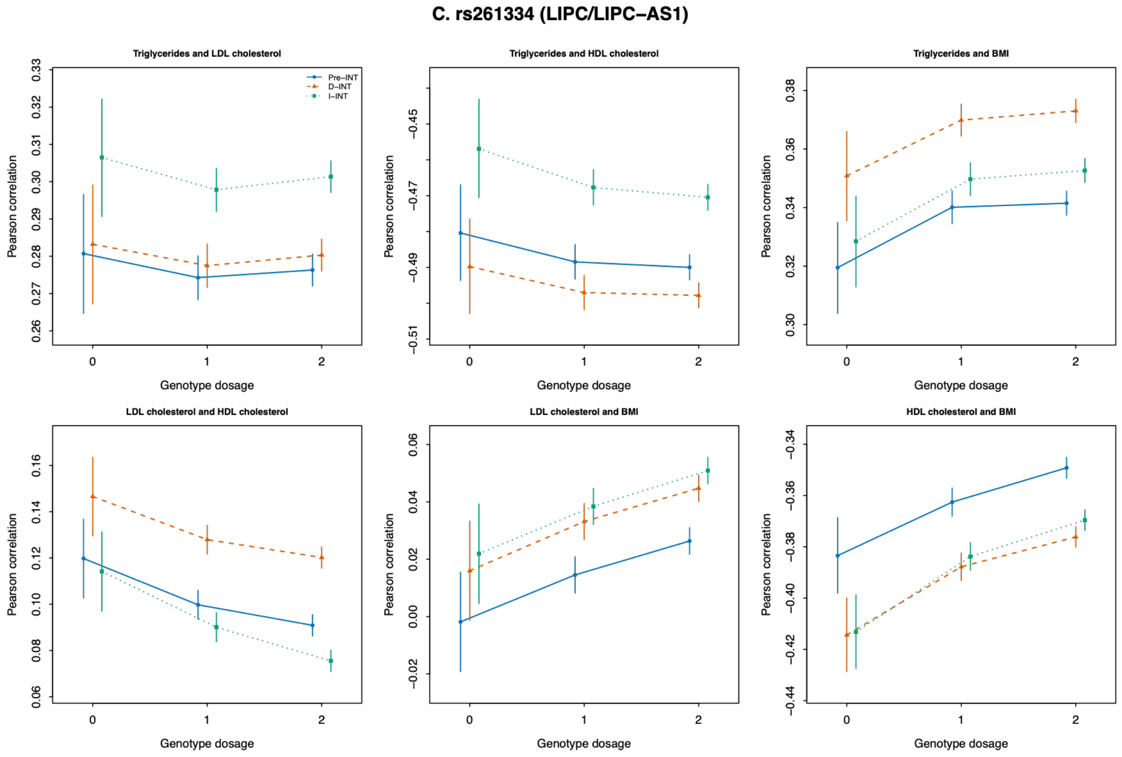

C. rs261334 (LIPC/LIPC−AS1)
Triglycerides and LDL cholesterol
Triglycerides and HDL cholesterol
Triglycerides and BMI
LDL cholesterol and HDL cholesterol
LDL cholesterol and BMI
HDL cholesterol and BMI
Pre−INT
D−INT
I−INT
Pearson correlation
Genotype dosage


(d)

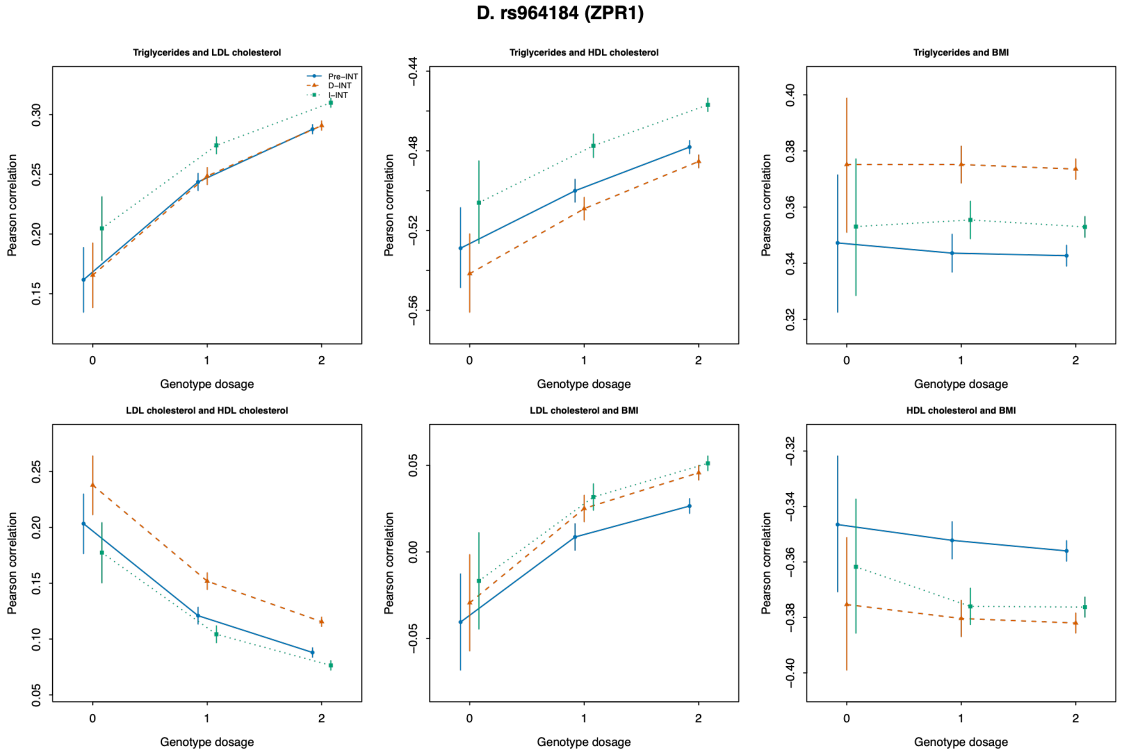

D. rs964184 (ZPR1)
Triglycerides and LDL cholesterol
Triglycerides and HDL cholesterol
Triglycerides and BMI
LDL cholesterol and HDL cholesterol
LDL cholesterol and BMI
HDL cholesterol and BMI
Pre−INT
D−INT
I−INT
Pearson correlation
Genotype dosage

(e)

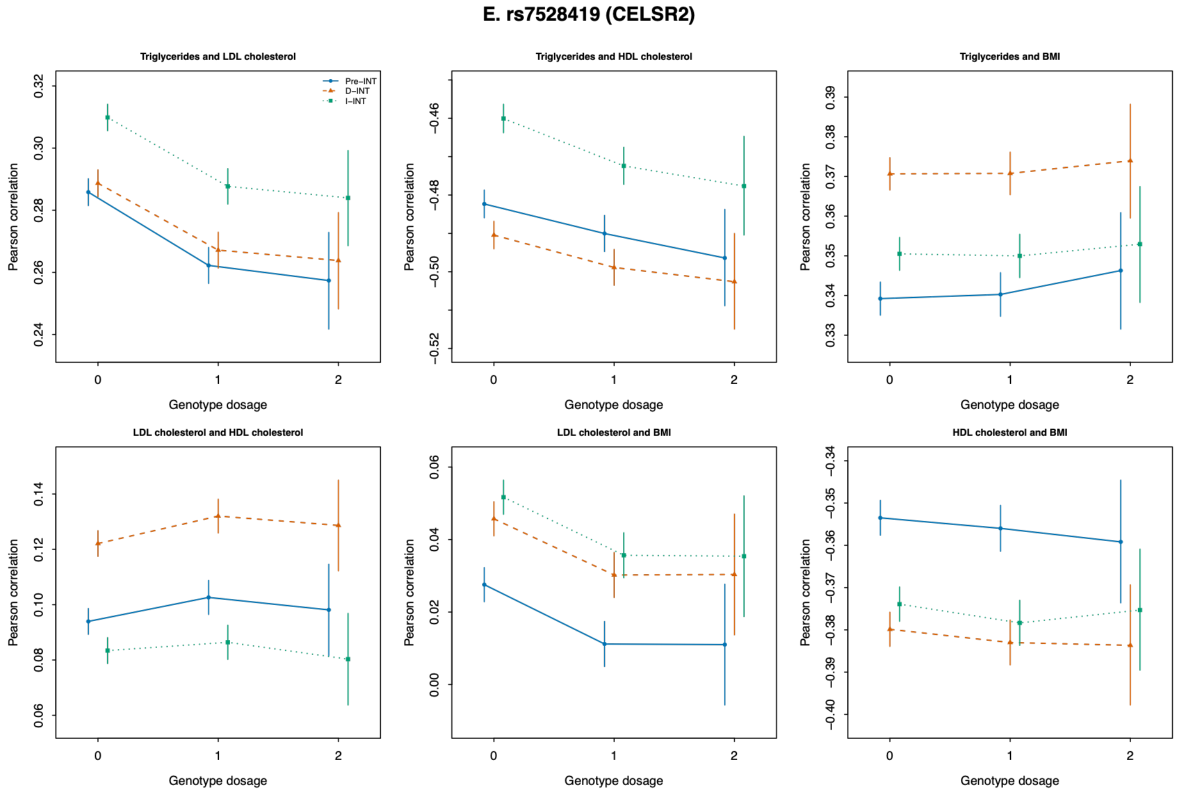


**Figure S16. Genotype-specific pairwise trait correlations for the five leading SCAMPI variants across three analysis scales**

Points show Pearson correlation estimates between each pair of triglycerides, LDL cholesterol, HDL cholesterol, and BMI within genotype categories 0, 1, and 2. Error bars show 95% confidence intervals based on Fisher's z transformation. Separate estimates are displayed for the pre-INT, direct INT, and indirect INT scales. Panels correspond to rs445925, rs4783961, rs261334, rs964184, and rs7528419. These diagnostics show the genotype specific covariance patterns contributing to the SCAMPI signals.

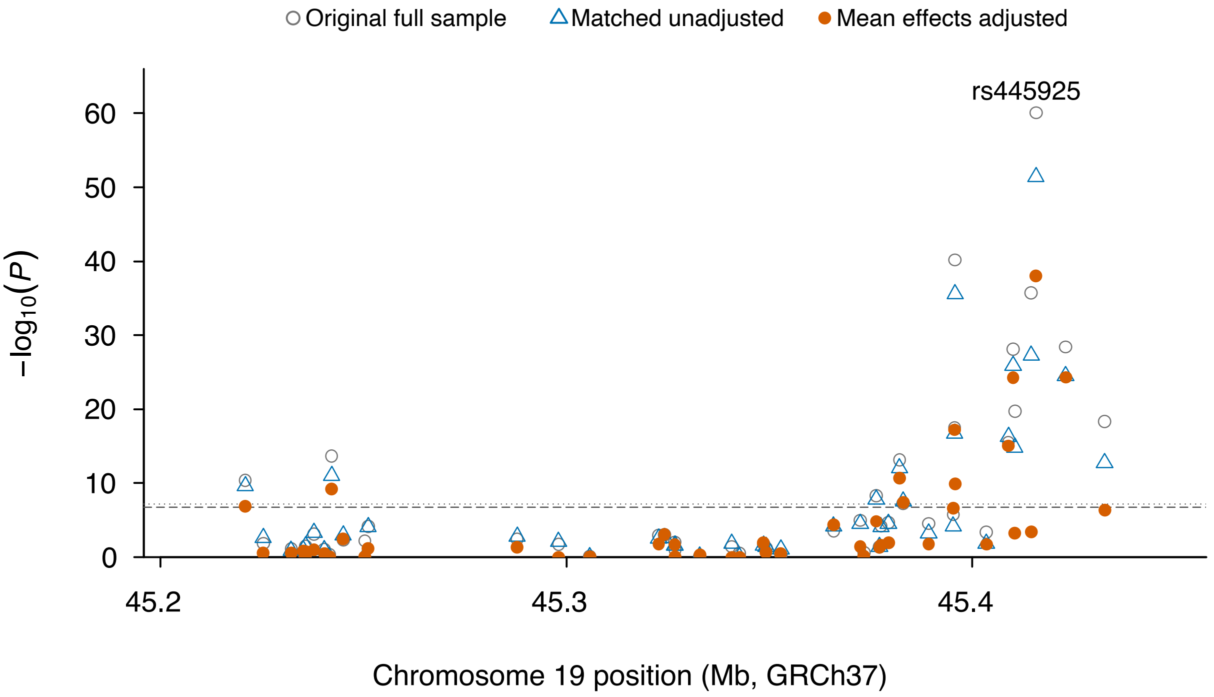


**Figure S17. Regional comparison of APOE/APOC1 association P-values before and after adjustment for the joint unphased rs429358/rs7412 genotype structure**

The x axis shows chromosome 19 position (GRCh37), and the y axis shows $-\log_{10}(P)$. Results for 47 variants are shown for the original full sample analysis, the matched subset without joint genotype adjustment, and the same subset with five indicators for the six common joint unphased rs429358/rs7412 genotype categories included in the DGLM mean model. The dashed and dotted horizontal lines indicate $P = 1.67 \times 10^{-7}$ and $P = 6.25 \times 10^{-8}$, respectively. Fourteen variants were significant in both unadjusted analyses at both thresholds; after mean adjustment, ten and nine remained significant at the respective thresholds.

**Figure S18.**

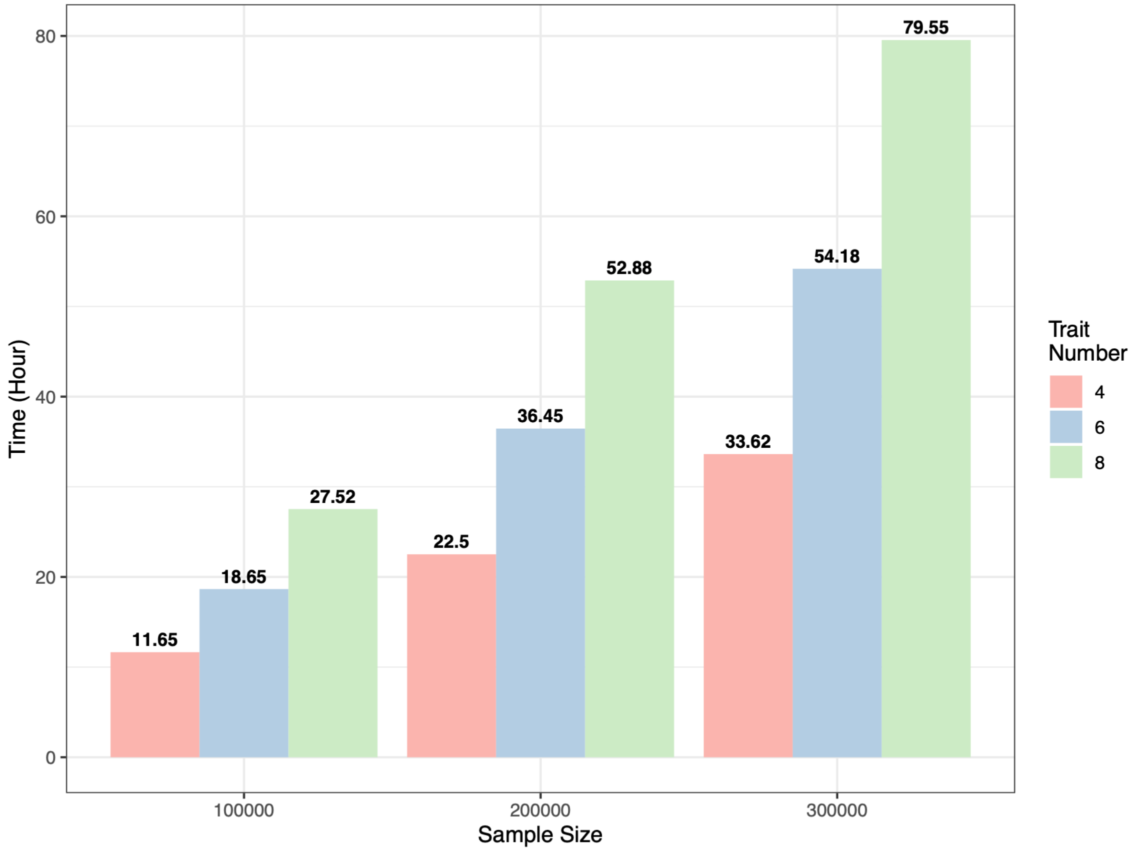


**Figure S18. Computational performance of SCAMPI for analyzing 6,000,000 SNPs**

The x axis shows sample size, with N = 100,000, 200,000, or 300,000, and bar colors represent analyses of 4, 6, or 8 traits. The left y axis shows the mean computation time in seconds for analyzing one SNP, estimated from 1,000 independent benchmark repetitions on the RSPH HPC system. The right y axis shows the corresponding projected wall clock time for analyzing 6,000,000 SNPs using 1,000 concurrent workers. Genome-wide times were obtained by linear extrapolation from the measured per SNP runtimes and are projections rather than observed end to end runtimes. Actual wall clock time will depend on available resources, scheduler policy, workload balance, data input and output, and other system overhead.

**Figure S19.**

**(a)**

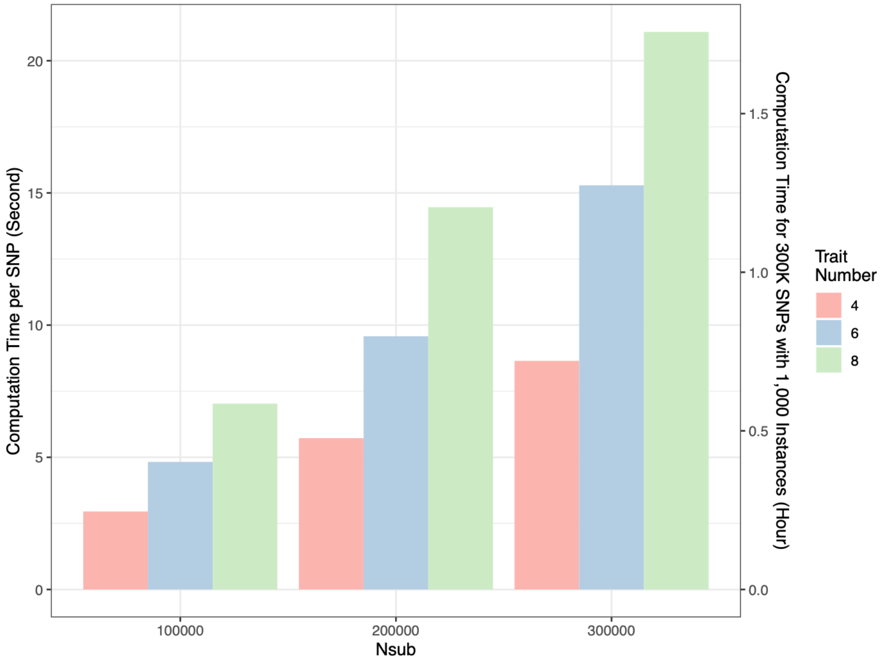


**(b)**

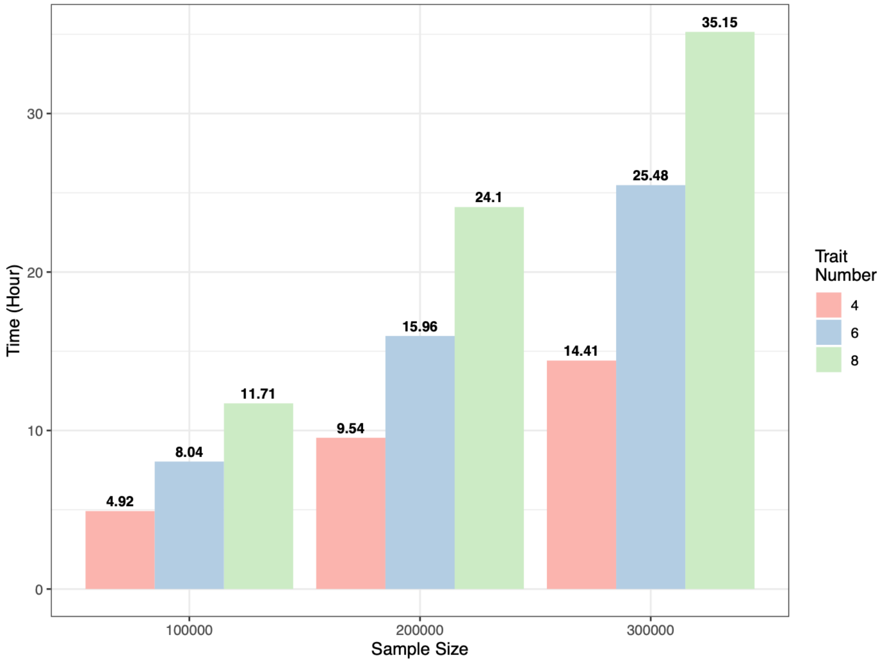

**Figure S19. Computational performance of SCAMPI estimated using Apple M1 chip**

The x axis shows sample size, with N = 100,000, 200,000, or 300,000, and bar colors represent analyses of 4, 6, or 8 traits. The left y axis shows the mean computation time in seconds for analyzing one SNP, estimated from 1,000 independent benchmark repetitions. The right y axis shows projected wall clock time using 1,000 concurrent workers. Panel (a) shows the projection for 300,000 SNPs, and panel (b) shows the projection for 6,000,000 SNPs. Genome-wide times were obtained by linear extrapolation from the measured per SNP runtimes and are projections rather than observed end to end runtimes.
